# Delphy: scalable, near-real-time Bayesian phylogenetics for outbreaks

**DOI:** 10.1101/2025.03.25.645253

**Authors:** Patrick Varilly, Mark Schifferli, Katherine Yang, Tim Burcham, Paul Cronan, Olivia Glennon, Olivia Jacks, Ellory Laning, Libby Marrs, Kyle Oba, Shannon Yeung, Edyth Parker, Ifeanyi Omah, Jonathan E. Pekar, Laura Luebbert, Kristian G. Andersen, Daniel J. Park, Stephen F. Schaffner, Bronwyn L. MacInnis, Christian Happi, Jacob E. Lemieux, Al Ozonoff, Michael D. Mitzenmacher, Ben Fry, Pardis C. Sabeti

**Affiliations:** Broad Institute of Harvard and MIT, 415 Main Street, Cambridge, MA 02142, USA; Fathom Information Design, Boston, MA 02114, USA; The Institute of Genomics and Global Health, Redeemer’s University, Ede, Osun State, Nigeria; Institute of Ecology and Evolution, University of Edinburgh, Edinburgh EH9 3FL, UK; Department of Parasitology and Entomology, Nnamdi Azikiwe University, Awka, Anambra state Nigeria; Department of Immunology and Microbiology, The Scripps Research Institute, La Jolla, CA 92037, USA; Scripps Research Translational Institute, La Jolla, CA 92037, USA; Department of Immunology and Infectious Diseases, Harvard T.H. Chan School of Public Health, Harvard University, Boston, MA 02115, USA; Massachusetts Consortium on Pathogen Readiness, Harvard Medical School, Harvard University, Boston, MA 02115, USA; Department of Biological Sciences, Faculty of Natural Sciences, Redeemer’s University, Ede, Osun State, Nigeria; Department of Medicine, Massachusetts General Hospital, Harvard Medical School, Boston, Massachusetts, USA; Department of Computer Science, School of Engineering and Applied Sciences, Harvard University, Boston, MA 02134, USA; Department of Organismic and Evolutionary Biology, Faculty of Arts and Sciences, Harvard University, Cambridge, MA 02138, USA; Howard Hughes Medical Institute, Chevy Chase, MD 20815, USA

## Abstract

Pathogen genomic analysis is central to tracking, understanding, and containing outbreaks, but complexity and high costs of state-of-the-art (SOTA) phylogenetic tools limit global access and impact. We introduce Delphy, an exact reformulation of Bayesian phylogenetics designed to transform its speed, scalability and accessibility while retaining SOTA accuracy. Delphy’s central data structure, an Explicit Mutation Annotated Tree, exploits the high sequence similarity in large-scale epidemic datasets for efficient tree exploration and convergence. By reproducing key analyses from recent major epidemics (Ebola, Zika, SARS-CoV-2, mpox, and H5N1), we demonstrate SOTA accuracy with up to 1,000x speedups. Assessing Delphy’s scalability, we show that a simulated dataset of 100,000 sequences can be analyzed in under a day–the largest such computation to date. We distribute Delphy as a client-side web application, enabling users worldwide to turn raw data into interactive results within minutes, without the data ever leaving the user’s machine. Delphy automatically identifies key viral lineages and mutations, as well as their emergence and prevalence through time, all with quantified uncertainties derived from a solid theoretical foundation. Delphy shows the power of Bayesian phylogenetics as a fast, accessible frontline tool for tackling future outbreaks.

## Introduction

Large-scale sequencing of pathogen genomes, now readily achievable thanks to next-generation sequencing, has the potential to provide unprecedented clarity in our understanding of pathogen spread and evolution during disease outbreaks. However, the resulting massive datasets–illustrated by the 16+ million publicly available SARS-CoV-2 genomes^1,2^–pose an equally unprecedented challenge for those seeking to analyze and interpret them.

Phylogenetics is the fundamental tool for organizing and interpreting pathogen genome sequences, inferring plausible trees of descent that explain how observed pathogens arose. Phylogenetic reconstruction enables a wide range of analysis, including: (a) dating the start of outbreaks^3–5^; (b) distinguishing zoonotic spillovers from human-to-human transmission^3,6^; (c) identifying major lineages^7^; (d) detecting early warning signs of concerning new lineages^8^; (e) revealing community and geographic-based spread patterns^6,9–11^; and (f) reconstructing detailed transmission networks^12–15^.

Among phylogenetic approaches, Bayesian phylogenetics, as implemented in widely used tools like BEAST^16^, BEAST2^17^ and Mr. Bayes^17,18^, is the gold standard. Phylogenetic tree reconstruction is inherently uncertain and Bayesian methods fully embrace and quantify this uncertainty, while simultaneously inferring important covariates, such as geographic spread^6^ or changes in viral population size through time^6,19^.

Despite its advantages, Bayesian phylogenetics is in current practice technically complex. A handful of specialized research groups can handle this complexity, but it remains inaccessible to most epidemiologists and public health bodies. These Bayesian techniques are also computationally expensive, so much so that, in current practice, outbreak-tracking infrastructures (e.g., NextStrain^20^, Cov2Tree^21^) rely on more approximate methods, such as maximum-parsimony^22,23^ or maximum-likelihood^24–26^. These methods typically generate a single tree topology, which is then post-processed using ad-hoc techniques to infer uncertainties and covariates^12,20,27,28^.

When applying Bayesian phylogenetics to sequences that are nearly identical, it is beneficial to reformulate the calculations in terms of local mutations instead of all mutated sites in a dataset. Such an exact reformulation is always possible, but should yield especially large efficiencies when mutations are sparse. Outbreak sequences often differ from their closest related sequence at as few as 0 to 2 sites^23,29,30^, and so clearly exhibit such sparsity.

Recent work exploiting the low diversity of outbreak data sequences has dramatically improved the performance and scalability of maximum-parsimony^22,23^ and maximum-likelihood methods^25,26^, suggesting that similar gains are indeed possible in a Bayesian framework.

However, previous attempts were limited in significant ways, including fixed tree topology, simplified evolution models, inability to handle missing data, no site rate heterogeneity, and absence of ancestry prior^31–39^. These efforts also missed key opportunities for computational efficiencies, such as avoiding operations in inner loops that scale with genome size and leveraging parallelism. To date, no Bayesian phylogenetics tool using an explicit mutation representation can simultaneously address all the complexities of real-world genomic epidemiology datasets needed to be broadly useful in this context.

In this work, we present Delphy, the result of reimagining and rebuilding every portion of the Bayesian phylogenetics pipeline to make it fast, accurate, accessible, and scalable for large-scale outbreak data. Our key goals are to: (a) enable any frontline worker to analyze their own data using SOTA methods, without specialized training or computational resources exceeding those of a standard laptop; (b) allow them to generate publication-quality figures from raw aligned sequences within an hour, making Bayesian phylogenetics a standard part of the real-time public health responses worldwide; and (c) to ensure efficient scaling to accommodate the growing volume of genomic data, without sacrificing accuracy.

### Delphy implements Bayesian Phylogenetics using Explicit Mutation-Annotated Trees (EMATs)

Delphy’s computations are organized around Explicit Mutation-Annotated Trees (EMAT), which are concrete hypotheses of how an ancestral sequence replicated and evolved, mutation by mutation, into the observed dated sequences (Figure 1A). An EMAT is a timed bifurcating tree: tips correspond to observed sequences; inner nodes correspond to the most-recent common ancestor (MRCA) of their descendent tips; and branches trace out lineages of successive sequences. Every point on the tree thus represents a sequence that existed at a specific time and contributed to the ancestry of an observed sequence. EMATs represent the root sequence explicitly and all other sequences implicitly, in terms of accumulated mutations from the root.

**Figure 1:**
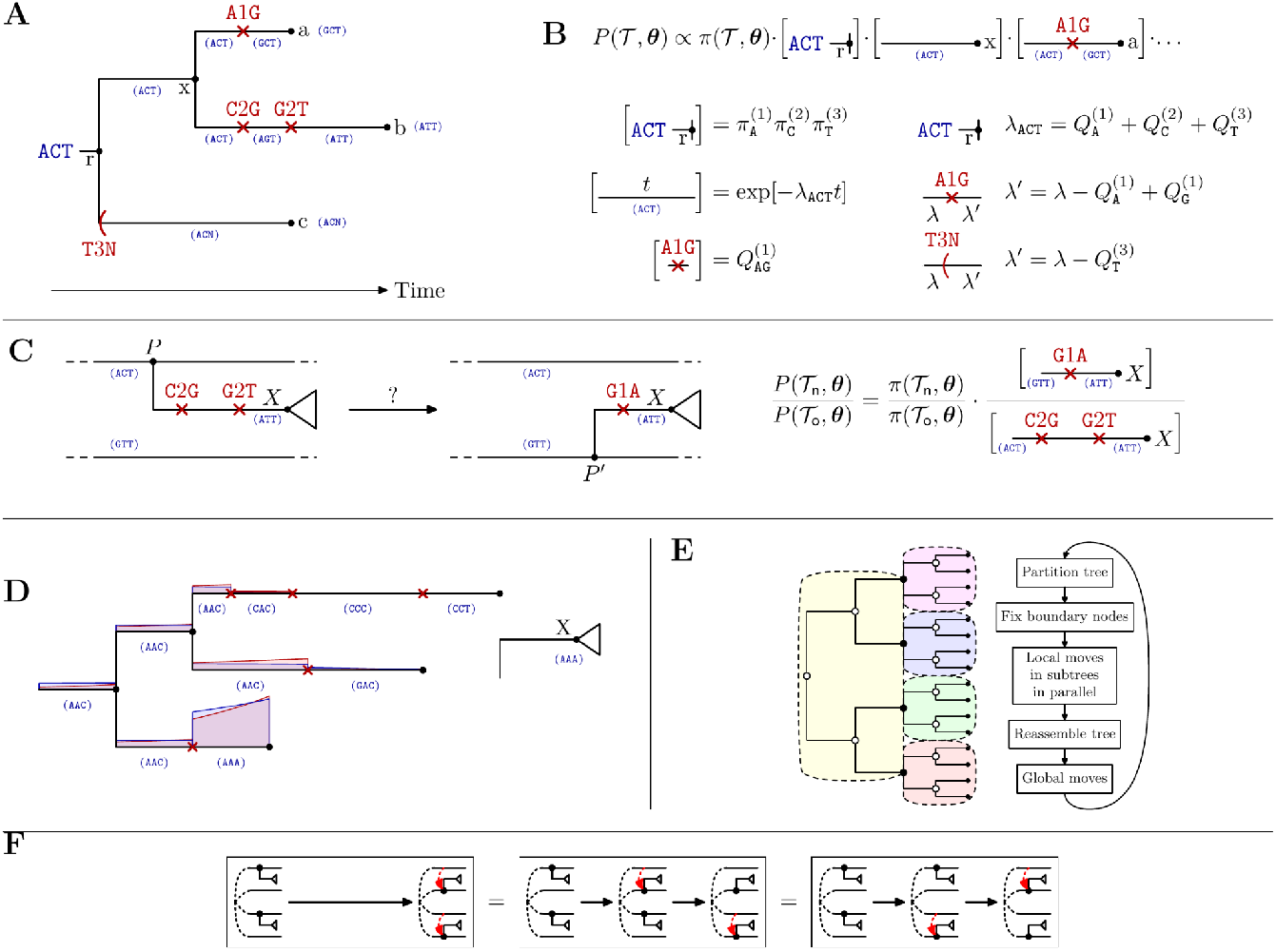
Key elements of Delphy. **A**. Phylogenetic trees are represented by explicit mutation-annotated trees (EMATs), timed bifurcating trees with an explicit root sequence and explicit mutation and “missation” events on branches at explicit times; every point has a specific but implicitly encoded sequence (in parentheses). **B**. The posterior distribution *P(T*,*θ)* sampled by Delphy is dominated by a *genetic prior*, a product of three kinds of simple terms that mirror the tree structure; the value of the genetic prior can be calculated with the simple rules shown, in time proportional to the number of elements in the tree (see text and Methods for more detail). **C**. Schematic of a subtree pruning and regrafting (SPR) move. When pruning a subtree rooted at X from P and deciding at what point P’ to regraft it, we fix everything except the branch above X, so the ratio of posteriors of the new tree *T*_*n*_ and old tree *T*_*o*_ is dominated by a simple ratio of small numbers of factors, dependent only on what happens along that branch. **D**. Schematic of a mutation-directed SPR (mdSPR) move: when evaluating possible regrafting locations for the subtree rooted at X, the EMAT tree structure facilitates proposals according to a simple Jukes-Cantor model (red) that closely tracks the real conditional posterior (blue); hence, plausible large-scale rearrangements are proposed and accepted with high probability. **E**. Schematic of scaling strategy: the proposal and acceptance of local tree rearrangements in an EMAT is independent of what happens in distant portions of the tree; a tree can thus be partitioned into smaller subtrees, each subject to independent, parallel Markov-Chain Monte Carlo (MCMC) moves, and then reassembled to apply global moves. **F**. Moves can be parallelized because the probability of proceeding from an initial state to a particular final state in parallel is equal to that when moves are interleaved arbitrarily and attempted serially.

Branches contain explicit, time-stamped mutation events and may also contain *missation* events–EMAT-specific analogs to mutations, which mark points where data for specific sites is missing in all downstream observed sequences (see below). EMATs are inspired by UShER’s Mutation-Annotated Trees (MATs), extending them by: (a) assigning time-stamps to each node and each mutation; (b) allowing multiple mutations on the same site along a single branch; and (c) explicitly encoding missing data.

Given a set of dated observed sequences, Delphy samples plausible EMATs *T* that link them and associated parameters *θ* according to a *posterior distribution P(T*,*θ*). This distribution is proportional to two factors. The first, a *prior* distribution on (*T*,*θ*), represents the probability that randomly-drawn sequences are ancestrally related as in *T*, and evolve from a random starting point accordingly. The parameters *θ* include the mutation rate, the relative likelihood of different mutations, and the viral population size over time. The second factor, the *likelihood* of observing the sequences given the model (*T*,*θ*), is currently trivial because Delphy neglects sequencing error–it is 1 if the tip sequences in (*T*,*θ*) match the observations, and 0 otherwise.

The prior distribution is largely shaped by a *genetic prior*, which represents the probability that a randomly chosen root sequence evolves along *T* according to the explicit mutations therein (Figure 1B). Evolution is modelled independently at each site *ℓ* as a Continuous-Time Markov Chain with transition rate matrix Q^(*ℓ*)^_ab_, and escape rates Q^(*ℓ*)^_a_ = -Q^(*ℓ*)^_aa_ . The stationary distribution π^(*ℓ*)^_a_ of the chain at site *ℓ* serves as the prior distribution for the root sequence. In an EMAT, the genetic prior factors into simple terms: one for the root involving π^(*ℓ*)^_a_, one for each branch involving Q^(*ℓ*)^_a_, and one for each mutation involving Q^(*ℓ*)^_ab_ . Apart from the genetic prior, the remaining factors of the posterior, denoted by *π(T*,*θ*), relate to ancestry and parameter priors in standard ways (Methods and SI).

The genetic prior functions similarly to the Felsenstein tree likelihood in traditional Bayesian phylogenetics^16,17,40^, making EMAT sampling analogous to sampling tree topologies in that framework. For a fixed topology and tip sequences, integrating the genetic prior over all possible root sequences and mutation histories yields the Felsenstein tree likelihood (see SI). Whereas traditional methods effectively perform this integration analytically, Delphy performs it statistically. That is, Delphy simultaneously explores tree topologies and actual realizations of the root sequences and mutation histories on them, of which only a few are typically plausible given the observed data. In contrast, calculating the Felsenstein tree likelihood for a fixed topology accounts for all possible assignments of root sequences and mutation histories, but the overwhelming computational effort is spent on configurations that are vanishingly rare.

Delphy samples models (*T*,*θ*) from the posterior distribution *P*(*T*,*θ*) using a standard Markov-Chain Monte Carlo (MCMC) algorithm, whereby a sequence of (correlated) samples is generated by repeatedly proposing small changes, called *moves*, which are probabilistically accepted to achieve the correct sampling distribution^41,42^. The most common acceptance probability is given by the Metropolis-Hastings (MH) criterion, which involves the ratio of the posterior *P*(*T*,*θ*) before and after the proposed change^43,44^. Moves where most factors in this ratio cancel out can be proposed and evaluated efficiently.

Delphy’s central move, *subtree pruning and regrafting* (SPR, Figure 1C), is computationally efficient on EMATs. SPR moves propose detaching a subtree rooted at X from its attachment point P and reattaching it elsewhere at P’. In an EMAT, a new mutational history must also be proposed, which we currently restrict to changes along the P’-X branch, so that the posterior ratio in the MH criterion consists of the few factors arising from that branch. Since mutations on that branch are overwhelmingly dictated by the differences between sequences at the endpoints, we use a simplified Jukes-Cantor model for proposals and rely on the MH criterion to correct for approximation errors (see SI). When handling missing data, missations on the path from P to P’ (see below) introduce additional but manageable complexity (see SI).

Delphy is much faster than traditional Bayesian phylogenetics tools because it does far less work: evaluating moves on EMATs with a genetic prior involves orders of magnitude fewer operations than evaluating analogous moves on tree topologies with the Felsenstein tree likelihood. Concretely, calculating ratios of Felsenstein tree likelihoods may involve hundreds of thousands of operations for realistic datasets, since partial site likelihoods must be recalculated for every variable site across the entire dataset and for every node between the two attachment points and the root. In contrast, posterior ratios for moves on EMATs require only a handful of local operations, proportional to the number of affected mutations. The tradeoff is that EMAT-based moves are less statistically efficient than those on tree topologies. While the number of moves between uncorrelated samples in the MCMC is larger, this ratio is typically substantially smaller than the ratio of costs per move.

The local nature of most moves in Delphy opens the door to analysis of pandemic-scale datasets: the genetic prior, unlike the Felsenstein tree likelihood, factorizes over subtrees, so Delphy can parallelize moves in different regions of the tree (Figures 1E and 1F). For example, Figure 1C shows how the genetic prior ratio for an SPR move depends only on factors linked to the P-X and P’-X branches. If the rest of the posterior is also local, posterior ratios for two consecutive moves on different parts of the tree are independent of move order. In a maximum-parsimony context, this motivates the parallelization strategy used in the program matOptimize: it independently evaluates the potential removal of mutations of many SPR moves concurrently, then applies the largest possible subset of non-conflicting beneficial moves. In Delphy’s MCMC framework, we periodically partition the tree, apply local MCMC moves in each partition in parallel, then reassemble the whole tree for global moves (Figure 1E). This preserves correct sampling because it is equivalent to attempting local moves sequentially in some arbitrary interleaving (Figure 1F). By ensuring that any EMAT partitioning can be proposed, we maintain ergodicity of the underlying Markov chain (a mild approximation must be introduced when coupling to a coalescent prior—see SI for details).

Delphy’s EMATs facilitate efficient proposals of large-scale beneficial rearrangements through a process we call “*mutation-directed SPR* (mdSPR)” (Figure 1D), which enhances convergence without requiring fine-tuning of MCMC move parameters. An mdSPR move generalizes the core idea of UShER from maximum-parsimony to a Bayesian context: instead of placing a new sample at the most parsimonious attachment point, mdSPR proposes arbitrary attachment points while biasing towards those that require fewer mutations, even when far away. Because mutations are explicitly represented, Delphy quickly evaluates the minimum number of mutations required for any proposed P’-X branch in an SPR move. Assuming a Jukes-Cantor model with no site-rate heterogeneity and dominated by parsimonious mutational histories, one can write a closed-form conditional posterior for P’ (blue) that can be sampled to propose a concrete regrafting point P’. The Jukes-Cantor-inspired proposal distribution closely tracks the real conditional posterior for P’ (red). Hence, the MH criterion achieves high acceptance probabilities even when P’ is far from P, with occasional rejections correcting the bias in the approximate proposal. We note that mdSPR’s robustness degrades when many tip dates have large uncertainties, so Delphy is currently unsuitable for analysing such datasets.

Finally, Delphy introduces *N-pruning*, a novel technique for handling missing data that maintains the speed and parallelizability of individual moves by preserving the genetic prior’s functional form. Naively, an EMAT fully specifies the sequence at each tip, imputing the state of sites with missing data. However, during SPR moves on subtrees with missing data, such imputation would require *global* rearrangements, as for example, in an SPR on a subtree whose tips have no data at site *ℓ*, while the state of that site at P and P’ differs. Delphy circumvents this tension by not imputing the states of sites with missing data for such subtrees. Since the state at the root of such a subtree must evolve into *something*, integrating over all possible mutational histories of those sites over that subtree causes the associated branch and mutation factors to disappear from the genetic prior while preserving its functional form (see lower-right corner of Fig 1B). This integration is equivalent to *partially* performing Felsenstein pruning only on those sites and subtrees which end in missing data. Conceptually, if we view an EMAT as a product of *site trees* (one per site) then N-pruning recursively prunes all branches on such site trees that lead to tips with missing data (Supplementary Figure 13). The missations on an EMAT coincide with the points where this pruning occurs. The main drawback of N-pruning is that it adds some bookkeeping and complexity to SPR moves (see SI). A second limitation is that Delphy cannot efficiently handle partially ambiguous states (e.g., Y = C or T), and instead treats them as missing data. However, in our benchmark datasets, such partial ambiguity appears quite rare, so we do not consider this a significant limitation.

### Delphy makes Bayesian phylogenetics accessible worldwide via a web-based interface and analysis workflow

Delphy provides a fully web-based implementation of Bayesian phylogenetics, eliminating the need for installation or compilation, and ensuring that data stays with the data generator. Unlike similar tools, it can run directly in a web browser as a client-side application (https://delphy.fathom.info). The only required input is a multiple-sequence alignment in FASTA or MAPLE^25^ format, which must include dates (possibly inexact) at the end of the sequence IDs. Users can optionally provide a metadata file to annotate samples with categorical labels (see Methods). A web front-end, written in Typescript, processes these inputs and communicates with Delphy’s computational core, written in C++. This core, precompiled into WebAssembly, runs seamlessly in the browser, not in a remote server, so data never leaves a user’s machine. The front-end manages execution, retrieves results, and provides an interactive interface to explore results.

Delphy’s web interface guides users through the entire workflow, from data input to interactive analysis and result export (Figure 2). Users can monitor the core MCMC run for burn-in and convergence, analyze outputs interactively, and export publication-ready summaries and raw outputs. Built-in analyses include standard Bayesian phylogenetics outputs, such as the maximum-clade-credibility tree (MCC, calculated at interactive speeds through clade fingerprinting—see SI), the key lineages (identified with high-support inner nodes of the MCC), distributions of inferred time for tips with uncertain dates, time-to-most-recent-common-ancestor (tMRCA) estimates for the whole dataset and sublineages, and ancestral state reconstruction for metadata. Delphy’s explicit-mutation representation enables additional analyses, including the identities and time distributions of mutations between any two nodes, as well as experimental visualizations like lineage and mutation prevalence curves (see Methods). Throughout, the interface emphasizes inference uncertainties, by displaying posterior supports, 95% highest-posterior density (HPD) ranges for time and prevalence distributions, and ambiguous mutation placements – while keeping these details unobtrusive for occasional users. We expect to continuously improve and extend these automatic analyses, and our web-based distribution model will make these enhancements immediately available. (Previous versions can be run locally for reproducibility; see Code and Data Availability.)

**Figure 2:**
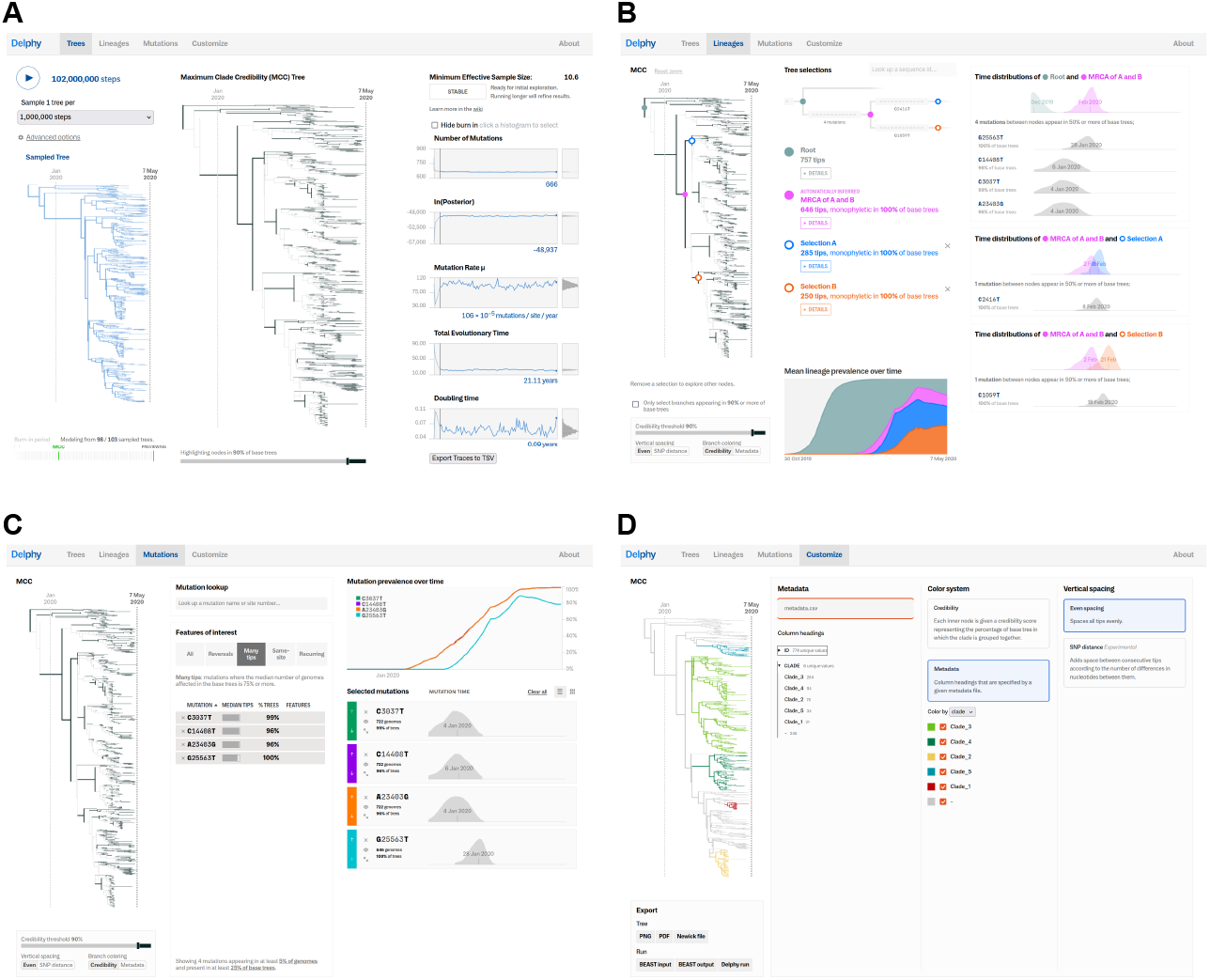
Delphy’s web interface. **A**. The *Trees Panel* is where the MCMC run is controlled and its progress monitored. After ingesting a dated multiple-sequence alignment in FASTA format, Delphy constructs a rough initial tree (left). Pressing the Play button starts the run. The Maximum-Clade-Credibility (MCC) tree (middle) shows an up-to-date summary of sample posteriors, with high-confidence nodes highlighted. Traces of various observables (right) allow for monitoring progress. A simple heuristic suggests a cutover point between the burn-in and production phases of the run, which can be overridden if needed (see Methods); all subsequent analysis uses only the production samples. **B**. The *Lineages Panel* is a lineage-centric view of the run results. A user can hover over the MCC (left) to select up to two nodes (blue and orange) and their common ancestor (pink). A node on the MCC stands for the most-recent common ancestor (MRCA) of its downstream tips, and the high-support nodes correspond to distinguishable lineages. The dominant relationships between these nodes are summarized in a minimap (top center). Their time to MRCA (tMRCA) distributions, as well as the identity and time distribution of the mutations between these nodes are shown (right). Using the underlying population model, Delphy estimates the time-varying distribution for which of the selected nodes is the closest ancestor of a random additional sample, i.e., a lineage prevalence histogram (bottom center; see Methods). All distributions can be probed further by hovering. **C**. The *Mutations Panel* is a mutation-centric view of the run results. Delphy automatically identifies potentially interesting mutations (center) and selects ones that rise to high prominence. For each selected mutation, the time distributions (bottom right) and their prevalence (top right) is shown. Hovering over a mutation shows the uncertainty in the prevalence curves and the approximate point on the tree where the mutation occurs. **D**. The *Customize Panel* provides a way to annotate the MCC with categorical metadata, such as clades or geographical labels. A naïve Fitch parsimony algorithm propagates metadata to the interior nodes of the MCC (colored tree on the left). Separately, a user can export a view of the MCC (in PNG or PDF formats for direct use, or in Newick format for post-processing). More advanced users can export the raw run results in BEAST2 format, for post-processing, or in a custom “Delphy’ format that can be reloaded later.

### Delphy reproduces the output of existing Bayesian tools at ∼1,000x speed

We expect Delphy’s results to be statistically indistinguishable from those of existing tools for the models we have implemented, and to remain so as we further extend it with other models. This is because Delphy samples the same distribution of trees sampled by existing Bayesian tools once the explicit mutation events are projected out. To facilitate direct comparisons, Delphy provides an option to export the XML input and output files for an equivalent BEAST2 run.

We benchmarked Delphy against BEAST2, a SOTA Bayesian phylogenetics tool and IQ-Tree 2 + TreeTime, the maximum-likelihood core of the NextStrain pipeline, running on published SARS-CoV-2, Zika, and Ebola datasets. These include 757 SARS-CoV-2 sequences from early 2020 ^45^, (2) 174 Zika sequences from 2015-2016^4^, and (3) 81 Ebola sequences from early 2014^3^. We detail the results for the SARS-CoV-2 dataset below. (See Supplementary Figure 2-4 for Zika and Ebola). All three datasets are available as demos at https://delphy.fathom.info.

The Delphy- and BEAST2-inferred MCC trees for the SARS-CoV-2 dataset show excellent agreement (Fig 3A and 3B). The MCCs, point summaries of the entire posterior distribution, have the same overall structure and timing. Note that the vertical rearrangements of the low-support nodes are not significant, and are expected even between two independent BEAST2 runs. The maximum-likelihood tree (Figure 3C) also captures the essential lineage structure, but under that approach the tMRCAs of major lineages are inferred less accurately.

**Figure 3:**
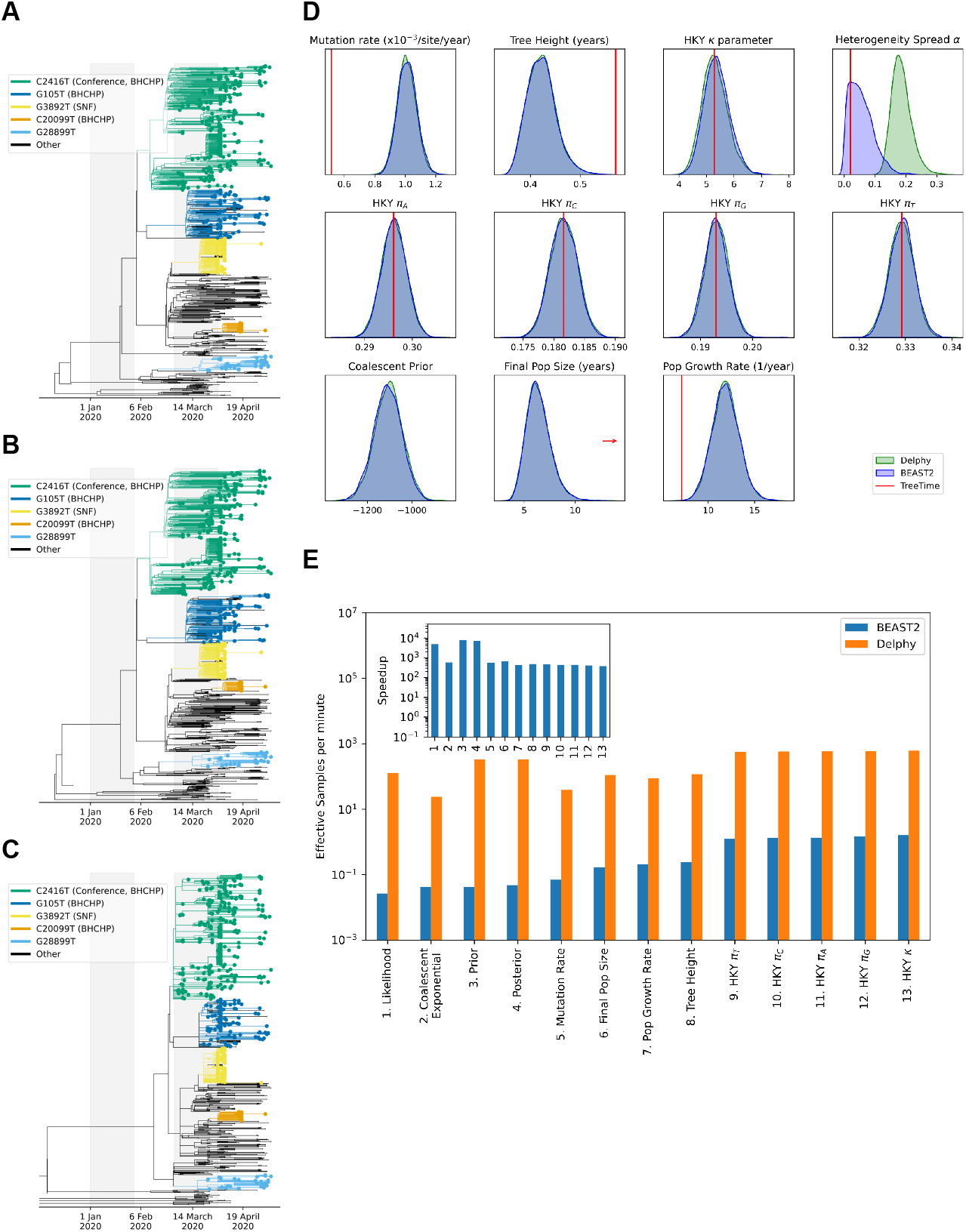
Delphy is consistent with existing Bayesian tools. **A**. Maximum-Clade-Credibility (MCC) tree for 757 SARS-CoV-2 samples from Lemieux et al. (Lemieux)^45^ as produced by Delphy (compare to Fig 3A in Lemieux^45^). **B**. Analog for BEAST2. **C**. Maximum-likelihood timed tree, as calculated by IQ-Tree 2 and TreeTime. **D**. Distributions of key observables, as produced by Delphy (green) and BEAST2 (blue), and maximum-likelihood estimate (red); the discrepancy in the site-rate heterogeneity arises from Delphy’s use of a continuous Gamma model vs. a 4-category discrete approximation in the BEAST2 run (Methods). **E**. Runtime efficiency of Delphy vs BEAST2.

The distributions of several key parameters, such as mutation rate, also show excellent agreement between Delphy and BEAST2 (Figure 3D). The only exception is the site-rate heterogeneity parameter alpha, due to a deliberate model difference: Delphy uses a continuous Gamma model whereas BEAST2 uses a discrete 4-category approximation owing to technical limitations. Direct comparison of runs with no site-rate heterogeneity shows no significant discrepancies (Supplementary Figure 1). In contrast, maximum-likelihood methods appear to accurately infer some parameters but not others (particularly the population curve, and hence, the mutation rate and tree height).

While their outputs for the SARS-CoV-2 dataset are essentially the same, Delphy achieves a 1000x higher statistical efficiency than BEAST2, measured as the rate at which an MCMC run produces uncorrelated samples of the underlying posterior distribution (Figure 3E). This was measured by calculating the Effective Sample Size (as estimated by BEAST2’s LogAnalyser) per unit run time. This three-orders-of-magnitude speedup was robust across observables. For the smaller Zika and Ebola datasets, Delphy achieved ∼10x speedups as expected: its advantages grow with dataset size (see Discussion).

### Delphy’s computational core is flexible–mpox case study

Delphy shares the same mathematical foundations as other Bayesian methods, allowing it to eventually support a wide range of model variations at the same level of accuracy. These include diverse evolution models, advanced site-rate heterogeneity models, flexible population models (using coalescent, birth-death, or epidemic priors) and incorporating geography biases. For its initial release, Delphy’s uses a single-partition Hasegawa-Kishino-Yano (HKY) model for evolution^46^ and a coalescent model with exponential population growth (see SI for details).

Future updates will relax these restrictions based on users’ needs.

As a proof-of-concept of Delphy’s flexibility, its initial public release includes a model for analyzing human mpox virus (hMPXV) sequences^5,6^ (see Methods). Unlike most viruses, hMPXV accumulates mutations primarily through APOBEC3 activity in humans rather than replication errors^5^. APOBEC3 targets TC and GA dimers, mutating them to TT and AA dimers, respectively. Delphy formalizes this with a 2-partition model closely resembling the post-zoonosis model in O’Toole et al (O’Toole)^5^ and Parker et al (Parker)^6^—see Methods.

Delphy’s analysis of hMPXV-1 sequences, descendants of the spillover that led to mpox infections worldwide starting in 2022, closely aligns with BEAST results despite minor modeling differences. Figure 4 presents Delphy’s results for 177 hMPXV-1 sequences from Parker. While we omitted non-hMPXV-1 sequences to avoid having to model the zoonotic spillover event itself (see Methods), Delphy’s agreement with BEAST is excellent. The clade breakdown in the MCC is clearly recovered (Figure 4A). Despite the crudeness of Delphy’s simple parsimony on the MCC, the central conclusion of Parker is also clearly recovered: the spillover event at the root of this hMPXV-1 outbreak likely occurred in the the South-South (SS) region of Nigeria, in particular, in Rivers state (however, unlike the sophisticated phylogeography in Fig 5B of Parker, Delphy’s simple parsimony on the MCC lacks the power to establish this finer geographic nuance with high certainty). Most of the differences from Parker–such as the slightly more recent tMRCA estimates, the higher non-APOBEC3 mutation rate, and slightly faster doubling rate–can be attributed to the absence of the pre-spillover samples in Delphy’s analysis; we suspect that purifying selection in the sparsely sampled pre-spillover portion of the tree suppresses the observed non-APOBEC3 rate in Parker (see Methods). Indeed, adapting the BEAST run in Parker to include only hMPXV-1 samples and avoid modeling the spillover and phylogeographic movements yields much closer agreement (mean tMRCA of 17 March 2016, with 95% HPD 25 Aug 2015 - 8 Sep 2016; non-APOBEC3 rate of 7.8×10^-6^ mutations per site per year, with 95% HPD 6.5×10^-6^ to 9.1×10^-6^; APOBEC3 rate of 1.3×10^-4^ mutations per site per year, with 95% HPD 1.1×10^-4^ to 1.4×10^-4^; and doubling time of 1.8 years, with 95% HPD of 1.4 to 2.3 years). Similar agreement between Delphy’s 2-partition hMPXV model and BEAST’s is observed when applied to the earlier dataset from O’Toole (Supplementary Figure 6).

**Figure 4:**
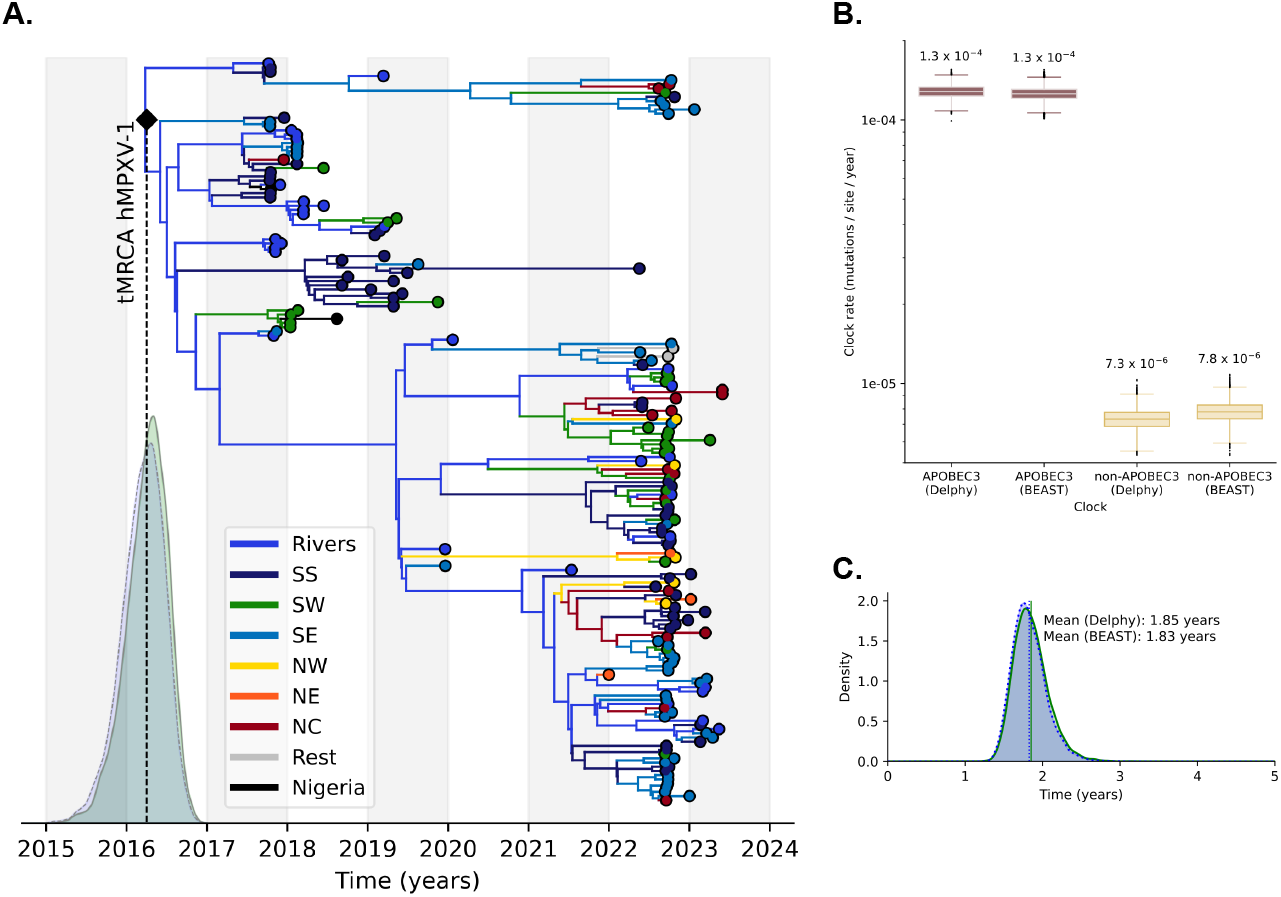
Delphy is flexible: **A**. Maximum-Clade-Credibility (MCC) tree for 177 hMPXV-1 samples from Parker et al. (Parker)^6^ as produced by Delphy (compare to Parker Figs 2A, 3A and Extended Data Figure 1). Time to most-recent-common-ancestor (tMRCA) distributions are for Delphy (green) and BEAST (blue). Tips and branches are colored by parsimonious assignment of Nigerian state on the MCC. All states besides Rivers in the South-South (SS) region are grouped by region. The original zoonosis event likely occurred in the South-South region, probably in Rivers state. **B**. Inferred mutation rates owing to APOBEC3 and non-APOBEC3 mechanisms (compare to Fig 2D of Parker). **D**. Inferred doubling time under exponential growth model (Delphy, green; BEAST, blue; compare to Fig 2C of Parker).

**Figure 5:**
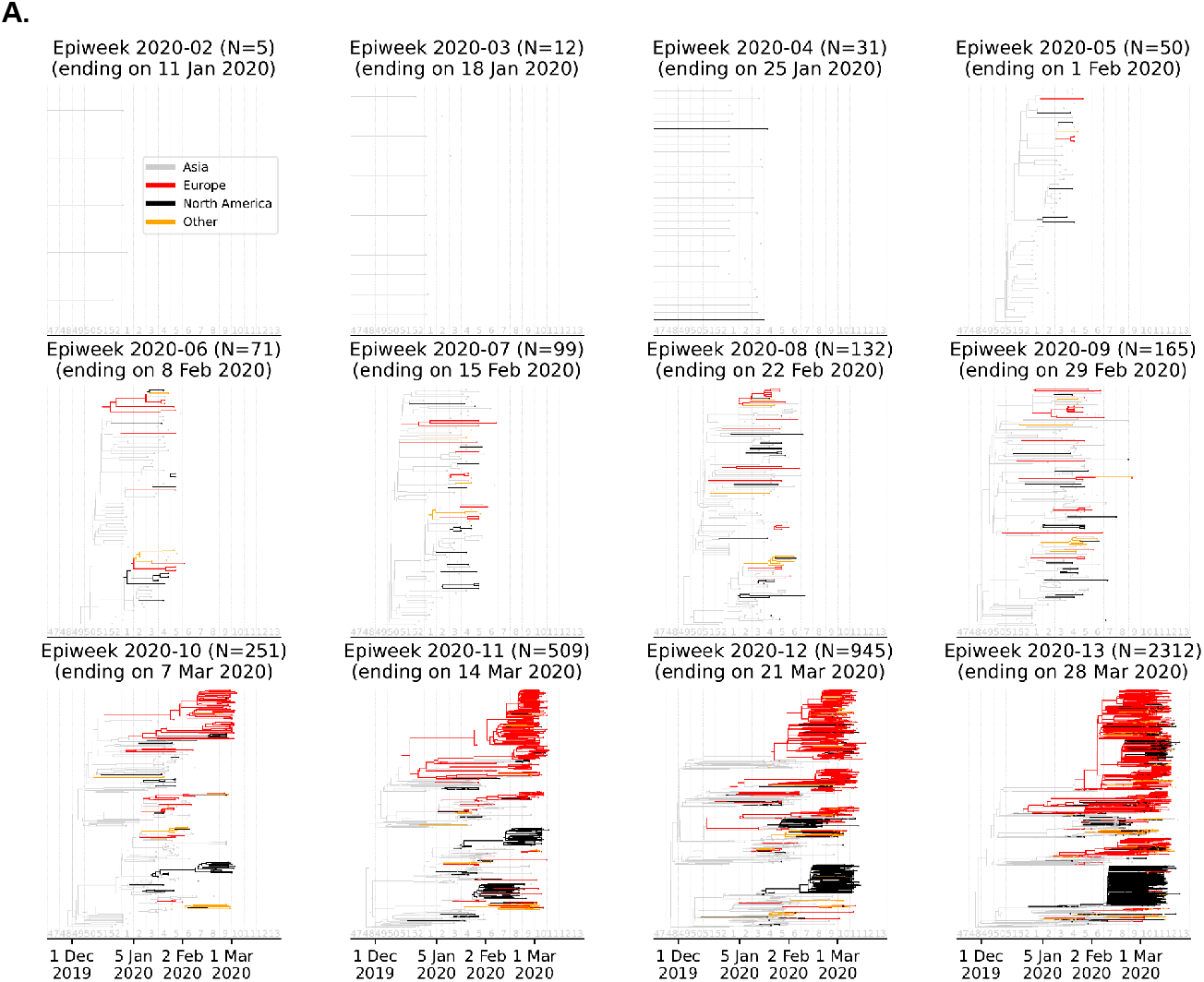
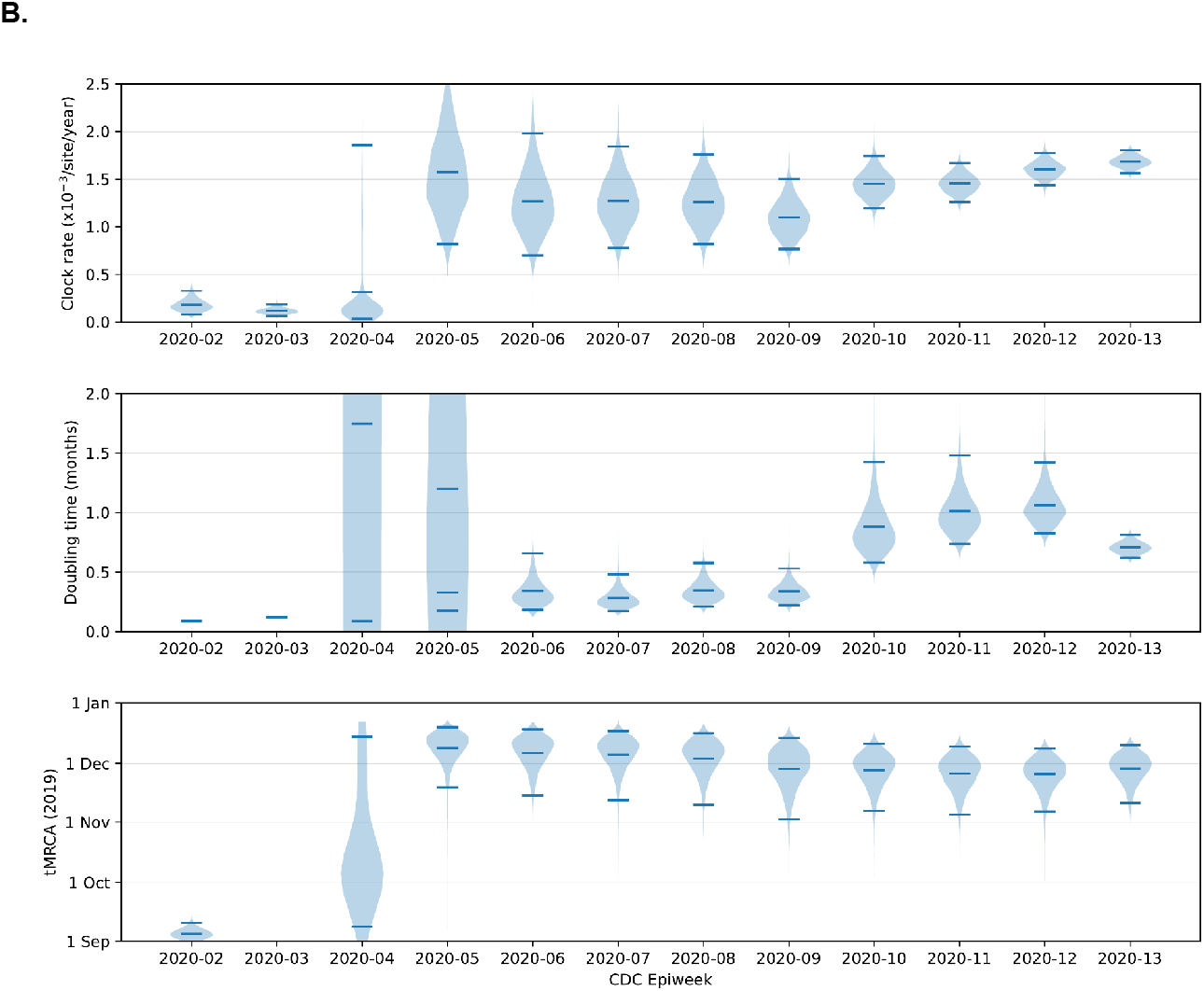
Delphy’s view of the first weeks of the COVID-19 pandemic. **A**. Maximum-Clade-Credibility (MCC) trees derived from the SARS-CoV-2 sequences submitted to GISAID up to 28 Mar 2020 (CDC Epiweek to 2020-13). Branches colored by continent as derived from naïve parsimony. Before CDC week 2020-04 (ending on Sat 25 Jan 2020), the data is clearly insufficient to infer trees reliably. From 2020-05, trees are relatively stable. Many independent introductions to both Europe and North America, as well as persistent local transmission chains, are already plainly visible by the end of CDC week 2020-06 (ending on Sat 8 Feb 2020). **B**. Distributions for mutation rate, doubling times and tMRCAs derived from GISAID data submitted by successive CDC weeks. The mutation rate is overestimated vs. consensus value of ∼1.0 x 10^-3^ / site / year, possibly owing to no masking of known problematic sites and minimal culling of outliers, as would happen near the early stages of an outbreak (see Methods).

**Figure 6:**
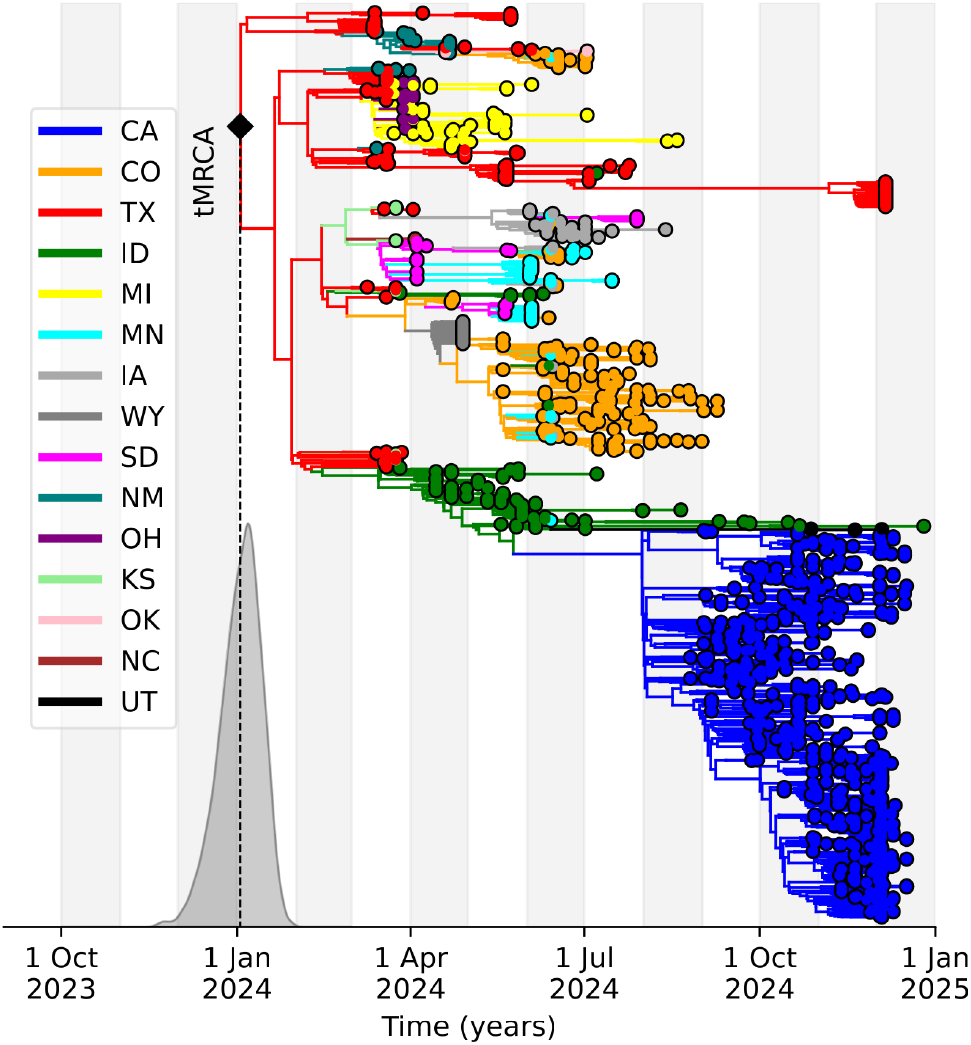
Delphy’s reconstruction of the H5N1 influenza outbreak in U.S. cattle, 2024-2025. Delphy’s phylogenetic tree, based on 1889 viral sequences, captures the key dynamics of the outbreak: an initial spillover in Texas in early 2024, followed by gradual state-to-state spread, with large localized outbreaks in Colorado around spring-summer 2024 and in California around summer-fall 2024. The reconstruction assumes an exponentially growing viral population.

### Delphy enables real-time outbreak response–SARS-CoV-2 and H5N1 case studies

To assess Delphy’s utility for real-time public health response, we analyzed all complete SARS-CoV-2 sequences submitted to GISAID 1 January to 28 March 2020 (CDC Epiweeks 2020-02 through 2020-13). To emulate the limited knowledge and urgency of outbreak investigations, we applied minimal processing–trimming sequence ends and removing clear outliers–without masking sites that were later identified as problematic (see Methods; see also Supplementary Figures 7-8 for results using all sequences *collected* by each CDC Epiweek).

We generated inferred trees and associated parameter distributions (mutation rates, doubling times and tMRCAs), as they would have been available to a public health responder in near real time (Figures 4A-B). Even a simple parsimony-based inference of ancestral locations on the MCC reveals that by 8 February 2020 (CDC Epiweek 2020-06), many independent introductions from Asia into Europe and North America had occurred, followed by onward local transmission. Despite potential biases from minimal data filtering, the basic parameters of the growing epidemic–including a mutation rate of 1-2 x 10^-3^ mutations / site / year, a doubling time of 10-20 days, and a tMRCA around early Dec 2019–would all have been evident by 1 February 2020 (CDC Epiweek 2020-05).

We are applying Delphy to the ongoing, uncontrolled H5N1 influenza outbreak in U.S. cattle, first detected in early 2024^47^. We have created a daily-updated Delphy analysis of the latest dated H5N1 sequences–as curated by Kristian Andersen’s lab and with dates and locations from GenBank–available at https://delphy.fathom.info/us-h5n1-latest. Figure 6 shows Delphy’s MCC tree for the latest available dated sequences as of mid-March 2025 (Methods), with states color-coded by inferred location using Delphy’s naïve parsimony over the MCC. Delphy accurately recovers the known outbreak dynamics^47–49^: originating in Texas farms and spreading slowly to other states, with a tMRCA into cattle around 2 January 2024 (95% HPD 12 December 2024 - 22 January 2024). The overall mutation rate across all segments is 4.1 x 10^-3^ / site / year, 95% HPD 3.8 - 4.3, consistent with previous influenza estimates^19^. Notably, Delphy assumes an exponentially growing viral population throughout, a constraint future work will address.

A significant portion of recent public H5N1 sequences lack precise dates or locations—about one third as of mid-March 2025, likely covering the preceding three months. Incorporating these sequences, which carry large uncertainties, presents formidable convergence challenges likely not unique to Delphy. While these could perhaps be overcome by imposing stronger priors on collection dates than implied by the sequence metadata, we have instead excluded these sequences from this paper and from our daily-updated trees. However, data generators, who presumably know precise metadata, can easily adapt our Delphy runs for private analyses. Should their policy on making precise metadata public change in the future, our public runs will automatically incorporate all sequences.

### Delphy makes pandemic-scale Bayesian phylogenetics possible

To be useful in a pandemic, Bayesian phylogenetics tools must accurately and rapidly handle very large datasets. To evaluate Delphy's performance on pandemic-scale data, we generated controlled synthetic trees of increasing size, and measured how quickly and accurately Delphy reconstructed them from the observed sequences at their tips.

We simulated SARS-CoV-2-like outbreaks expanding exponentially over 6 months, containing 100 to 100,000 samples, sampled at a rate proportional to the viral population at the time (Figure 7A; see Supplementary Figure 10 for analogous results in a constant-size population). We constructed the ancestry tree of these samples using coalescent simulation and evolved a 30,000-site random root sequence under an HKY model with a mutation rate of 10^-3^ mutations/site/year and a transition-transversion ratio of 5. This process produced N = 100 to 100,000 dated and complete sequences with a known true phylogeny.

**Figure 7:**
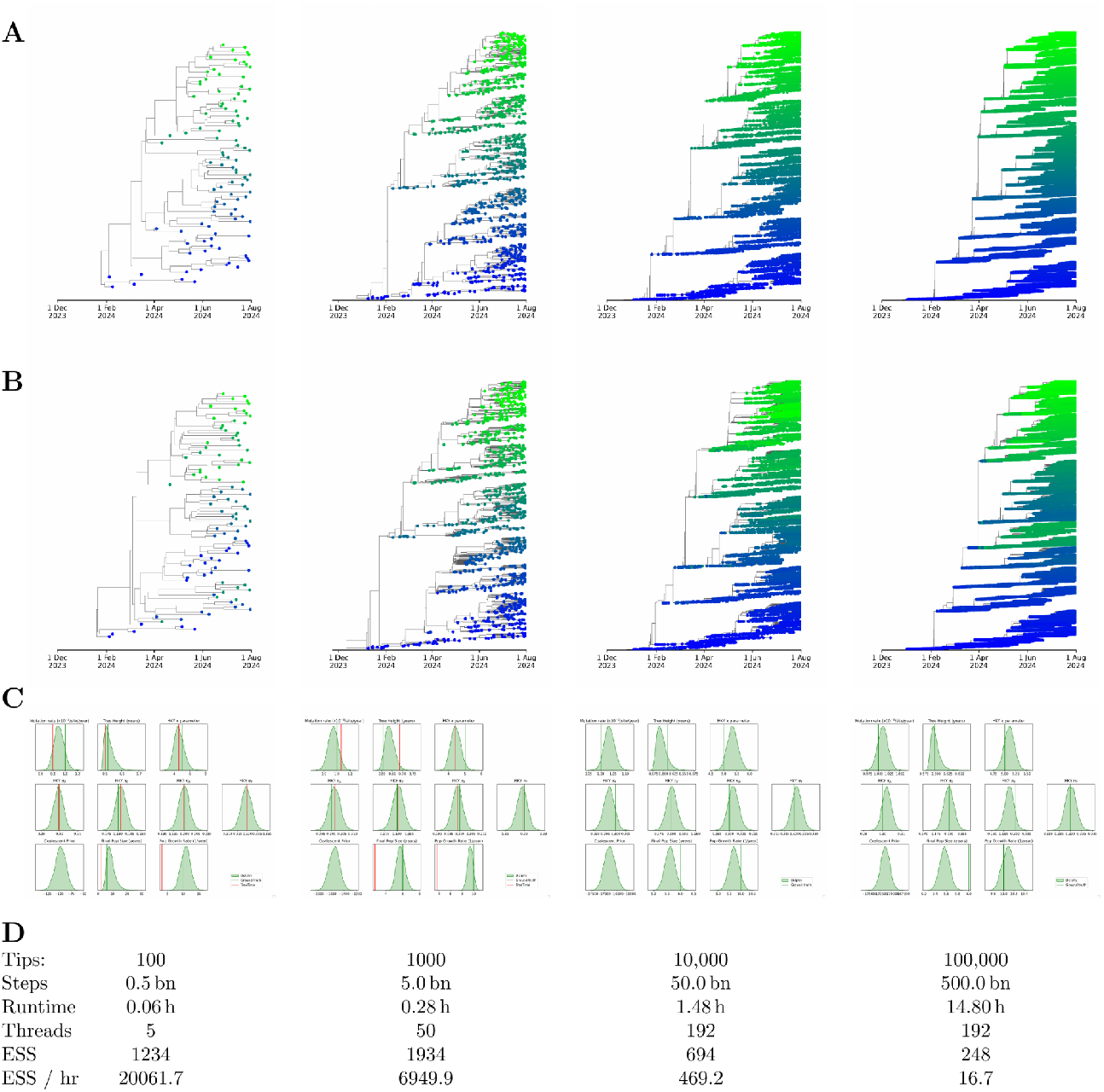
Delphy’s scaling behavior in the current proof-of-concept implementation. **A**. Simulated phylogenetic trees including 100, 1000, 10,000 and 100,000 sequences; tips colored blue to green by vertical ordering in simulated tree (see Methods). **B**. Maximum-Clade-Credibility (MCC) trees inferred by Delphy from the simulated tip sequences (corresponding tips in A and B have the same color). **C**. Inferred parameter distribution vs. known inputs (green) and maximum-likelihood estimates (red, only for 100 and 1000 tips). **D**. Run times and ESSs (posterior) for each run on a 96 vCPU AWS instance.

Delphy’s inferred phylogeny (Figure 7B) and model parameters (Figure 7C) closely matched the ground truth. The largest benchmark, with N = 100,000 samples, achieved an ESS of 248 in under 15 hours on a 96 vCPU AWS instance. For comparison, the largest Bayesian phylogenetics runs to date^50^ involved ∼40,000 sequences, used fixed topologies with polytomies, and took ∼3 weeks to complete (personal communication). Thus, even in this early implementation of the parallelized MCMC, Delphy processes pandemic-scale data fast enough to inform real-time public health decision-making, something that is not possible with SOTA Bayesian phylogenetic tools.

The main challenge in scaling Delphy to hundreds of thousands of sequences or more lies in numerical inaccuracies in the parallelized coalescent prior when too many branches are active simultaneously. Note, for instance, the discrepancy in the simulated and inferred final population size for the 100,000-sample benchmark in Figure 7C. We have mitigated this discrepancy by increasing the resolution of the underlying numerical discretization, but that adds significant computational cost, reducing ESS/hr for large runs (see Supplementary Figure 9).

Separately, our parallelization implementation needs further optimization; for example, the 100,000-sample run used less than 25% of the machine’s compute capacity at any given time.

## Discussion

Delphy demonstrates that Bayesian phylogenetics can be significantly accelerated for outbreak applications by an exact reformulation in terms of explicit mutations. By structuring every computational step in its inner loops in terms of mutations, Delphy simplifies each operation individually compared to established Bayesian tools. Yet, owing to their shared Bayesian foundation, Delphy’s results are statistically indistinguishable from those tools’ results. Our current implementation is already powerful enough to analyze real sequence data from emerging outbreaks. Conversely, for thinly sampled data spanning evolutionary timescales, an explicit-mutation approach remains exact but may offer a smaller efficiency gain, or even a loss, compared to existing Bayesian phylogenetics approaches.

Equally important, Delphy demonstrates that the barrier to entry for Bayesian phylogenetics can be significantly lowered, providing epidemiologists and public health workers worldwide the ability to use this powerful tool to inform real-time outbreak response. Our web application allows users to analyze their own data with little training and obtain publication-quality results in minutes. Two equally important factors contribute to this accessibility. First, distributing the computational core via WebAssembly in a web browser –despite resulting in 2-3x performance drop– eliminates the need to compile and install binaries, a substantial roadblock for our target audience. Second, the interactive front-end guides users through the calculation with carefully chosen defaults and automated downstream analyses, making it far more approachable. This is all carried out while respecting privacy, data ownership and cost sharing: no data ever leaves a user’s computer, and no organization has to unfairly bear the cost of analyzing others’ data.

Delphy is effective for typical genomic epidemiology datasets largely due to their low sequence diversity. For example, the 1-million-sequence maximum parsimony tree for SARS-CoV-2 in^23^ contains about 1 million mutations, meaning each sequence, on average, is just 1 mutation away from the nearest sequence in the tree. In such cases, the posterior distribution is dominated by nearly parsimonious trees^29,30,51^. While maximum parsimony loses timing information and cannot disambiguate between equally parsimonious trees, Delphy adds enough additional detail about the underlying evolutionary dynamics to address both shortcomings. By maintaining a Bayesian framework, rather than simplifying to maximum-likelihood, Delphy preserves the ability to accurately represent phylogenetic uncertainty and to incorporate additional layers of inference like phylodynamics, outbreak reconstruction or hypothesis testing. When confronted with datasets that are locally or globally far from parsimonious, Delphy degrades gracefully by remaining exact but slowing down, rather than silently introducing arbitrary artifacts and errors.

Delphy's treatment of missing data is novel, to our knowledge, and may be usefully applied in other contexts. At heart, N-pruning transforms a difficult problem with potentially enormous uncertainty into a simpler problem with little uncertainty. By obviating the need to impute missing data near the tips of the tree, while exploring explicit imputations of uncertain mutational histories in the tree’s interior, N-pruning avoids the key pitfall of using an explicit-mutation representation while maintaining its key benefits.

Delphy’s mdSPR move (Figure 1C) enables large-scale rearrangements of trees without requiring fine tuning of the move’s details; this robustness is key to opening Bayesian phylogenetics to a much broader audience than has previously been possible. One view on mdSPR is that it generalizes UShER's elemental step of finding the most parsimonious attachment point for any tip, with three key differences: (a) the global parsimony requirement is relaxed, leading to completely local moves that can be scaled; (b) because rearrangement is attempted continuously, there is little danger of an unfortunate early decision leading the search for a parsimonious tree into a suboptimal minimum; and (c) because of its Bayesian formulation, moves can only improve the efficiency of sampling, not change the sampled posterior distribution.

Delphy’s Bayesian foundations make it possible, in principle, to implement the myriad features and extensions that have accumulated in existing tools^16–18^. Our initial focus has been on enabling rapid understanding of emerging outbreaks undergoing exponential growth. In the near-term, we plan to extend Delphy to handle common complications arising in this scenario, including flexible non-exponential population models (e.g., Skygrid^52^), multiple partitions for codon effects, more flexible substitution models like GTR^53^ or UNREST^54^, nonstrict molecular clocks^55^, etc. We expect such features to be driven by users' needs, and welcome others' contributions towards this goal. However, the flexibility of existing tools, accumulated over decades, is unmatched, so investigations requiring such flexibility will remain within the purview of existing tools for the foreseeable future. During Delphy’s development, we did not see a clear path to efficiently implement Delphy’s key ideas of EMATs and parallel MCMC moves in existing tools like BEAST, but we remain hopeful that others might. Such an approach would save the enormous effort of recreating the features of existing Bayesian tools and thus benefit a broader community.

Finally, Delphy's parallelism implementation is a proof-of-concept with optimization needed to further improve performance. For example: when splitting and reassembling subtrees, we copy data instead of sharing it; we serially recalculate from scratch many intermediate quantities, such as log-posterior values, on tree reassembly; we have not yet attempted to use GPUs for massive parallelization; and we have not yet explored directly factorizable models of ancestry, such as birth-death and epidemiological models. Our belief is that the systematic pursuit of these kinds of improvements will – in the spirit of UShER and matOptimize for maximum parsimony phylogenetics and MAPLE for maximum-likelihood phylogenetics – soon unlock pandemic-scale Bayesian phylogenetics at unprecedented levels of detail.

## Code and data availability

Delphy is open-source. The code for its computational core and binaries for x86-64 Linux are available at https://github.com/broadinstitute/delphy. Its web interface is hosted at https://delphy.fathom.info, with source code available at https://github.com/fathominfo/delphy-web; the code includes instructions for local deployment. The ‘Delphy’ format is described at https://github.com/broadinstitute/delphy/blob/main/doc/dphy_file_format.md.

Ready-to-use Google Colab tutorials for downloading and formatting sequencing data from the NCBI Virus database, as well as user-provided data, are available at https://colab.research.google.com/github/broadinstitute/delphy/blob/main/tutorials/delphy_workflow.ipynb (blank notebook) and https://colab.research.google.com/github/broadinstitute/delphy/blob/main/tutorials/delphy_workflow_example.ipynb (example tutorial). These notebooks streamline the uniform formatting of sequencing data and the associated metadata for further analysis with Delphy, and then run Delphy using a pre-compiled binary. The output files can be visualized on the Delphy web interface.

All data and scripts for generating the figures in this paper are available at https://github.com/broadinstitute/delphy-2025-paper-data. Sapling, the special-purpose program we developed to efficiently simulate very large trees, is available at https://github.com/broadinstitute/sapling.

## Acknowledgements

We would like to thank all members of the Fathom Information Design team for their continued input in the design of Delphy. We also thank Alexei Drummond and Erick Matsen for highlighting the existing literature on explicit-mutation representations in Bayesian phylogenetics. We would also like to thank Ivan Specht and Frank Liu for valuable feedback on initial versions of this manuscript, and other members of the Sabeti lab for their input.

This work is made possible by support from Flu Lab and a cohort of generous donors through TED’s Audacious Project, including the ELMA Foundation, MacKenzie Scott, the Skoll Foundation, and Open Philanthropy. Funding was also provided by the US CDC Office of Advanced Molecular Detection Contract # 75D30122C14365 and Grant # NU50CK000629 (Pathogen Genomic Centers of Excellence); NIH/NIAID Grants # U19AI110818 (Genomic Centers for Infectious Diseases), # U01AI151812 (West African Research Network for Infectious Diseases), and # U19AI135995-07S2 (Consortium for Viral Systems Biology). I.O. is supported by the Wellcome Trust Hosts, Pathogens & Global Health program [Wellcome Trust, Grant number 218471/Z/19/Z] in partnership with Tackling Infectious Disease to Benefit Africa, TIBA.

## Methods

For brevity, we summarize the essentials of Delphy’s operation here, and refer the reader to the SI for full details.

### Explicit Mutation-Annotated Trees (EMATs)

Delphy represents trees internally as a collection of explicitly timed nodes and a reference sequence. Each node points to an earlier parent (except for the root) and to two later children (except for tips). Every point in the tree can be identified by a node and a time between the node and that of the node’s parent. Every such point has an associated sequence, which is encoded in terms of successive differences from the reference sequence. The reference sequence consists of an arbitrary string of L states from the set {A,C,G,T}. From there, the root stores a list of tuples (a,*ℓ*,b), at most 1 per value of *ℓ*, which record a difference at site *ℓ* between the state a of the reference sequence and state b of the root sequence; for convenience, we refer to these as mutations as being “above the root”. Below the root, every other node is annotated with an ordered sequence of mutations from the node’s parent to the node itself.

Each mutation consists of a tuple (a,*ℓ*,b,t) which records that the point immediately preceding the time t on the branch between the node and its parent has state a at site *ℓ*, while the point immediately following t has state b. With this structure, the sequence at any point x on the tree can be reconstructed by starting with the reference sequence, applying the mutations above the root, and then successively applying in order all the mutations encountered on the unique path from the root to x.

Each node also includes a set of missations, consisting of tuples (a, *ℓ*), at most 1 per value of *ℓ*, which record that for all tips downstream of the point immediately below the node’s parent, the state of site *ℓ* is unknown (“missing”), as represented by an N in the input data, but this is not true of the point immediately above the node’s parent; there, the state at site *ℓ* is a. For a given tree topology, mutational history and tip sequences, the missations on the tree are completely determined. Missations are encoded using two complementary structures: (a) an ordered sequence of disjoint, nonconsecutive half-open intervals [*ℓ*_start_, *ℓ*_end_), and (b) a map of sites *ℓ* to states a containing an entry for *ℓ* only when the state a differs from the corresponding state in the reference sequence. This representation reflects that (a) missing data in real-world sequences typically consists of a few relatively long “gaps”, and (b) that if we arrange for the reference sequence to not deviate much from the root sequence, and the time between the root and any node is small relative to the mutation rate, then the map of differences from the reference will typically be sparse.

To support uncertain tip dates, we associate a minimum and maximum time with each tip. These are equal when the tip date is known exactly.

EMATs have evident consistency requirements: (1) a node’s time must be later than its parent’s; if a mutation at site *ℓ* has a starting state a, then the sequence deduced by the above procedure at the point immediately above the mutation must have state a at *ℓ*; (3) similarly, if a missation from state a at site *ℓ* is associated with a node, then the state immediately above the node’s parent, as deduced by the above procedure, must be a, and there must not be any mutations at site *ℓ* downstream of the missation; (4) if a missation for site *ℓ* is associated with a node, then neither the node’s sibling nor any of its ancestors or descendants has a missation at *ℓ*. We note that a set of experimentally determined sequences consisting exclusively of states {A,C,G,T,N} and having precise timestamps can be represented exactly and compactly by a suitably constructed EMAT.

### Posterior distribution

Delphy samples trees T and associated model parameters θ using MCMC according to the following (unnormalized) posterior distribution:

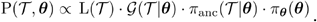

The factors are as follows:

- π_θ_(θ) is the prior distribution for the model parameters θ.
- π_*anc*_ (*T*|θ) is an ancestry prior for the tree’s topology given the model parameters θ.
- *G*(*T*, θ) is a genetic prior for the particular mutational history decorating the EMAT T.
- *L*(*T*) is the likelihood of the data given the EMAT T; since Delphy currently restricts tip sequences to be either definite (A,C,G,T) or completely missing (N), this likelihood is simply 1 if the EMAT is consistent (see above) or 0 otherwise.

The first two priors are described below. The genetic prior is the probability that a random root sequence evolved with the evolution model parametrized by θ produces the mutational history in EMAT T. Explicitly,

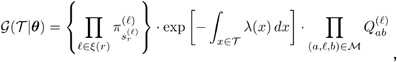

where:

- 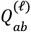 is the rate at which state a at site *ℓ* transitions to state b, as parametrized by the evolutionary model parameters in θ.
- 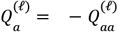 is the rate at which state a at site *ℓ* transitions to any other state.
- 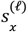 is the state of site *ℓ* at point x on T.
- ξ(*x*) is the set of sites for which at least one tip below point x on T is informative (i.e., its state is not missing).
- 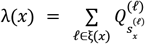 is the sequence-dependent genome-wide mutation rate at point x on T (see N-pruning discussion below).
- 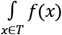 is an integral of a function *f*(*x*) defined at every point on the tree, defined as the sum of time integrals along each branch of T.
- 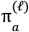 is the probability that the root sequence has state a at site *ℓ*.
- *r* is the root of the tree.
- *M* is the set of all mutations across the tree.

The SI describes in further detail the motivation for these choices, considerations for efficient calculations, and a discussion on its equivalence to the standard tree likelihood based on Felsenstein pruning.

### Parameter and Ancestry Priors

Delphy’s priors, π_θ_ (θ) and π_*anc*_ (*T*|θ), are currently as follows:

- Tip dates are à priori uniformly distributed between their minimum and maximum times.
- Tip times are, à priori, uniformly distributed between their minimum and maximum times.
- The transition rate matrices have the form 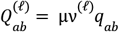, where μ is an overall per-site mutation rate, the quantities ν^(*ℓ*)^ are site-relative rates, and *q*_*ab*_ are the normalized transition rate matrix elements of an HKY evolution model with transition-transversion rate *κ* and stationary state frequencies π_*a*_ . This form implies a strict molecular clock.
- The mutation rate μ has an improper uniform prior.
- The site-relative rates ν ^(*ℓ*)^ are either fixed to 1 (no site-rate heterogeneity), or have à priori ν^(*ℓ*)^∼*Gamma*(α, α), with α∼*Expo*(1) à priori.
- The HKY parameters are chosen such that, à priori, {π_*a*_ } ∼*Dir*(1, 1, 1, 1) and *log*(*κ*) ∼ *N*(1, 1. 25^2^).
- The ancestry prior is a standard coalescent prior with an exponentially growing population curve 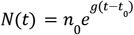, where *t*_0_ is the time of the latest tip (see below for a discussion on parallelization). The final effective population size *n*_0_ has an an improper 1/x prior, while the growth rate *g* has a Laplace prior with mean 0.001 per year and scale 30.701135 per year.

These choices are suitable for the early stages of an exponentially growing viral outbreak, and are currently fixed in Delphy, but their details are not essential. We expect to evolve Delphy to make prior specification more flexible in the future. Most of the above details are the same as the defaults provided by BEAUTi2, and coincide with those used in Lemieux et al^45^. The priors are discussed further in the SI.

### MCMC moves

Delphy samples trees and model parameters from the above posterior distribution using Markov Chain Monte Carlo (MCMC). We distinguish between *local* moves that affect only a few nodes, and *global* moves that affect the whole tree.

The following global moves are used:

- Gibbs sampling of mutation rate: The form of *P*(*T*, θ) when T and all model parameters except μ are fixed is simply a Gamma distribution with simple parameters, so μ may be directly Gibbs sampled (see SI for details).
- Evolution model parameters: Delta-exchange moves on the stationary frequencies π_*a*_ (pick a random pair of distinct states a and b and a displacement δ∼*U*(0, 0. 01), and propose 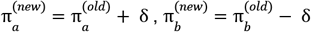, rejecting immediately if either proposed value is outside [0,1]). Scale moves on *κ* (multiply by a random factor δ∼*U*(3/4, 4/3)). These are performed in batches of 20, alternating between π_*a*_ and *κ*.
- Site-rate heterogeneity parameters: a series of 10 consecutive scaling moves on α, with factors δ∼*U*(0. 9, 1/0. 9), using an effective posterior where the dependence on ν ^(*ℓ*)^ has been marginalized out, followed by L independent Gibbs samples for { ν ^(*ℓ*)^}.
- Population model parameters: A batch of 100 moves interleaving scaling moves on *n*_0_ with a factor δ∼*U*(3/4, 4/3) and random walks on *g* with uniform step size of at most 1 / year.

The relative simplicity of global moves in the explicit-mutation representation, including the marginalization of {ν^(*ℓ*)^} for making moves for α, was first highlighted by Lartillot^34^. Full details of these moves are given in the SI.

The following local moves are used:

- *Node displacement*: pick a random node X and displace it forwards or backwards without crossing a mutation, changing the tree topology, or escaping the bounds for a tip date.
- *Branch reform*: pick a random branch and pick random new times for the mutations on the branch.
- *Subtree pruning and regrafting (SPR)*: pick a random node X (not the root) with parent P; prune the subtree rooted at X from attachment point P; pick a new attachment point P’ (see below) and reattach the subtree there; propose a new site-homogeneous Jukes-Cantor mutational history from P’ to X that is compatible with the fixed sequences at P’ and X, and rely on the Metropolis-Hastings criterion to accept/reject proposals so that the mutational histories follow the actual evolution model.

Full details and subtleties of the local moves are given in the SI. In particular, the mutational history proposal of the SPR move is implemented in a way that scales with the number of site differences between the sequences of P’ and X, not the genome size. The interaction of SPR moves with our handling of missing data is, unfortunately, quite involved.

A specific SPR move involves a proposal for the regrafting point P’. Many choices are possible, and while Delphy implements the standard subtree slide proposal^16^, it occasionally makes a mutation-directed SPR (mdSPR) proposal. As described in the main text (Figure 1D), the distribution of the point P’ is what would result if the evolution model were a site-homogeneous Jukes-Cantor model and the mutational history from P’ to X were parsimonious, while leaving all else unchanged. In practice, we adapt this basic idea in three key ways:

- When scanning a tree to count the minimum number of mutations that need to exist in the P’-X branch, we ignore mutations on sites that are missing at X (no spurious mutations will be introduced owing to N-pruning).
- We often restrict the new attachment point to be in the vicinity of the old attachment point, defined as all points reachable from the old attachment point without crossing more than 1 mutation. About 1% of the time, we do scan the entire tree, which is expensive but allows for large-scale rearrangements.
- We dampen the proposal distribution by an “annealing” exponent *f* = 0. 8 ; this proposes beneficial rearrangements less eagerly than the Jukes-Cantor model would suggest, but leaves enough flexibility for errors in this proposal to survive the Metropolis-Hastings criterion and be accepted. This is particularly helpful for accepting rough but highly beneficial large-scale rearrangements, then applying fine-grained corrections to them.
- The exact proposal probability for a branch involves expensive incomplete Gamma functions; we approximate those by the mid-point value of the underlying integrals.

Explicit procedures and formulas may be found in the SI.

### Missing data: N-pruning and missations

Missing data significantly complicates the use of an explicit-mutation representation. The straightforward approach, where full mutational histories are inferred and missing data is imputed, can be costly in practice: SPR moves relating to a subtree where all tips are uninformative at a site *ℓ*, and which transfer that subtree from a point where that site has state a to one with a different state b have to either (a) introduce a spurious a->b mutation along the attachment branch, leading to high rejection rates, or (b) resample the mutational history of the site over the entire subtree. Neither solution is satisfactory, especially considering that it is not unusual for a subtree to contain thousands of (consecutive) sites with missing data.

Delphy’s solution is to partially integrate over all possible state assignments of sites at nodes below which there is no information. Notably, this preserves the properties of the form of the posterior that make an explicit-mutation representation worthwhile: locality and factorizability. Concretely, the effect of this partial integration is merely to add the restrictions to *ℓ* ∈ ξ(*x*) in the formulas for *G*(*T*, θ). The partial integration starts from the tips and proceeds upstream until reaching a node that has an informative downstream tip. The point at which this integration stops is precisely the point at which an EMAT has a missation.

Although the bookkeeping complications are substantial, it is possible to efficiently update the missations on a tree as subtrees are pruned and regrafted and mutational histories are changed. Because topological rearrangements may introduce, move or remove missations, they affect the proposals and acceptance probabilities of all MCMC moves. Full details are given in the SI.

### Parallelizable coalescent

Delphy uses the usual Kingman coalescent as its ancestry prior:

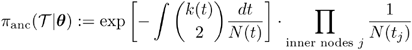

Here, k(t) is the number of active lineages at time t (i.e., the number of branches whose endpoints bracket the time t), while N(t) is the product of effective population size and generation time. The above integral can be calculated exactly for many functional forms of N(t) by partitioning the time range into intervals whose endpoints coincide with each node and tip of the tree. However, the integral intrinsically couples all the active branches at any given time through the quadratic dependence on k(t), which significantly hinders parallelization. In particular, given a partitioning of a tree into subtrees, the acceptance probability of an SPR move in one subtree that prunes a branch at t_1_ and grafts it at t_2_ depends on the configuration of all other subtrees with branches overlapping the range between t_1_ and t_2_. Taken at face value, this coupling only allows for efficient parallelization if the partitioning scheme produces subtrees that do not overlap in time, a severe restriction.

To circumvent this problem, Delphy introduces an augmented coalescent prior, whereby the coupling between subtrees is mediated by intermediate variables. For a fixed value of these intermediates, the subtrees can evolve in parallel with complete independence. However, for correct sampling, the intermediate variables must also be sampled when the subtrees are reassembled, a step we incorporate when performing all other global MCMC moves. We emphasize that while our present scheme is correct and permits parallelization, we think it is neither the only way nor the best way to parallelize the coalescent prior: we expect better approaches to emerge in the future.

Delphy’s current scheme proceeds in two steps. First, the integral in the coalescent prior is discretized into a Riemann sum, with the time domain divided into a few hundred cells of width Δ:

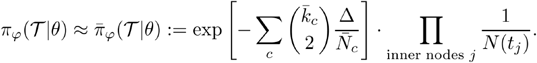

Here, c is the index of each cell, while the barred variables 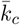 and 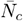 are the averages of k(t) and N(t) over the range of cell c. We use this form of the coalescent prior for global moves, e.g., those involving changes to the population parameter models. For local moves, we periodically partition the tree into P partitions, indexed by p, as discussed in the main text. We correspondingly split 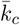 to into values 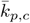, which encode the average number of active lineages in cell c and partition p, such that 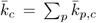. Denote by P_c_ the number of subtrees that span cell c. We introduce a set of augmented variables 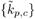 for every cell c spanned by partition p and propose the following joint distribution for them:

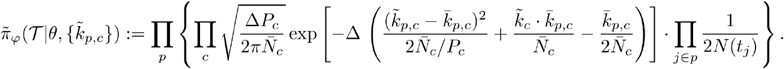

The augmented variables 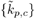 are Gaussian random variables that couple to 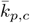, so one can analytically integrate the above distribution over all values of 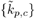. The mean, dispersion and couplings of 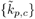 are chosen so that the result of integrating over all values of 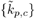 is the Riemann sum above. Hence, sampling the augmented system and then dropping the sampled values of 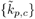 is equivalent to sampling the original system.

The augmented coalescent prior has two key properties:

- For fixed values of 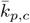, one can directly Gibbs sample values of 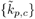.
- For fixed values of 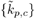, the distribution is a product of independent factors, one per partition.

As a result, sampling over subtrees can proceed completely independently and in parallel for fixed values of 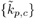, as described in the main text, and the 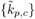 can themselves be directly sampled every time the subtrees are reassembled to execute global moves. See the SI for full details of this procedure, and a discussion on a subtle interaction between the precise partitioning scheme and correct sampling.

### Lineage and mutation prevalence curves

Delphy's interface allows a user to mark one or more of a tree's inner nodes, indexed by A, each of which can be regarded as the founding virion of a particular lineage; the root is always implicitly marked in this way. It then calculates “lineage prevalence curves” *u*_*A*_ (*t*) for all marked nodes: these are the probability distribution of the closest marked ancestor of a random member of the total viral population at time *t*. Here, we describe how that calculation is performed.

First, consider a single posterior tree. Let *k*_*A*_ (*t*) be the number of active lineages at time *t* whose closest marked ancestor is A. In the coalescent model, if we draw a random member of the total viral population at time *t*, the probability that it coalesces with one of these *k*_*A*_ (*t*) active lineages between times *t* − *dt* and *t* is given by [*k*_*A*_ (*t*) / *N*(*t*)] *dt*. Hence, the probability that it does not coalesce at all is given by 1 − [*k*(*t*) / *N*(*t*)] *dt*, where 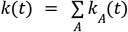.Thus, *u*_*A*_ (*t*) obeys the following differential equation:

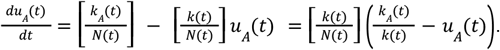

The initial condition that *u*_*A*_(*t*) = 0 for all A and all t preceding the root time completely determines *u*_*A*_(*t*). The solution to this differential equation is an exponentially moving average of [*k*_*A*_(*t*)/*k*(*t*)] with decay rate [*k*(*t*)/*N*(*t*)], as follows:

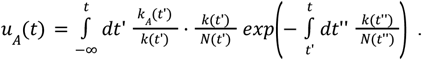

Delphy solves the above differential equation numerically, but the analytical solution makes the meaning of *u* (*t*) clearer. When *k*_*A*_ (*t*) is large relative to *N*(*t*) and the rate at which these two quantities vary is low, the tree can be thought of as densely sampled at time *t*: since the rate of coalescence is high but the number of active lineages is not varying quickly, most active lineages are immediately upstream of a tip. In that regime, *u*_*A*_ (*t*) closely approximates *k*_*A*_ (*t*)/*k*(*t*), which is essentially the lineage prevalence of A that would be estimated by direct case counts. However, outside that regime, *u*_*A*_ (*t*) integrates all the available information from the population model and the rest of the tree to smooth out such direct case count estimates.

When a user selects an inner node X on the MCC in Delphy's Lineages panel, we interpret that as the smallest clade that contains the particular set S of tips downstream of X in the MCC.

Correspondingly, in each posterior tree, that action marks the MRCA of the tips S, which is well-defined for all posterior trees. When a single inner node in the tree is marked multiple times, ties are broken arbitrarily. The corresponding lineage prevalence curves are calculated for each posterior tree, and the mean and 95% HPD range of each such family of curves is displayed to the user.

A completely analogous procedure is used to calculate the “mutation prevalence curves” in the Mutations panel. For a selection mutation (a,*ℓ*,b), we calculate the probability *u*(*t*) that a random member of the total viral population at time *t* has as its closest ancestor in the tree a virion where site *ℓ* has state b. As above, *u*(*t*) is calculated for all posterior trees, and the mean and 95% HPD at each value of *t* is displayed to the user.

### Automatic detection of burn-in cutoff

The Delphy web interface implements a simple but effective heuristic for suggesting a location for the burn-in cutoff in the MCMC run. For a particular observable, we first calculate the mean and standard deviation during the second half of the run. Then we find the latest time for which the observable is over 5 standard deviations from the mean. Finally, we find the earliest time thereafter at which the observable is within 2 standard deviations of the mean. We repeat this process over several observables (currently log-posterior, mutation rate and total evolutionary time) and pick the latest resulting time as the suggested cutoff. Unless and until a user overrides this suggestion, the cutoff is continuously updated, although not unsurprisingly, the suggestion stops changing once the run is well into the production phase.

The intent of the above heuristic is to identify a point near the end of the initial burn-in portion of the trace, and then advance to the earliest subsequent “normal” part of the trace. Empirically, we have found this procedure to make similar choices for the cutoff as a human would.

Importantly for Delphy’s accessibility goal, the cutoff suggestion is made automatically: experience with early users showed that requiring a user to manually pick a cutoff led to either needless friction or, worse, no cutoff being picked at all, leading to important biases in the result.

### Delphy input formats

Delphy can currently read a multiple-sequence alignment (MSA) of input sequences in one of two formats: FASTA and MAPLE^25^. In both formats, a full sequence ID is extracted from each description line from just after the initial ‘>’ up to but excluding the end of the line or the first space (hence, sequence IDs must not contain spaces). The full sequence ID must consist of fields separated by a vertical bar (‘|’). The first field is used as a short sequence ID in the Delphy user interface and for metadata annotation; the last field must be a date specification; remaining fields are ignored. A date specification can be in one of the following four formats: exact date (‘2025-01-24’), month (‘2025-01’), year (‘2025’) and arbitrary date range (‘2025-01-20/2025-01-24’).

Metadata should be input as a comma-separated value (.csv) or tab-separated value (.tsv) file with a header row as its first line. One column should be called “id” or “accession” (case-insensitive); its values should be one of the short sequence IDs from the input MSA. The remaining columns may have any names and values. Values may be quoted with double-quotes (“), and missing values may be indicated by leaving a column empty or by using the special values ‘-’, ‘noknown’ or ‘none’ (case-insensitive).

Sample MSA and metadata files can be downloaded for the demos on Delphy’s landing page.

### APOBEC3-aware evolution model for mpox

Inspired by^5,6^, we have added an option in Delphy to use a specialized evolution model suitable for mpox sequences, which can be activated in the web interface under “Advanced Options”.

We first classify each site as having APOBEC3 context according to the sequence of the first sample in the input file (with missing sites replaced by the consensus across all input sequences). A site has APOBEC3 context if it and the preceding site form a T**C** or a T**T** dimer, or if it and its subsequent site form a **G**A or **A**A dimer. For sites without APOBEC3 context, we use a transition rate matrix for a Jukes-Cantor evolutionary model with mutation rate μ and no site-rate heterogeneity:

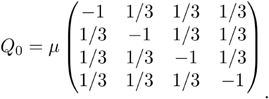

In sites with APOBEC3 context, we add transitions from C->T and G->A at a rate μ* as follows:

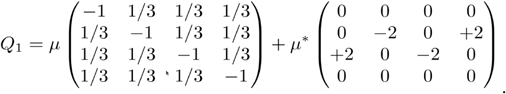

The factor of 2 is chosen to match the conventions used in the BEAST runs to define the APOBEC3 rate (the effective rate at which transitions would be observed in a sequence of APOBEC3 sites that are 50% C/G and 50% T/A).

The above setup retains the essence of that of the BEAST runs in O’Toole et al and Parker et al^5,6^ but differs minimally in its details. By deciding on the APOBEC context using the first sequence, we avoid having to previously estimate an ML tree and extract its root sequence (the assumption is that the APOBEC3 context of the vast majority sites is stable throughout the entire tree), and allows us to handle arbitrary new datasets, possibly aligned to a different reference, without complications. By preserving the polymerase mutation mechanism in sites with APOBEC3 context (i.e., the first term in the definition of Q_1_), we avoid having to identify only the sites that have not yet mutated (i.e., only the TC and GA sites) and avoid technical difficulties with completely irreversible transition matrices. For simplicity, we also use a simple Jukes-Cantor model with no site rate heterogeneity instead of a more sophisticated GTR model with 4 gamma rate categories. Since Delphy does not yet implement a Skygrid population model, we model the viral population as growing strictly exponentially, as observed in O’Toole et al and Parker et al^5,6^. Finally, by design, we have not implemented a transition between a pre-spillover and post-spillover portion of the tree, as our interest is in public health response to a growing outbreak that is known to be spreading through humans.

### Benchmarks

All benchmarks described below are available in separate folders in the GitHub repo https://github.com/broadinstitute/delphy-2025-paper-data. Each benchmark is organized as a series of numbered scripts. Unless noted otherwise, the scripts are self-contained and will download all external data where necessary. The GitHub repo also includes all the intermediate and final results files for the runs shown in the paper, as well as many of the plots here and in the SI (files that are too large for GitHub or that include GISAID data subject to its data use agreement are available upon request). We intend this repo to be executable documentation of every detail of the benchmarks: while all scripts ran correctly at the time of submission, we do not intend to modify them in the future to ensure they continue to run indefinitely.

We used the following versions of each tool: Delphy 1.0 (build 2036, commit 06a7ee4), MAFFT 7.505 (2022/Apr/10), BEAST2 2.6.2, BEAST X 10.5.0-beta5 Prerelease #1d511b10c2, Sapling 0.1.1 (build 2, commit a0b9da1), BEAGLE commit `3a8d3e6` (Sun Mar 10 2024), IQ-TREE 2.3.6 and TreeTime 0.11.4. Unless otherwise noted, all calculations were done on AWS c7a.2xlarge instances (8 vCPUs). All Delphy runs were performed twice independently, and we checked for agreement and convergence visually using Tracer^56^.

### SARS-CoV-2 data from Lemieux et al 2021 (‘sars-cov-2-lemieux’)

We obtained the accession IDs of the 772 samples that went into making Figure 3A of Ref.^45^ from the authors (‘sample_ids.csv’). Of these, we downloaded the 757 of these that are publicly available in GenBank. We aligned these onto the SARS-CoV-2 reference sequence NC_045512.2 (dated to 2019-12) using mafft^57^ with options --auto and --keeplength. As in^45^, we then masked the initial 267 and final 230 sites.

Delphy was run twice for 2 billion steps, recording trace output every 200,000 steps and a posterior tree sample every 2,000,000 steps. BEAST2 was run using the “equivalent” XML file output by Delphy without any changes; in particular, we used 200 million steps, with the rough heuristic that 10 Delphy steps achieve the work of 1 BEAST2 step. Separate Delphy and BEAST2 runs were prepared with and without site-rate heterogeneity enabled.

MCCs were constructed using TreeAnnotator2^17^ with a 30% burn-in and the option “--heights ca”, so that inner node times are the mean tMRCA of the downstream tips (this coincides with the MCC in Figure 3A of^45^).

An unrooted ML tree was constructed with IQ-Tree 2 with the option “-m HKY+FO” or “-m HKY+FO+G4” to disable or enable site-rate heterogeneity. We then used TreeTime to root it and to estimate the population growth rates, with the options “--coalescent skyline --n-skyline 2 --stochastic-resolve”.

MCCs were plotted using the baltic library, with inner node metadata inferred via simple parsimony on the MCC (ties resolved arbitrarily). ESSs were calculated for all traces using LogAnalyser2^17^ with a burn-in of 30%.

### Zika data from Metsky et al 2017 (‘zika-metsky-2017’)

We extracted the sequences used in^4^ from the BEAST XML file in its Supplementary Data (‘SupplementaryData/BEAST input and output/Phylogenetic analyses and model selection/SRD06-strict-exponential.xml’). These are aligned to the reference sequence KX197192.1, with the initial 107 sites and final 428 sites trimmed. Runs and analysis otherwise follow the same pattern as the SARS-CoV-2 dataset above.

### Ebola data from Gire et al 2014 (‘ebola-gire-2014’)

We extracted the sequences used in ^3^ from the BEAST XML file in Supplementary File 3 of the paper (‘beast/2014_GN.SL_SRD.HKY_strict_ctmc.exp.xml’). In this XML file, the raw sequences are partitioned into the genic and intergenic regions, so that the mapping to the coordinates of the reference KJ660346 sequence is scrambled. We manually deduced the inverse mapping and to reconstitute an MSA aligned to this reference sequence. Runs and analysis otherwise follow the same pattern as the SARS-CoV-2 dataset above.

### Mpox data from O’Toole et al 2023 (‘mpox-otoole-2023’)

We extracted the sequences used in^5^ from one of the BEAST XML file in its companion GitHub repository (‘data/apobec3_2partition.epoch.xml’ in https://github.com/hmpxv/apobec3, commit c0b4c9b). We removed one sample from GISAID (EPI_ISL_13983888), whose sequence is not publicly available, as well as the two pre-spillover sequences dating from before 2017 (KJ642617 from 1971 and KJ642615 from 1978). Except for the GISAID sequence, the remaining 41 sequences form the ‘hMPXV-1’ ingroup in Figure 3C of ^5^. In the XML file, the raw sequences are partitioned into two partitions, one for sites in APOBEC3 context and the other for the remaining sites. Every site of every sequence is marked ‘N’ in at least one partition, so we trivially reconstitute the original aligned sequences.

Delphy was run twice for 200 million steps using the `--v0-mpox-hack` option. Trace output every 20,000 steps and a posterior tree sample every 200,000 steps. As with the earlier benchmarks, MCCs were constructed using TreeAnnotator2^17^ with a 30% burn-in and the option “--heights ca”, so that inner node times are the mean tMRCA of the downstream tips.

### Mpox data from Parker et al 2025 (‘mpox-parker-2025’)

We extracted the sequences used in^6^ from one of the BEAST XML file in its companion GitHub repository (‘BEAST/Mpox_2epoch_combinedDTA.xml.zip’ in https://github.com/andersen-lab/Mpox_West_Africa, commit 2b481da). We removed the three samples from GISAID (EPIISL-13953610, EPIISL-13983888 and EPIISL-15008577), whose sequences are not publicly available, as well as all pre-spillover sequences and sequences dating from before 2017. Except for the GISAID sequence, the remaining 177 sequences form the ‘hMPXV-1’ clade in ^6^. In the XML file, the raw sequences are split over two partitions, one for sites in APOBEC3 context and the other for the remaining sites. Every site of every sequence is marked ‘N’ in at least one partition, so we trivially reconstitute the original aligned sequences.

Delphy was run twice for 1 billion steps using the `--v0-mpox-hack` option. Trace output every 100,000 steps and a posterior tree sample every 1,000,000 steps. As with the earlier benchmarks, MCCs were constructed using TreeAnnotator2^17^ with a 30% burn-in and the option “--heights ca”, so that inner node times are the mean tMRCA of the downstream tips.

For the BEAST comparisons, we modified the original BEAST X XML input file (BEAST/Mpox_2epoch_combinedDTA.xml.zip) to include the same sequences as the Delphy runs, and to remove all configuration relating to phylogeography, detecting the spillover event, and separately modeling evolution before and after the spillover. We also shortened the chain to 50M steps, as this was sufficient to ensure convergence (posterior ESS = 738). The script to modify this input file and the resulting XML file we ran is included in our data repo (mpox-parker-2025-beast.xml).

### SARS-CoV-2 data from GISAID submitted/collected by each CDC week (‘sars-cov-2-gisaid-week-by-week’)

We downloaded from GISAID all (unaligned) SARS-CoV-2 sequences collected on or before 2020-03-31 with accompanying metadata. We filtered out any sequences from non-human hosts, or with length below 20,000 bases, or with uncertain dates. For the results in Figure 6, we included a sequence in a CDC epiweek if its Submission Date preceded the end of that week, and filtered sequences to those submitted in the range 2019-12-01 to 2020-03-28 (end of CDC epiweek 2020-13) and collected on or after 2019-12-01. For the results in Supplementary Figures 7 and 8, we included a sequence in a CDC epiweek if its Collection Date preceded the end of that week, and filtered sequences to those submitted in the range 2019-12-01 to 2024-12-31 and collected on or after 2019-12-01. The sequences passing these filters were grouped into “sequences up to CDC epiweek N”, then aligned to NC_045512.2 using mafft, after which their initial 268 and final 230 sites were masked (as in ^45^). No further masking was applied.

Quick diagnostic runs of sequences up to CDC epiweek N for increasing N rapidly revealed clear outlier sequences (e.g., inducing a tMRCA to early 2019, lying in an isolated long branch with 10s to 100s of mutations, collection dates far preceding reported first cases in a particular region), which we marked for removal in all runs. After several rounds of iterative refinement, no trees contained obvious outliers. We verified that most (but not all) of these offending sequences had been identified as outliers near the beginning of the pandemic, e.g., in ^58^, were marked as “under investigation” in GISAID, or eventually appeared in the exclusion lists of NextStrain or sarscov2phylo. See Supplementary Tables 1 and 2 for a full annotated list of the 94 excluded sequences.

Delphy was run on AWS c7a.4xlarge instances (16 vCPUs) for sequences collected by the end of each CDC epiweek N as above for the SARS-CoV-2 sequences from^45^, with the following options: 5 million steps per tip, a log and tree sampling rate that resulted in 10,000 log samples and 200 tree samples, and a number of threads that was at most 1 per 100 tips and at most 2*16 = 32 in total.

Analysis proceeded as above, except that some trees were so large that TreeAnnotator2 had problems deriving an MCC. Instead, we used the faster MCC calculation used by the Delphy web interface, exposed as a command-line tool `delphy_mcc`, which is hard-coded to use a 30% burn-in and to calculate MCC inner node times as the mean tMRCA of the downstream tips.

### Simulated trees for scaling assessment (‘sims’)

We prepared two groups of 4 SARS-CoV-2-like datasets to assess Delphy’s scalability. Each group contained a simulation with N = 100, 1000, 10,000 and 100,000 tips. In the first group, the simulations used an exponentially growing viral population curve N(t) = n_0_ e^g(t - t0)^, with n_0_ = 6 years, g = 10 / year and t_0_ = 2024-07-31. In the second group, the simulations used a constant-size viral population curve N(t) = n_0_, with n_0_ = 2 years. Given a concrete population curve, we picked N sample times t in the range [2024-01-01, 2024-07-31] distributed according to N(t), mimicking a uniform sampling probability for any member of the viral population. We then executed a standard coalescent simulation^59^ to link these samples into a random ancestry. Next, we picked a random root sequence of 30,000 sites with the state (A, C, G or T) of each site independently sampled from the discrete distribution **π** = [0.30, 0.18, 0.20, 0.32]. We then evolved this random root sequence along the ancestry using the Gillespie algorithm^60^ according to a site-homogeneous HKY model with mutation rate 1×10^-3^ mutations / site / year and a transition-transversion ratio κ = 5. This simulation produces an EMAT with a particular topology and specific mutations at certain times. We recorded this tree in Newick format, a summary of key characteristics of this tree in JSON format, and produced FASTA (only for N <= 1000) and MAPLE files for the dated sequences of all the tips. To perform these large-scale simulations efficiently and output the result as MAPLE files, we wrote a small special-purpose program called *sapling* (see Code and Data Availability).

Delphy was then run twice on a 96-vCPU c7a.24xlarge AWS instance, with the following options: 5 million steps per tip, a log and tree sampling rate that resulted in 10,000 log samples and 200 tree samples, and a number of threads that was at most 1 per 20 tips and at most 2*96 = 192 in total. The runs for N = 100,000 were also run with ∼10,000 cells for discretizing the coalescent prior instead of the default ∼400 (see SI for a comparison of the results when using 625, 1250, 2500 and 5000 cells instead).

For the 100,000-sample runs, we found TreeAnnotator2 unable to extract an MCC, running out of 8GB of memory after 2 hours of runtime. In contrast, `delphy_mcc` completed this task in a few minutes. For consistency, we used `delphy_mcc` to calculate the MCCs of all the runs in the scaling dataset.

### H5N1 in cattle dataset (‘h5n1-andersen-2025’)

We cloned the Andersen lab’s `avian-influenza` data repository from GitHub^61^ at commit ebfdf65 (5 March 2025). This data repository was first used in^47^ and is updated automatically every day. Of the 7011 unique SRRs in the specified commit, we filtered to the 3,069 SRRs whose metadata indicated a cattle host, and further filtered to the 2,850 sequences that had not been retracted. For each of H5N1’s 8 segments, we aligned these 2,850 sequences to the reference genome A/cattle/Texas/24-008749-003/2024 (SRR28752635), as found in `avian-influenza/reference`, using mafft as above. We then concatenated the sequences for all 8 segments, from longest to shortest, to obtain a single linear sequence for each sample (this concatenation assumes no significant reassortment has taken place over the history of all samples, which appears to be the case as of March 2025).

For dating the sequences, we identified the 1,888-sequence subset of the above with corresponding GenBank accessions. While the dates in the SRR metadata are almost all given as a year, sequences deposited in GenBank (often ∼3 months after publication in the SRA) have dates resolved to a single day. A single sequence without a GenBank mapping (SRR29455632) had an exact date in the SRR metadata. We prepared the set of 1,889 cattle-host, nonretracted sequences with exact dates as a FASTA file for input to Delphy. While we also prepared a file for the larger set of 2,850 sequences including those with uncertain dates, runs with these exhibited severe convergence problems and were excluded. For geographic analysis, we used the `geo_loc_name` from GenBank, which is present for all 1,888 GenBank accessions and resolves to a US state in 1,689 samples.

Delphy was run twice for 10 billion steps, with trace output every 1 million steps and a posterior tree sample every 10 million steps. As with the earlier benchmarks, MCCs were constructed using TreeAnnotator2^17^ with a 30% burn-in and the option “--heights ca”, so that inner node times are the mean tMRCA of the downstream tips.

## References

1. Khare, S. et al. GISAID’s Role in Pandemic Response. China CDC Wkly 3, 1049–1051 (2021).

2. GISAID. https://gisaid.org.

3. Gire, S. K. et al. Genomic surveillance elucidates Ebola virus origin and transmission during the 2014 outbreak. Science 345, 1369–1372 (2014).

4. Metsky, H. C. et al. Zika virus evolution and spread in the Americas. Nature 546, 411–415 (2017).

5. O’Toole, Á. et al. APOBEC3 deaminase editing in mpox virus as evidence for sustained human transmission since at least 2016. Science 382, 595–600 (2023).

6. Parker, E. et al. Genomic epidemiology uncovers the timing and origin of the emergence of mpox in humans. medRxiv (2024) doi:10.1101/2024.06.18.24309104.

7. Rambaut, A. et al. A dynamic nomenclature proposal for SARS-CoV-2 lineages to assist genomic epidemiology. Nat Microbiol 5, 1403–1407 (2020).

8. Proposed new B.1 sublineage circulating in India · Issue #38 · cov-lineages/pango-designation. GitHub https://github.com/cov-lineages/pango-designation/issues/38.

9. Kraemer, M. U. G. et al. Spatiotemporal invasion dynamics of SARS-CoV-2 lineage B.1.1.7 emergence. Science 373, 889–895 (2021).

10. McCrone, J. T. et al. Context-specific emergence and growth of the SARS-CoV-2 Delta variant. Nature 610, 154–160 (2022).

11. Viana, R. et al. Rapid epidemic expansion of the SARS-CoV-2 Omicron variant in southern Africa. Nature 603, 679–686 (2022).

12. Didelot, X., Fraser, C., Gardy, J. & Colijn, C. Genomic Infectious Disease Epidemiology in Partially Sampled and Ongoing Outbreaks. Mol Biol Evol 34, 997–1007 (2017).

13. Lau, M. S. Y., Marion, G., Streftaris, G. & Gibson, G. A Systematic Bayesian Integration of Epidemiological and Genetic Data. PLoS Comput. Biol. 11, e1004633 (2015).

14. Klinkenberg, D., Backer, J. A., Didelot, X., Colijn, C. & Wallinga, J. Simultaneous inference of phylogenetic and transmission trees in infectious disease outbreaks. PLoS Comput. Biol. 13, e1005495 (2017).

15. Specht, I. et al. JUNIPER: Reconstructing Transmission Events from Next-Generation Sequencing Data at Scale. medRxiv (2025) doi:10.1101/2025.03.02.25323192.

16. Suchard, M. A. et al. Bayesian phylogenetic and phylodynamic data integration using BEAST 1.10. Virus Evol 4, vey016 (2018).

17. Bouckaert, R. et al. BEAST 2.5: An advanced software platform for Bayesian evolutionary analysis. PLoS Comput Biol 15, e1006650 (2019).

18. Ronquist, F. et al. MrBayes 3.2: efficient Bayesian phylogenetic inference and model choice across a large model space. Syst Biol 61, 539–542 (2012).

19. Rambaut, A. et al. The genomic and epidemiological dynamics of human influenza A virus. Nature 453, 615–619 (2008).

20. Hadfield, J. et al. Nextstrain: real-time tracking of pathogen evolution. Bioinformatics 34, 4121–4123 (2018).

21. McBroome, J. et al. A Daily-Updated Database and Tools for Comprehensive SARS-CoV-2 Mutation-Annotated Trees. Mol Biol Evol 38, 5819–5824 (2021).

22. Turakhia, Y. et al. Ultrafast Sample placement on Existing tRees (UShER) enables real-time phylogenetics for the SARS-CoV-2 pandemic. Nat. Genet. 53, 809–816 (2021).

23. Ye, C. et al. matOptimize: a parallel tree optimization method enables online phylogenetics for SARS-CoV-2. Bioinformatics 38, 3734–3740 (2022).

24. Minh, B. Q. et al. IQ-TREE 2: New Models and Efficient Methods for Phylogenetic Inference in the Genomic Era. Mol. Biol. Evol. 37, 1530–1534 (2020).

25. De Maio, N. et al. Maximum likelihood pandemic-scale phylogenetics. bioRxiv (2022) doi:10.1101/2022.03.22.485312.

26. Ly-Trong, N., Bielow, C., De Maio, N. & Minh, B. Q. CMAPLE: Efficient Phylogenetic Inference in the Pandemic Era. Mol Biol Evol 41, (2024).

27. Sagulenko, P., Puller, V. & Neher, R. A. TreeTime: Maximum-likelihood phylodynamic analysis. Virus Evol 4, vex042 (2018).

28. du Plessis, L. et al. Establishment and lineage dynamics of the SARS-CoV-2 epidemic in the UK. Science 371, 708–712 (2021).

29. Kramer, A. M. et al. Online Phylogenetics with matOptimize Produces Equivalent Trees and is Dramatically More Efficient for Large SARS-CoV-2 Phylogenies than de novo and Maximum-Likelihood Implementations. Syst Biol 72, 1039–1051 (2023).

30. Wertheim, J. O., Steel, M. & Sanderson, M. J. Accuracy in Near-Perfect Virus Phylogenies. Syst Biol 71, 426–438 (2022).

31. Nielsen, R. Mutations as missing data: inferences on the ages and distributions of nonsynonymous and synonymous mutations. Genetics 159, 401–411 (2001).

32. Nielsen, R. Mapping mutations on phylogenies. Syst Biol 51, 729–739 (2002).

33. Hwang, D. G. & Green, P. Bayesian Markov chain Monte Carlo sequence analysis reveals varying neutral substitution patterns in mammalian evolution. Proc Natl Acad Sci U S A 101, 13994–14001 (2004).

34. Lartillot, N. Conjugate Gibbs Sampling for Bayesian Phylogenetic Models. (2007) doi:10.1089/cmb.2006.13.1701.

35. Mateiu, L. & Rannala, B. Inferring complex DNA substitution processes on phylogenies using uniformization and data augmentation. Syst Biol 55, 259–269 (2006).

36. Rodrigue, N., Philippe, H. & Lartillot, N. Uniformization for sampling realizations of Markov processes: applications to Bayesian implementations of codon substitution models. Bioinformatics 24, 56–62 (2008).

37. de Koning, A. P. J., Gu, W. & Pollock, D. D. Rapid likelihood analysis on large phylogenies using partial sampling of substitution histories. Mol Biol Evol 27, 249–265 (2010).

38. de Koning, A. P. J., Gu, W., Castoe, T. A. & Pollock, D. D. Phylogenetics, likelihood, evolution and complexity. Bioinformatics 28, 2989–2990 (2012).

39. Irvahn, J. & Minin, V. N. Phylogenetic stochastic mapping without matrix exponentiation. J Comput Biol 21, 676–690 (2014).

40. Felsenstein, J. Evolutionary trees from DNA sequences: a maximum likelihood approach. J. Mol. Evol. 17, 368–376 (1981).

41. Brooks, S., Gelman, A., Jones, G. & Meng, X.-L. Handbook of Markov Chain Monte Carlo. (CRC Press, 2011).

42. Frenkel, D. & Smit, B. Understanding Molecular Simulation: From Algorithms to Applications. (Elsevier, 2023).

43. Metropolis, N., Rosenbluth, A. W., Rosenbluth, M. N., Teller, A. H. & Teller, E. Equation of state calculations by fast computing machines. J. Chem. Phys. 21, 1087–1092 (1953).

44. Hastings, W. K. Monte Carlo sampling methods using Markov chains and their applications. Biometrika 57, 97–109 (1970).

45. Lemieux, J. E. et al. Phylogenetic analysis of SARS-CoV-2 in Boston highlights the impact of superspreading events. Science 371, (2021).

46. Hasegawa, M., Kishino, H. & Yano, T. Dating of the human-ape splitting by a molecular clock of mitochondrial DNA. J. Mol. Evol. 22, 160–174 (1985).

47. Preliminary report on genomic epidemiology of the 2024 H5N1 influenza A virus outbreak in U.S. cattle (Part 1 of 2). Virological https://virological.org/t/preliminary-report-on-genomic-epidemiology-of-the-2024-h5n1-influenza-a-virus-outbreak-in-u-s-cattle-part-1-of-2/970 (2024).

48. Nguyen, T.-Q. et al. Emergence and interstate spread of highly pathogenic avian influenza A(H5N1) in dairy cattle. bioRxiv 2024.05.01.591751 (2024) doi:10.1101/2024.05.01.591751.

49. Nextstrain: H5N1 Cattle Outbreak. https://nextstrain.org/avian-flu/h5n1-cattle-outbreak/.

50. Tsui, J. L.-H. et al. Genomic assessment of invasion dynamics of SARS-CoV-2 Omicron BA.1. Science 381, 336–343 (2023).

51. Howard-Snyder, W. et al. Densely sampled phylogenies frequently deviate from maximum parsimony in simple and local ways. (2023).

52. Gill, M. S. et al. Improving Bayesian population dynamics inference: a coalescent-based model for multiple loci. Mol Biol Evol 30, 713–724 (2013).

53. Tavaré, S. Some probabilistic and statistical problems in the analysis of DNA sequences. Lecture of mathematics for life science 17, 57 (1986).

54. Yang, Z. Estimating the pattern of nucleotide substitution. J. Mol. Evol. 39, 105–111 (1994).

55. Drummond, A. J., Ho, S. Y. W., Phillips, M. J. & Rambaut, A. Relaxed phylogenetics and dating with confidence. PLoS Biol 4, e88 (2006).

56. Rambaut, A., Drummond, A. J., Xie, D., Baele, G. & Suchard, M. A. Posterior Summarization in Bayesian Phylogenetics Using Tracer 1.7. Syst Biol 67, 901–904 (2018).

57. Katoh, K. & Standley, D. M. MAFFT multiple sequence alignment software version 7: improvements in performance and usability. Mol Biol Evol 30, 772–780 (2013).

58. Temporal signal and the evolutionary rate of 2019 n-CoV using 47 genomes collected by Feb 01 2020. Virological https://virological.org/t/temporal-signal-and-the-evolutionary-rate-of-2019-n-cov-using-47-genomes-collected-by-feb-01-2020/379 (2020).

59. Felsenstein, J. Inferring Phylogenies. (Sinauer Associates Incorporated, 2004).

60. Gillespie, D. T. Exact stochastic simulation of coupled chemical reactions. J. Phys. Chem. 81, 2340–2361 (1977).

61. GitHub - andersen-lab/avian-influenza: Consensus sequences for U.S. H5N1 clade 2.3.4.4b. GitHub https://github.com/andersen-lab/avian-influenza.

