## Supplementary Information for "Delphy: scalable, near-real-time Bayesian phylogenetics for outbreaks"



March 25, 2025


#### 1 Supplementary Figures and Tables

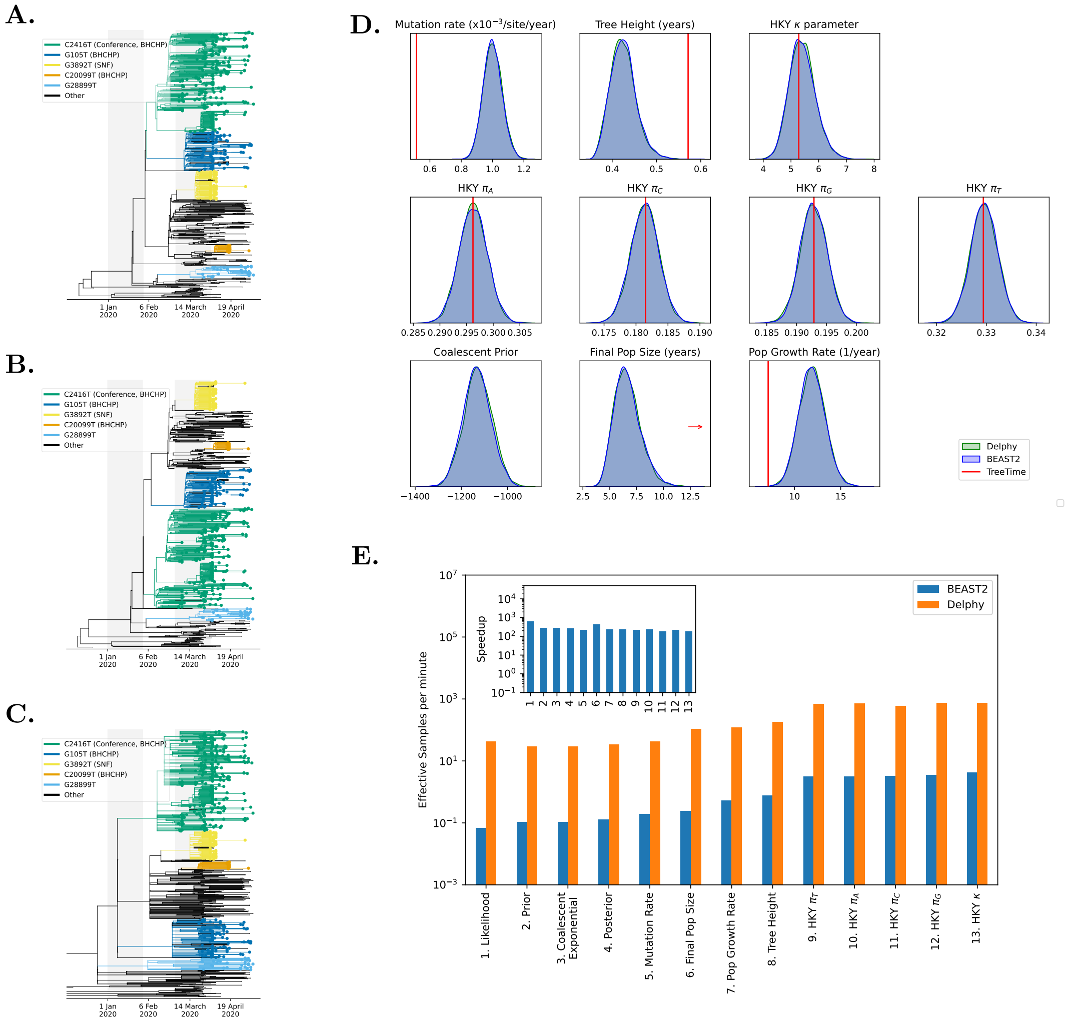


**Supplementary Figure 1: Comparison of Delphy, BEAST2 and ML for the SARS-CoV-2 dataset with no site-rate heterogeneity (compare to Figure 3 in the main text). A.** Maximum-Clade-Credibility (MCC) as produced by Delphy. **B.** Analog for BEAST2. **C.** Maximum-likelihood timed tree, as calculated by IQ-Tree 2 and TreeTime. **D.** Distributions of key observables, as produced by Delphy (green) and BEAST2 (blue), and maximum-likelihood estimate (red). **E.** Runtime efficiency of Delphy vs BEAST2.

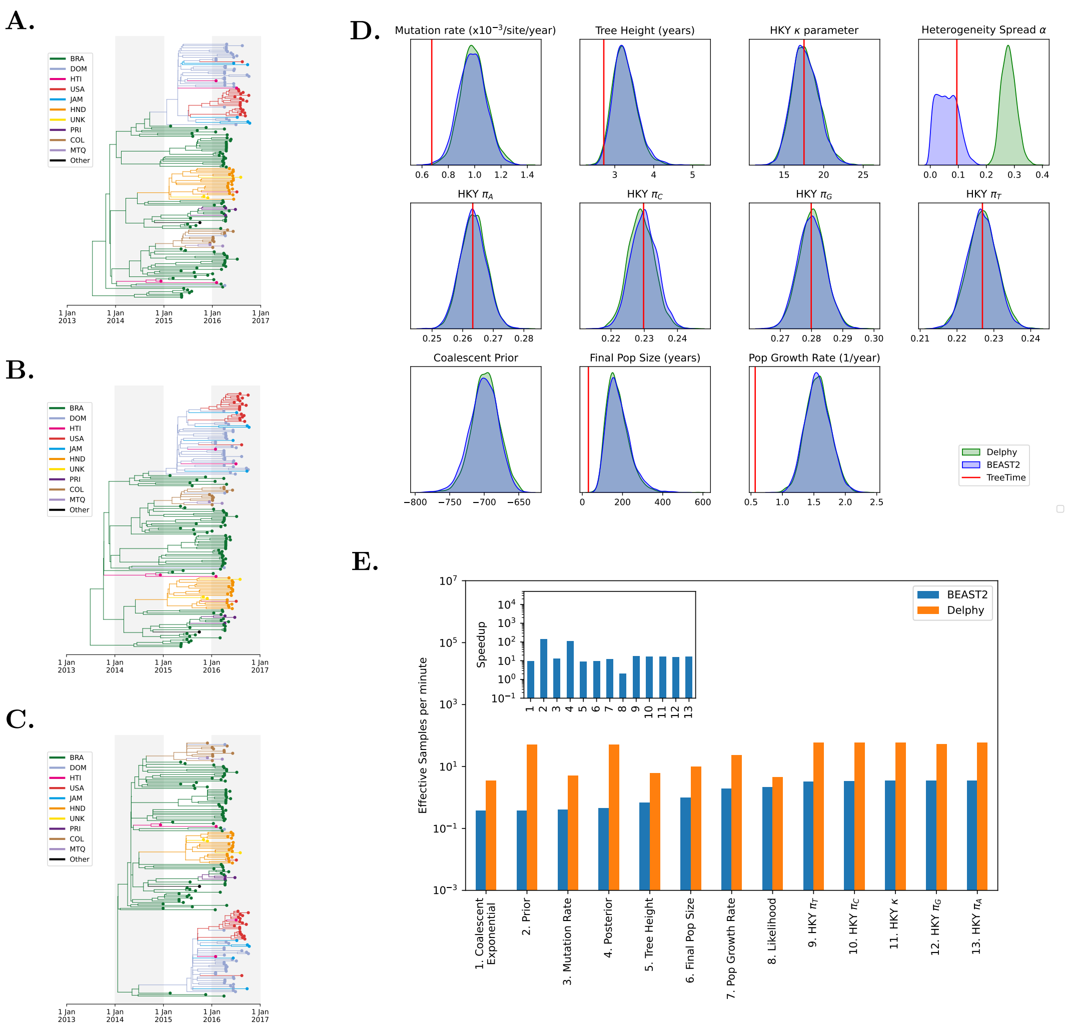


**Supplementary Figure 2: Comparison of Delphy, BEAST2 and ML for the Zika dataset with site-rate heterogeneity (compare to Figure 3 in the main text). A.** Maximum-Clade-Credibility (MCC) as produced by Delphy. **B.** Analog for BEAST2. **C.** Maximum-likelihood timed tree, as calculated by IQ-Tree 2 and TreeTime. **D.** Distributions of key observables, as produced by Delphy (green) and BEAST2 (blue), and maximum-likelihood estimate (red). **E.** Runtime efficiency of Delphy vs BEAST2.

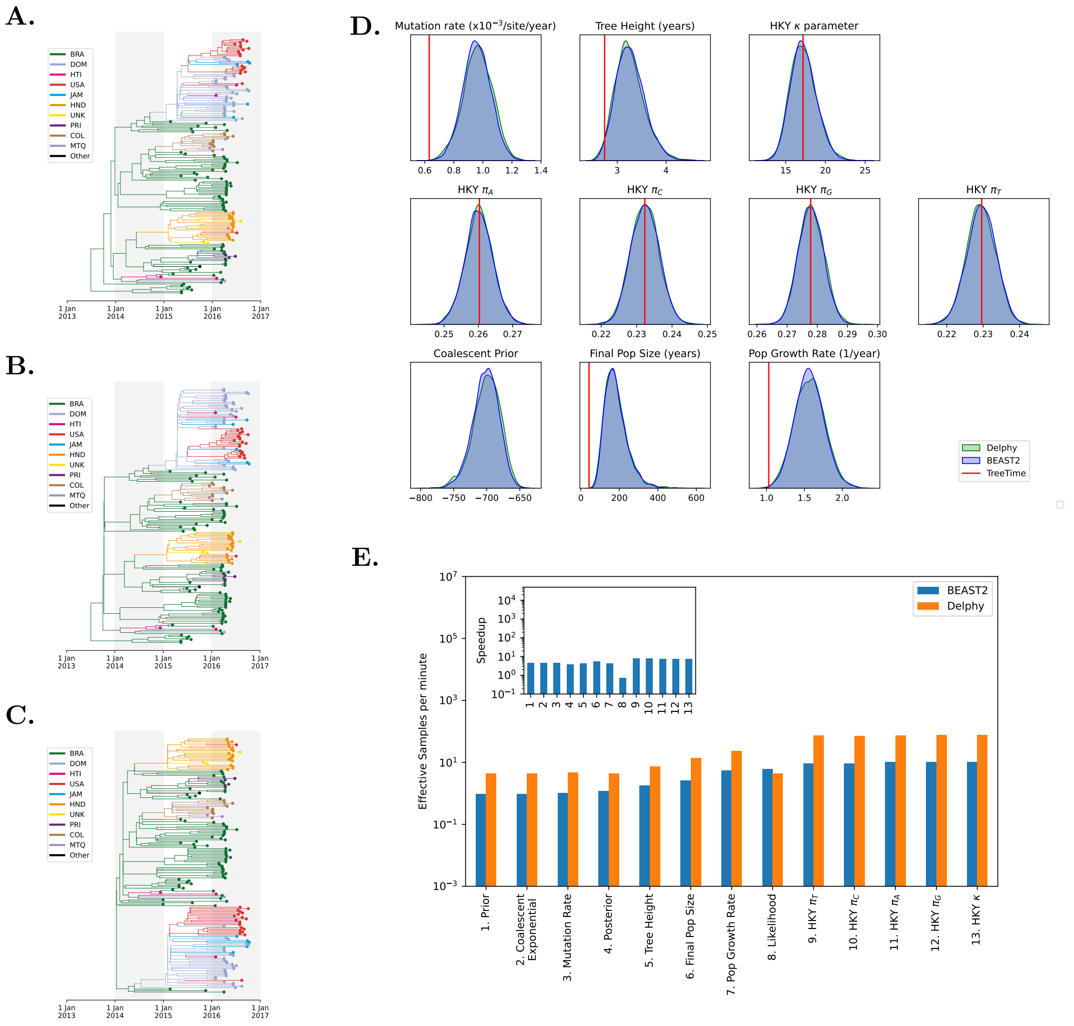


**Supplementary Figure 3: Comparison of Delphy, BEAST2 and ML for the Zika dataset without site-rate heterogeneity (compare to Figure 3 in the main text). A.** Maximum-Clade-Credibility (MCC) as produced by Delphy. **B.** Analog for BEAST2. **C.** Maximum-likelihood timed tree, as calculated by IQ-Tree 2 and TreeTime. **D.** Distributions of key observables, as produced by Delphy (green) and BEAST2 (blue), and maximum-likelihood estimate (red). **E.** Runtime efficiency of Delphy vs BEAST2.

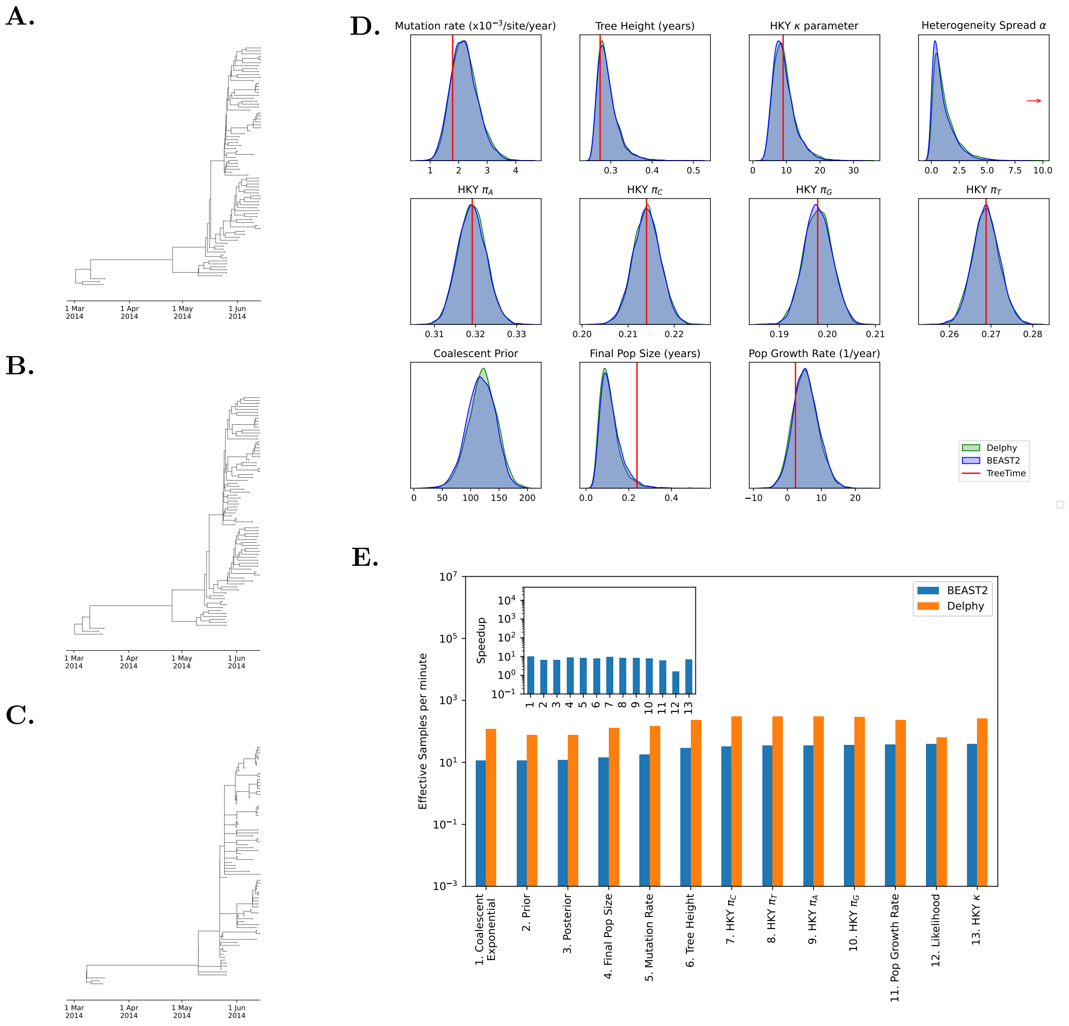


**Supplementary Figure 4: Comparison of Delphy, BEAST2 and ML for the Ebola dataset with site-rate heterogeneity (compare to Figure 3 in the main text). A.** Maximum-Clade-Credibility (MCC) as produced by Delphy. **B.** Analog for BEAST2. **C.** Maximum-likelihood timed tree, as calculated by IQ-Tree 2 and TreeTime. **D.** Distributions of key observables, as produced by Delphy (green) and BEAST2 (blue), and maximum-likelihood estimate (red). **E.** Runtime efficiency of Delphy vs BEAST2.

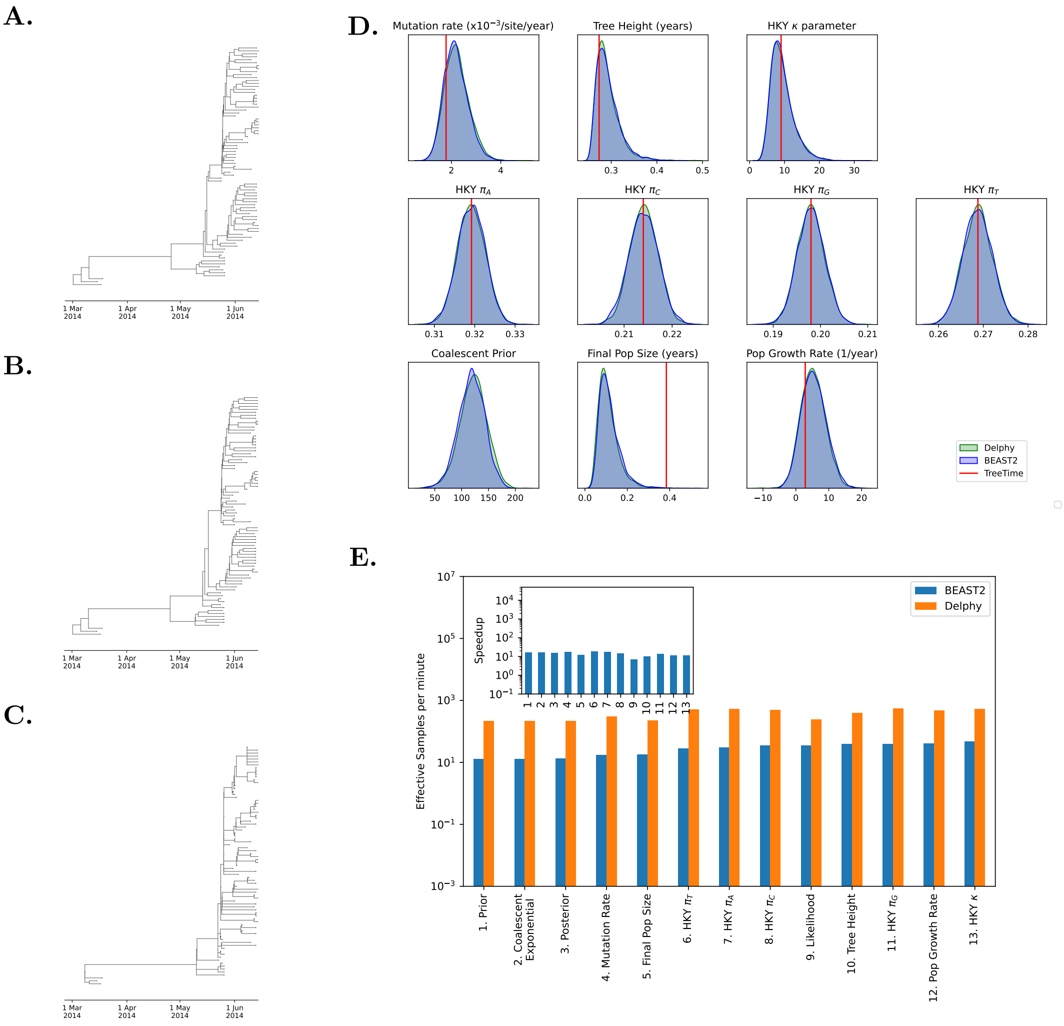


**Supplementary Figure 5: Comparison of Delphy, BEAST2 and ML for the Ebola dataset without site-rate heterogeneity (compare to Figure 3 in the main text). A.** Maximum-Clade-Credibility (MCC) as produced by Delphy. **B.** Analog for BEAST2. **C.** Maximum-likelihood timed tree, as calculated by IQ-Tree 2 and TreeTime. **D.** Distributions of key observables, as produced by Delphy (green) and BEAST2 (blue), and maximum-likelihood estimate (red). **E.** Runtime efficiency of Delphy vs BEAST2.

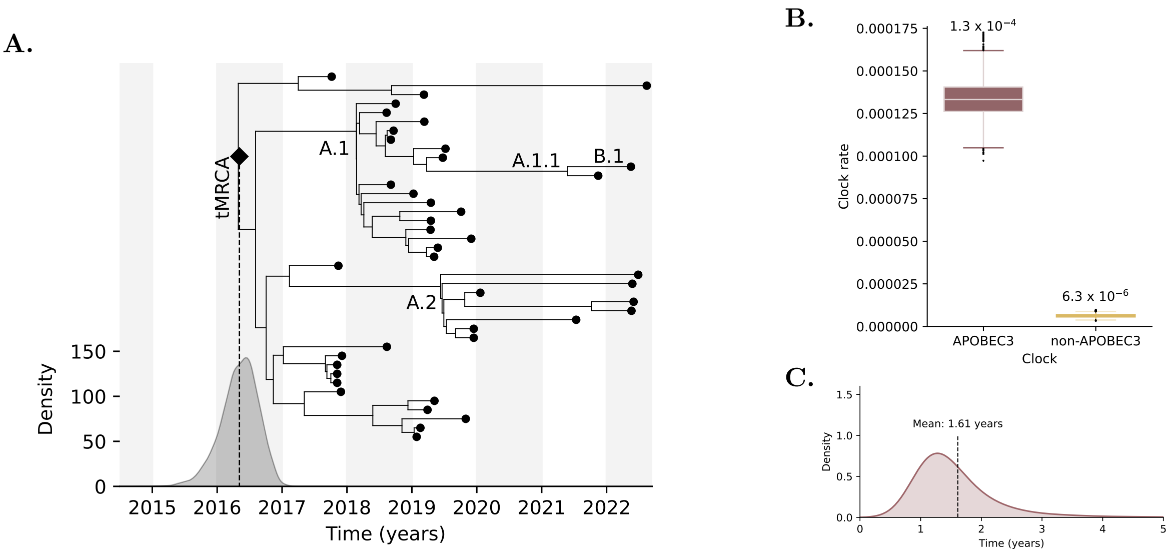


**Supplementary Figure 6: Delphy applied to the mpox dataset from O'Toole et al 2023 [20]. A.** MCC for 41 hMPXV-1 samples from [20] as produced by Delphy (compare to Fig 3C in [20]). **B.** Inferred mutation rates owing to APOBEC3 and non-APOBEC3 mechanisms (compare to Fig S17A of [20], where the APOBEC3 rate is $1.2 \times 10^{-4}$ per site per year and the non-APOBEC3 rate is $4.2 \times 10^{-6}$ per site per year). **C.** Inferred doubling time under exponential growth model (compare to Fig S17B of [20]], with a reported mean of $\sim 1.5$ years).

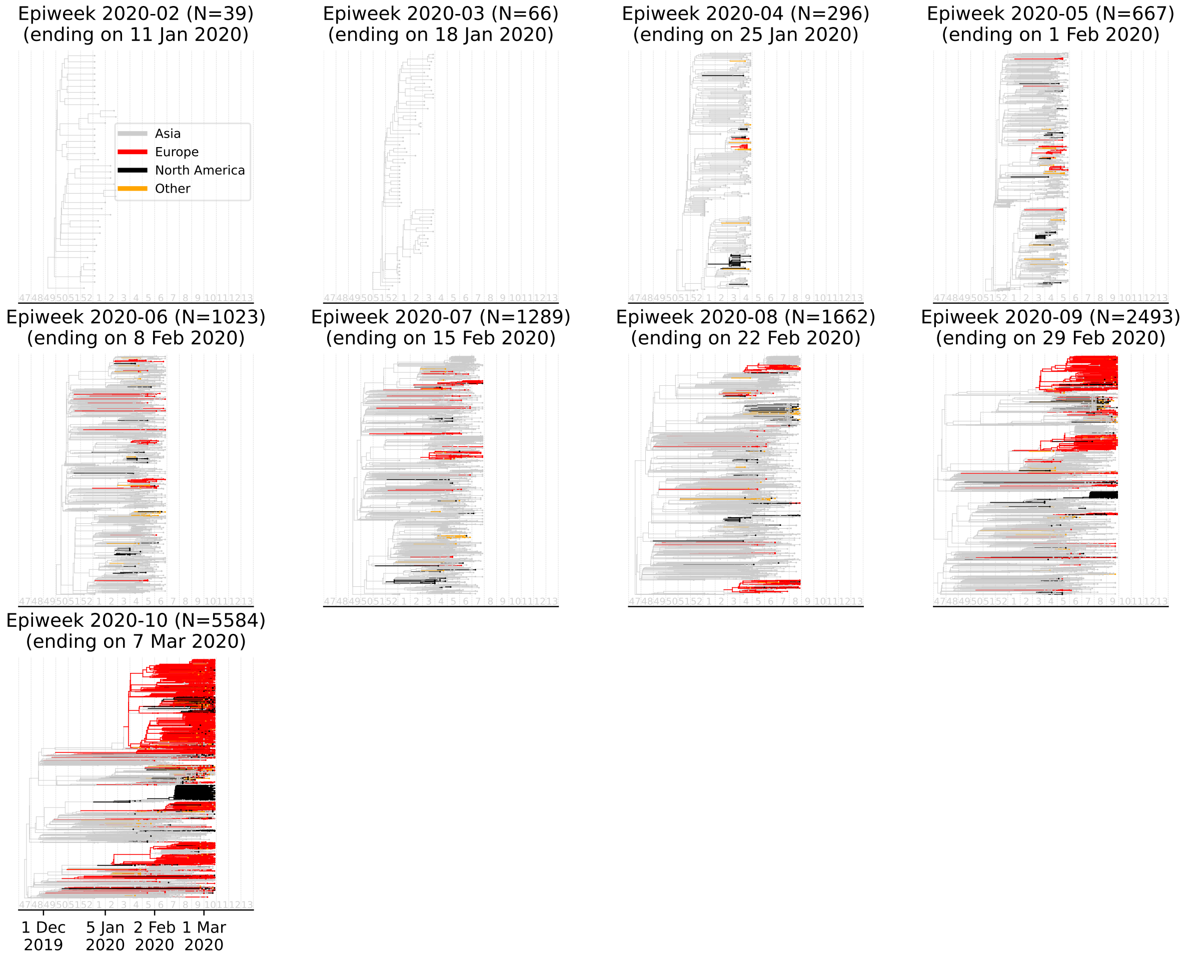


**Supplementary Figure 7: Delphy's view of the first weeks of the COVID-19 pandemic.** Analogous to Figure 5A in the main text, but including all sequences *collected* by the end of each CDC epiweek, not just submitted. Ends at CDC epiweek 202010 since the number of collected sequences explodes beyond that point. We suspect that sequences collected early on but only sequenced much later are of systematically lower quality, as reflected in qualitatively poorer trees and parameter distributions.

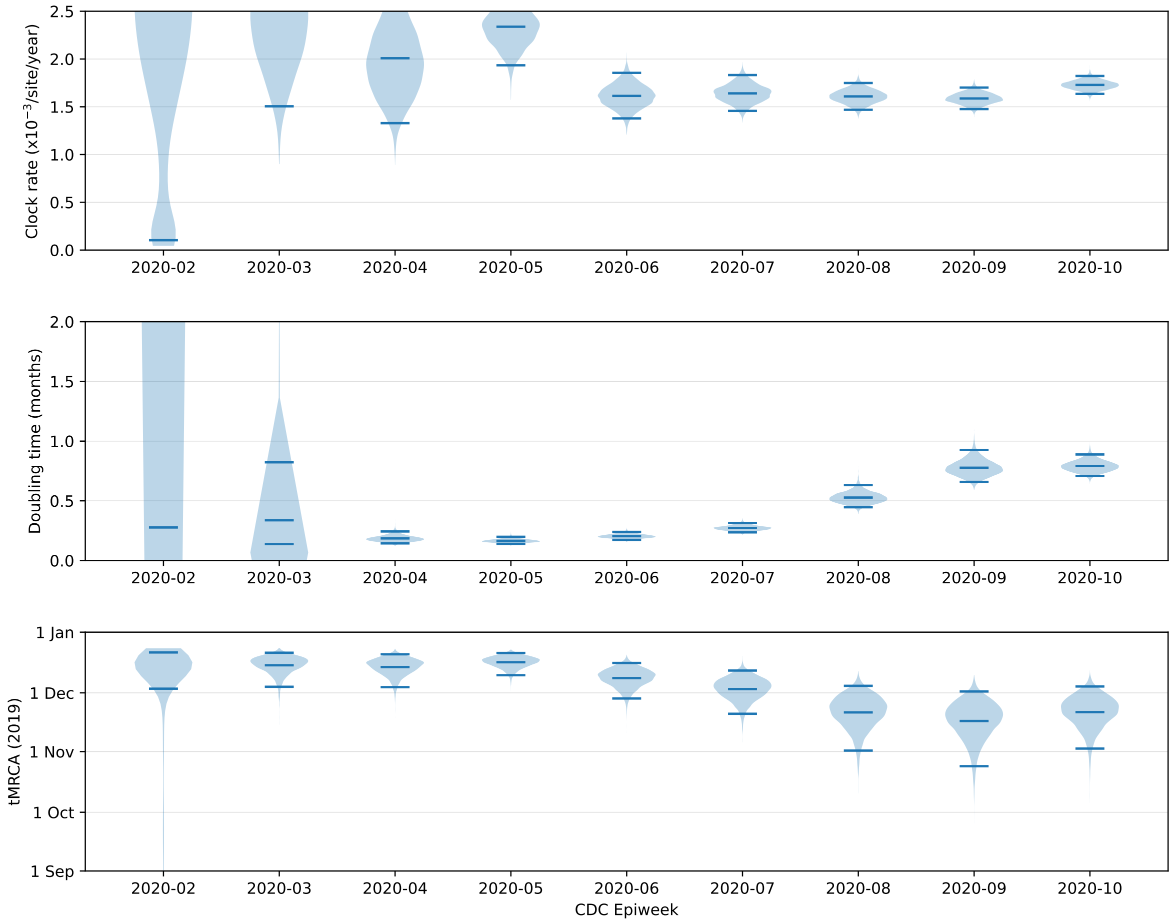


**Supplementary Figure 8: Delphy's view of the first weeks of the COVID-19 pandemic.** Analogous to Figure 5B in the main text, but including all sequences *collected* by the end of each CDC epiweek, not just submitted. Ends at CDC epiweek 202010 since the number of collected sequences explodes beyond that point. We suspect that sequences collected early on but only sequenced much later are of systematically lower quality, as reflected in qualitatively poorer trees and parameter distributions.

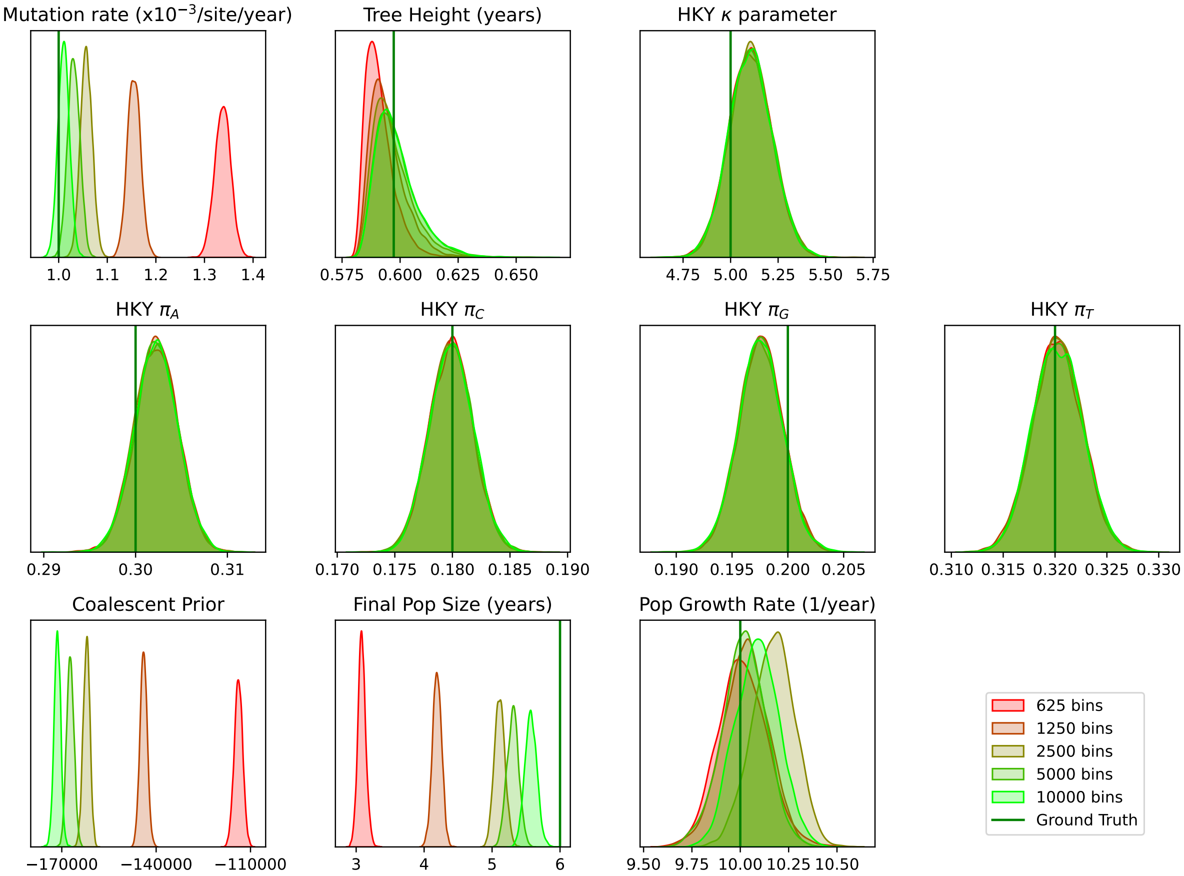


**Supplementary Figure 9: Overall accuracy for $N = 100\,000$ scaling dataset increases as the coalescent prior is discretized more finely.** When sampling intensity is so large that the coalescent prior's $k(t)/N(t)$ factor is comparable to the genome-wide mutation rate $\lambda(x)$, numerical artifacts in the discretized coalescent prior of Eq. (29) have an appreciable effect. By default, Delphy splits the time range of the tree into $\sim 400$ bins to discretize the coalescent prior. Here, we show the effect for the $N = 100\,000$ dataset of using 625, 1250, 2500, 5000 and 10 000 bins. As bin counts increase, the distributions for each observable approach the ground truth values (green vertical bar). In Figure 7 of the main text, the $N = 100\,000$ dataset used 10 000 bins, while all others used the default 400 bins.

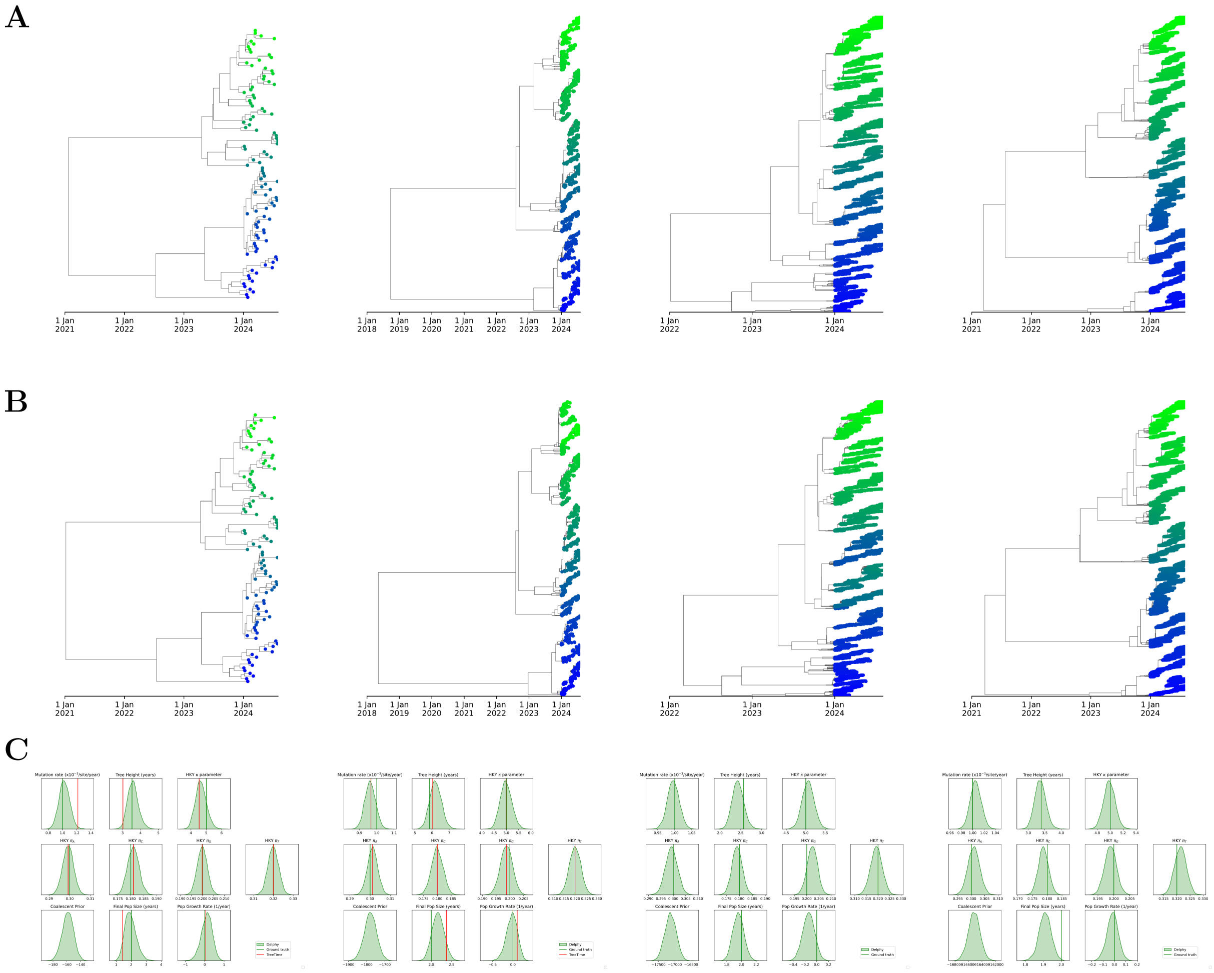


**D**

| | | | | |
|---|---|---|---|---|
| Tips: | 100 | 1000 | 10,000 | 100,000 |
| Steps | 0.5 bn | 5.0 bn | 50.0 bn | 500.0 bn |
| Runtime | 0.09 h | 0.35 h | 1.50 h | 13.42 h |
| Threads | 5 | 50 | 192 | 192 |
| ESS | 634 | 780 | 1365 | 380 |
| ESS / hr | 6860.5 | 2211.9 | 909.0 | 28.3 |

**Supplementary Figure 10: Delphy's scaling behavior, simulations with constant-size viral populations.** Analogous to Figure 7 in the main text, but with a constant effective viral population size of 2 years. As in Figure 7 of the main text, the $N = 100\,000$ dataset used $10\,000$ bins to discretize the coalescent prior as in Eq. (29), while all others used the default 400 bins.

| Accession ID | Name | First anomalous run | Name in Nextstrain | Date excluded in Nextstrain | Notes |
|---|---|---|---|---|---|
| EPI_ISL_406592 | hCoV-19/Guangdong/SZTH-001/2020 | Sub. by CDC-202005 | Shenzhen/SZTH-001/2020 | 2020-02-17 | 1,2,5,6 |
| EPI_ISL_408483 | hCoV-19/Shanghai/IVDC-SH-001/2020 | Sub. by CDC-202007 | Shanghai/IVDC-SH-001/2020 | 2020-02-09 | 2,5,6,7 |
| EPI_ISL_408487 | hCoV-19/Henan/IVDC-HeN-002/2020 | Sub. by CDC-202007 | Henan/IVDC-HeN-002/2020 | 2020-02-09 | 2,5,6,7 |
| EPI_ISL_412900 | hCoV-19/Wuhan/HBCDC-HB-04/2019 | Sub. by CDC-202009 | Wuhan/HBCDC-HB-04/2019 | 2020-06-08 | 2,5,6,7 |
| EPI_ISL_406595 | hCoV-19/Guangdong/SZTH-004/2020 | Sub. by CDC-202010 | Shenzhen/SZTH-004/2020 | 2020-02-17 | 1,2,5,6 |
| EPI_ISL_416030 | hCoV-19/Saudi Arabia/CV1249/2020 | Sub. by CDC-202011 | — | — | 7 |
| EPI_ISL_414935 | hCoV-19/Shandong/LY002-2/2020 | Sub. by CDC-202011 | — | — | 5,7 |
| EPI_ISL_527871 | hCoV-19/Yunnan/KMS1/2020 | Sub. by CDC-202011 | Yunnan/KMS1/2020 | 2021-01-09 | 6,7 |
| EPI_ISL_414934 | hCoV-19/Shandong/LY001-2/2020 | Sub. by CDC-202011 | Shandong/LY001-2/2020 | 2021-04-03 | 5,6,7 |
| EPI_ISL_415593 | hCoV-19/USA/WA-UW65/2020 | Sub. by CDC-202011 | USA/WA-UW65/2020 | 2020-06-08 | 5,6,7 |
| EPI_ISL_417919 | hCoV-19/Malaysia/186197/2020 | Sub. by CDC-202012 | Malaysia/186197/2020 | 2020-06-08 | 5,7 |
| EPI_ISL_417413 | hCoV-19/Turkey/6224-Ankara1034/2020 | Sub. by CDC-202012 | Turkey/6224-Ankara1034/2020 | 2020-06-08 | 5,7 |
| EPI_ISL_8311702 | hCoV-19/Lebanon/LAU-R107/2020 | Col. by CDC-202003 | — | — | 8 |
| EPI_ISL_8311703 | hCoV-19/Lebanon/LAU-R126/2020 | Col. by CDC-202003 | Lebanon/LAU-R126/2020 | 2023-06-09 | 6,8 |
| EPI_ISL_8311704 | hCoV-19/Lebanon/LAU-R128/2020 | Col. by CDC-202003 | Lebanon/LAU-R128/2020 | 2022-09-18 | 6,8 |
| EPI_ISL_8311705 | hCoV-19/Lebanon/LAU-R130/2020 | Col. by CDC-202003 | Lebanon/LAU-R130/2020 | 2022-09-18 | 6,8 |
| EPI_ISL_8311706 | hCoV-19/Lebanon/LAU-R135/2020 | Col. by CDC-202003 | Lebanon/LAU-R135/2020 | 2022-08-23 | 6,8 |
| EPI_ISL_8311707 | hCoV-19/Lebanon/LAU-R137/2020 | Col. by CDC-202003 | Lebanon/LAU-R137/2020 | 2022-09-18 | 6,8 |
| EPI_ISL_8311708 | hCoV-19/Lebanon/LAU-R150/2020 | Col. by CDC-202003 | Lebanon/LAU-R150/2020 | 2022-06-21 | 6,8 |
| EPI_ISL_8311709 | hCoV-19/Lebanon/LAU-R158/2020 | Col. by CDC-202003 | Lebanon/LAU-R158/2020 | 2022-09-18 | 6,8 |
| EPI_ISL_8311711 | hCoV-19/Lebanon/LAU-R178/2020 | Col. by CDC-202003 | Lebanon/LAU-R178/2020 | 2023-06-09 | 6,8 |
| EPI_ISL_8311729 | hCoV-19/Lebanon/LAU-R289/2020 | Col. by CDC-202003 | Lebanon/LAU-R289/2020 | 2022-09-18 | 6,8 |
| EPI_ISL_8311730 | hCoV-19/Lebanon/LAU-R293/2020 | Col. by CDC-202003 | Lebanon/LAU-R293/2020 | 2022-09-18 | 6,8 |
| EPI_ISL_8311731 | hCoV-19/Lebanon/LAU-R294/2020 | Col. by CDC-202003 | Lebanon/LAU-R294/2020 | 2022-10-24 | 6,8 |
| EPI_ISL_8311733 | hCoV-19/Lebanon/LAU-R328/2020 | Col. by CDC-202003 | Lebanon/LAU-R328/2020 | 2022-10-06 | 6,8 |
| EPI_ISL_8311734 | hCoV-19/Lebanon/LAU-R330/2020 | Col. by CDC-202003 | Lebanon/LAU-R330/2020 | 2022-09-18 | 6,8 |
| EPI_ISL_8311741 | hCoV-19/Lebanon/LAU-R350/2020 | Col. by CDC-202003 | Lebanon/LAU-R350/2020 | 2023-06-17 | 6,8 |
| EPI_ISL_8311755 | hCoV-19/Lebanon/LAU-R84/2020 | Col. by CDC-202003 | Lebanon/LAU-R84/2020 | 2022-09-18 | 6,8 |
| EPI_ISL_8311756 | hCoV-19/Lebanon/LAU-R89/2020 | Col. by CDC-202003 | Lebanon/LAU-R89/2020 | 2022-09-18 | 6,8 |
| EPI_ISL_8311758 | hCoV-19/Lebanon/LAU-R97/2020 | Col. by CDC-202003 | Lebanon/LAU-R97/2020 | 2022-09-18 | 6,8 |
| EPI_ISL_8311766 | hCoV-19/Lebanon/LAU-R101/2020 | Col. by CDC-202003 | — | — | 7,8 |
| EPI_ISL_8311767 | hCoV-19/Lebanon/LAU-R146/2020 | Col. by CDC-202003 | — | — | 7,8 |
| EPI_ISL_8311771 | hCoV-19/Lebanon/LAU-R104/2020 | Col. by CDC-202003 | Lebanon/LAU-R104/2020 | 2022-09-18 | 6,7,8 |
| EPI_ISL_8317204 | hCoV-19/Lebanon/LAU-R99/2020 | Col. by CDC-202003 | Lebanon/LAU-R99/2020 | 2023-06-18 | 6,8 |
| EPI_ISL_8317205 | hCoV-19/Lebanon/LAU-R95/2020 | Col. by CDC-202003 | Lebanon/LAU-R95/2020 | 2022-06-03 | 6,7,8 |
| EPI_ISL_8317209 | hCoV-19/Lebanon/LAU-R295/2020 | Col. by CDC-202003 | — | — | 7,8 |
| EPI_ISL_8317210 | hCoV-19/Lebanon/LAU-R351/2020 | Col. by CDC-202003 | Lebanon/LAU-R351/2020 | 2023-06-23 | 6,7,8 |
| EPI_ISL_2671842 | hCoV-19/Japan/20200409-129/2020 | Col. by CDC-202003 | Japan/20200409-129/2020 | 2021-06-25 | 6,7 |
| EPI_ISL_2716627 | hCoV-19/Sierra Leone/SL10/2020 | Col. by CDC-202003 | SierraLeone/SL10/2020 | 2021-06-01 | 6,7 |
| EPI_ISL_2716636 | hCoV-19/Sierra Leone/SL21/2020 | Col. by CDC-202003 | SierraLeone/SL21/2020 | 2021-06-01 | 6,7 |
| EPI_ISL_4405694 | hCoV-19/Argentina/PAIS-A1026/2020 | Col. by CDC-202003 | Argentina/PAIS-A1026/2020 | 2021-10-01 | 6 |
| EPI_ISL_7955525 | hCoV-19/Argentina/PAIS-C0160/2020 | Col. by CDC-202003 | Argentina/PAIS-C0160/2020 | 2023-06-23 | 6,7 |
| EPI_ISL_11575530 | hCoV-19/USA/CO_c8ftxs_0117/2020 | Col. by CDC-202003 | USA/CO_c8ftxs_20200117/2020 | 2023-06-18 | 6 |
| EPI_ISL_14307752 | hCoV-19/USA/MT-UMGC-02796/2020 | Col. by CDC-202003 | — | — | — |
| EPI_ISL_3804266 | hCoV-19/Niger/16250A/2020 | Col. by CDC-202003 | — | — | — |

1 Virological post, "Temporal signal and the evolutionary rate of 2019 n-CoV using 47 genomes collected by Feb 01 2020", from 3 Feb 2020 (https://virological.org/t/temporal-signal-and-the-evolutionary-rate-of-2019-n-cov-using-47-genomes-collected-by-feb-01-2020/379).

2 Bal et al, "Molecular characterization of SARS-CoV-2 in the first COVID-19 cluster in France reveals an amino acid deletion in nsp2 (Asp268del)", 28 Mar 2020 (https://doi.org/10.1016/j.cmi.2020.03.020).

3 Wang et al, "The establishment of reference sequence for SARS-CoV-2 and variation analysis", 13 Mar 2020 (https://doi.org/10.1002/jmv.25762).

4 Lv et al, "Detection of Phenotype-Related Mutations of COVID-19 via the Whole Genomic Data", 8 Jan 2021 (https://doi.org/10.1109/TCBB.2021.3049836).

5 sarscov2phylo exclusion list, available at https://raw.githubusercontent.com/roblanf/sarscov2phylo/master/excluded_sequences.tsv.

6 Nextstrain exclusion list, available at https://raw.githubusercontent.com/nextstrain/ncov/master/defaults/exclude.txt. Quoted dates reflect first commit where accession appears.

7 Marked as "Under Investigation" in GISAID.

8 Cluster of submissions from Lebanon likely incorrectly annotated with collection date 2020-01-10.

**Supplementary Table 1: Excluded GISAID accession IDs for SARS-CoV-2 week-by-week vignette (1 of 2).** After short trial runs for each data subset (submitted/collected by end of CDC epiweek $N$), we visually identified a handful of sequences as clear outliers and excluded them from the analysis. Most of these were also excluded from analyses at the time, are marked "Under Investigation" by GISAID, or are present in the exclusion lists of Nextstrain and/or `sarscov2phylo`.

| Accession ID | Name | First anomalous run | Name in Nextstrain | Date excluded in Nextstrain | Notes |
|---|---|---|---|---|---|
| EPI_ISL_2835566 | hCoV-19/USA/CA-SEARCH-103149/2020 | Col. by CDC-202004 | USA/CA-SEARCH-103149/2020 | 2021-06-09 | 6 |
| EPI_ISL_17116757 | hCoV-19/USA/WV-WV064576/2020 | Col. by CDC-202004 | USA/WV064576/2020 | 2021-09-01 | 6 |
| EPI_ISL_17116755 | hCoV-19/USA/WV-WV064569/2020 | Col. by CDC-202004 | USA/WV064569/2020 | 2021-09-01 | 6,7 |
| EPI_ISL_19045047 | hCoV-19/USA/CA-SEARCH-139716/2020 | Col. by CDC-202004 | — | — | 7 |
| EPI_ISL_19037038 | hCoV-19/Ghana/CRI-01/2020 | Col. by CDC-202004 | — | — | 7 |
| EPI_ISL_19037039 | hCoV-19/Ghana/CRI-02/2020 | Col. by CDC-202004 | — | — | 7 |
| EPI_ISL_19037040 | hCoV-19/Ghana/CRI-03/2020 | Col. by CDC-202004 | — | — | 7 |
| EPI_ISL_19037041 | hCoV-19/Ghana/CRI-04/2020 | Col. by CDC-202004 | — | — | 7 |
| EPI_ISL_19037042 | hCoV-19/Ghana/CRI-05/2020 | Col. by CDC-202004 | — | — | 7 |
| EPI_ISL_19037043 | hCoV-19/Ghana/CRI-06/2020 | Col. by CDC-202004 | — | — | 7 |
| EPI_ISL_19037044 | hCoV-19/Ghana/CRI-08/2020 | Col. by CDC-202004 | — | — | 7 |
| EPI_ISL_19037045 | hCoV-19/Ghana/CRI-09/2020 | Col. by CDC-202004 | — | — | 7 |
| EPI_ISL_19037046 | hCoV-19/Ghana/CRI-10/2020 | Col. by CDC-202004 | — | — | 7 |
| EPI_ISL_19037047 | hCoV-19/Ghana/CRI-11/2020 | Col. by CDC-202004 | — | — | 7 |
| EPI_ISL_19037048 | hCoV-19/Ghana/CRI-13/2020 | Col. by CDC-202004 | — | — | 7 |
| EPI_ISL_19037049 | hCoV-19/Ghana/CRI-14/2020 | Col. by CDC-202004 | — | — | 7 |
| EPI_ISL_19037050 | hCoV-19/Ghana/CRI-15/2020 | Col. by CDC-202004 | — | — | 7 |
| EPI_ISL_19037051 | hCoV-19/Ghana/CRI-16/2020 | Col. by CDC-202004 | — | — | 7 |
| EPI_ISL_17121378 | hCoV-19/USA/OK-PHL-0022663/2020 | Col. by CDC-202005 | USA/OK-PHL-0022663/2020 | 2023-06-18 | 6,7 |
| EPI_ISL_2426018 | hCoV-19/Norway/7651/2020 | Col. by CDC-202005 | Norway/7651/2020 | 2021-06-07 | 6 |
| EPI_ISL_3364539 | hCoV-19/USA/NY-GEO-0231/2020 | Col. by CDC-202005 | — | — | 7 |
| EPI_ISL_17846484 | hCoV-19/USA/un-KDHE-2352575/2020 | Col. by CDC-202006 | — | — | 7 |
| EPI_ISL_1603195 | hCoV-19/Spain/CL-COV00781/2020 | Col. by CDC-202006 | Spain/CL-COV00781/2020 | 2021-04-19 | 6,7 |
| EPI_ISL_4899903 | hCoV-19/Morocco/INH-108/2020 | Col. by CDC-202006 | Morocco/INH-108/2020 | 2021-10-13 | 6 |
| EPI_ISL_4899911 | hCoV-19/Morocco/INH-109/2020 | Col. by CDC-202006 | Morocco/INH-109/2020 | 2021-10-13 | 6 |
| EPI_ISL_4899898 | hCoV-19/Morocco/INH-107/2020 | Col. by CDC-202006 | Morocco/INH-107/2020 | 2021-10-13 | 6 |
| EPI_ISL_4899870 | hCoV-19/Morocco/INH-103/2020 | Col. by CDC-202006 | Morocco/INH-103/2020 | 2022-09-18 | 6 |
| EPI_ISL_4899881 | hCoV-19/Morocco/INH-104/2020 | Col. by CDC-202006 | Morocco/INH-104/2020 | 2021-10-19 | 6 |
| EPI_ISL_4899863 | hCoV-19/Morocco/INH-101/2020 | Col. by CDC-202006 | Morocco/INH-101/2020 | 2021-10-13 | 6 |
| EPI_ISL_4899888 | hCoV-19/Morocco/INH-105/2020 | Col. by CDC-202006 | Morocco/INH-105/2020 | 2021-10-14 | 6 |
| EPI_ISL_4899917 | hCoV-19/Morocco/INH-MN908947/2020 | Col. by CDC-202006 | Morocco/INH-MN908947/2020 | 2022-10-06 | 6 |
| EPI_ISL_4899892 | hCoV-19/Morocco/INH-106/2020 | Col. by CDC-202006 | — | — | — |
| EPI_ISL_1263332 | hCoV-19/Belgium/UZA-UA-CV0615326772/2020 | Col. by CDC-202006 | Belgium/UZA-UA-CV0615326772/2020 | 2021-03-18 | 6 |
| EPI_ISL_1265909 | hCoV-19/USA/OH-ODH-SC1040172/2020 | Col. by CDC-202007 | USA/OH-ODH-SC1040172/2020 | 2021-03-19 | 6,7 |
| EPI_ISL_1014733 | hCoV-19/Spain/MD-IBV-99018532/2020 | Col. by CDC-202008 | Spain/MD-IBV-99018532/2020 | 2021-02-18 | 6 |
| EPI_ISL_1311841 | hCoV-19/Netherlands/ZH-EMC-2080/2020 | Col. by CDC-202008 | Netherlands/ZH-EMC-2080/2020 | 2021-03-24 | 6 |
| EPI_ISL_1311840 | hCoV-19/Netherlands/ZH-EMC-2079/2020 | Col. by CDC-202008 | Netherlands/ZH-EMC-2079/2020 | 2021-03-24 | 6 |
| EPI_ISL_10980369 | hCoV-19/Zambia/MH23_107_4328/2020 | Col. by CDC-202008 | — | — | — |
| EPI_ISL_1136974 | hCoV-19/USA/NV-NSPHL-338695/2020 | Col. by CDC-202008 | USA/NV-NSPHL-338695/2020 | 2021-03-04 | 6 |
| EPI_ISL_9879582 | hCoV-19/Mongolia/UB-109429/2020 | Col. by CDC-202008 | Mongolia/UB-109429/2020 | 2022-04-30 | 6,7 |
| EPI_ISL_1167830 | hCoV-19/Chile/AR-265171/2020 | Col. by CDC-202008 | Chile/AR-265171/2020 | 2021-03-08 | 6 |
| EPI_ISL_7946128 | hCoV-19/USA/CA-SEARCH-58338/2020 | Col. by CDC-202008 | — | — | — |
| EPI_ISL_7946211 | hCoV-19/USA/CA-SEARCH-58362/2020 | Col. by CDC-202008 | — | — | — |
| EPI_ISL_7946167 | hCoV-19/USA/CA-SEARCH-58348/2020 | Col. by CDC-202008 | — | — | |
| EPI_ISL_10980370 | hCoV-19/Zambia/MH21_105_5506/2020 | Col. by CDC-202009 | — | — | 7 |
| EPI_ISL_417446 | hCoV-19/Italy/LOM-UniMI02/2020 | Col. by CDC-202009 | Italy/UniMI02/2020 | 2020-06-08 | 6,7 |
| EPI_ISL_2758215 | hCoV-19/India/un-IRSHA-CD210871/2020 | Col. by CDC-202010 | India/un-IRSHA-CD210871/2020 | 2021-06-05 | 6,7 |
| EPI_ISL_2758214 | hCoV-19/India/un-IRSHA-CD210927/2020 | Col. by CDC-202010 | India/un-IRSHA-CD210927/2020 | 2021-06-05 | 6,7 |
| EPI_ISL_2758213 | hCoV-19/India/un-IRSHA-CD210929/2020 | Col. by CDC-202010 | India/un-IRSHA-CD210929/2020 | 2021-06-05 | 6,7 |

1 Virological post, "Temporal signal and the evolutionary rate of 2019 n-CoV using 47 genomes collected by Feb 01 2020", from 3 Feb 2020 (https://virological.org/t/temporal-signal-and-the-evolutionary-rate-of-2019-n-cov-using-47-genomes-collected-by-feb-01-2020/379).

2 Bal et al, "Molecular characterization of SARS-CoV-2 in the first COVID-19 cluster in France reveals an amino acid deletion in nsp2 (Asp268del)", 28 Mar 2020 (https://doi.org/10.1016/j.cmi.2020.03.020).

3 Wang et al, "The establishment of reference sequence for SARS-CoV-2 and variation analysis", 13 Mar 2020 (https://doi.org/10.1002/jmv.25762).

4 Lv et al, "Detection of Phenotype-Related Mutations of COVID-19 via the Whole Genomic Data", 8 Jan 2021 (https://doi.org/10.1109/TCBB.2021.3049836).

5 sarscov2phylo exclusion list, available at https://raw.githubusercontent.com/roblanf/sarscov2phylo/master/excluded_sequences.tsv.

6 Nextstrain exclusion list, available at https://raw.githubusercontent.com/nextstrain/ncov/master/defaults/exclude.txt. Quoted dates reflect first commit where accession appears.

7 Marked as "Under Investigation" in GISAID.

8 Cluster of submissions from Lebanon likely incorrectly annotated with collection date 2020-01-10.

**Supplementary Table 2: Excluded GISAID accession IDs for SARS-CoV-2 week-by-week vignette (2 of 2).** Continued from previous figure.

#### 2 Explicit Mutation-Annotated Trees

Inspired by the Mutation-Annotated Trees (MATs) introduced by UShER [23] and matOptimize [27] to build parsimonious phylogenetic trees, and building on ideas first presented by Nielsen [19], we introduce *Explicit Mutation-Annotated Trees (EMATs)* (see Figure 11). Informally, an EMAT is a timed bifurcating tree with timed decorations on its branches where the sequence changes (*mutations*) or becomes ambiguous (*missations*) at a specific site, together with a reference sequence at a point above the root. Each leaf (tip) represents a dated sample with a known sequence, although that sequence may have missing data and/or gaps. As usual, all sequences are assumed to be aligned onto an $L$-site reference, and the sequences specify for each site $\ell$ ranging from 1 to $L$ one of: (a) the state (`A`, `C`, `G`, `T`) at that site; or (b) the fact that that site is in an alignment gap ('`-`') or was not (reliably) sequenced ('`N`'). As usual, we do not differentiate between gaps and missing data. We also do not distinguish ambiguous base calls (e.g., `Y` = `C` or `T`) and missing sites (`N`); in practice, this is not a significant limitation. Each inner node represents the most recent common ancestor of its children. Every other point in the tree represents some species along the lineage that connects a node to its parent. As a result, every point in an EMAT has a specific sequence at a subset of the sites in the genome. Mutations and missations encode these sequences and the relevant subsets of sites efficiently, analogously to how mutations in UShER's MATs encode the sequence of nodes in a parsimony tree. EMATs extend MATs by assigning times to both nodes and mutations (so in principle, there may be multiple mutations at the same site along a single branch). Further, missations permit treating missing data efficiently without imputing it; an EMAT with missations represents an appropriately weighted collection of corresponding EMATs where those missations have been replaced by all possible downstream mutations (see Section 5).

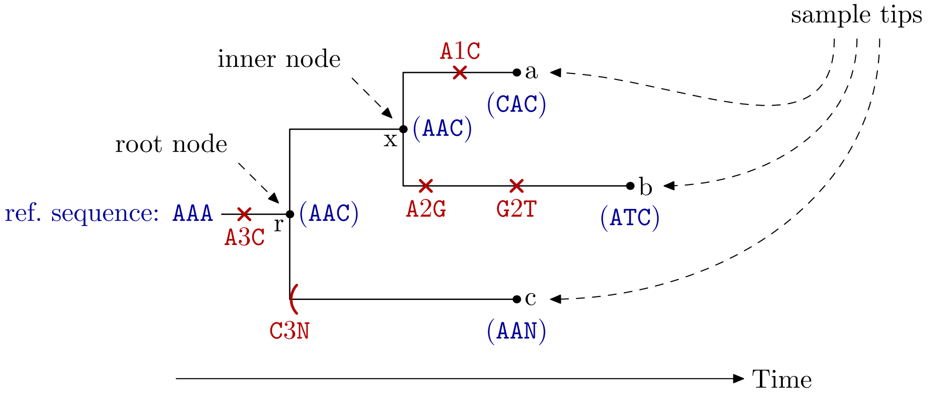


**Supplementary Figure 11:** An Explicit Mutation-Annotated Tree (EMAT). The nodes $a$, $b$ and $c$ are dated tips, whose sequences were determined experimentally. The node $x$ is an inner node, the most recent common ancestor of $a$ and $b$. The root node $r$ is the most recent common ancestor of all the tips. An EMAT explicitly encodes the reference sequence above the root, the mutation events (red crosses) and the missations (red parentheses). From this representation, the sequence of the species represented at any point in the tree is determined implicitly (shown in parentheses in blue below every node).

Formally, an EMAT $\mathcal{T}$ for $N$ dated input sequences is a timed binary tree with $N$ tips (leaves) and $N-1$ inner nodes. A node is indexed by $i$ ranging from 1 to $2N-1$. For notational convenience, we list tips first, so tip $i$ corresponds to input sequence $i$. The ancestor of node $i$ is denoted by $\varphi(i)$. The root node, with index $r$, has no ancestor. However, it is frequently useful to refer to the point *above the root*, which we denote by $\perp$. Formally, we set $\varphi(r) = \perp$. Each node has a time $t_i$ that grows towards the future. Times must be consistent with the tree topology, so $t_{\varphi(i)} < t_i$ for all $i$. Tip times correspond to the collection times of the associated samples. Presently, these must be precise instants in time, though the extension to allow tip times to sample an input time range, as with BEAST's "tip date sampling" feature, is straightforward.

Every node except the root node is the endpoint of a branch from $\varphi(i)$ to $i$, which we call *branch i*. A point $x$ on a tree is uniquely specified by the branch $i$ that it is on and a time $t$ in the interval $(t_{\varphi(i)}, t_i]$. When we need to give explicit coordinates for a point on the tree, we use the notation $[[i,t]]$; when referring to the point at the node $i$ itself, we often simply use $i$ in place of $[[i,t_i]]$. We denote by $\tau_i$ the length of branch $i$, given by $\tau_i = t_i - t_{\varphi(i)}$. The set of all points on branch $i$ is denoted by $B_i$. For completeness, we define the root branch $B_r$ to include the single point at the root, with coordinates $[[r,t_r]]$. The set of all points on the tree is $\mathcal{T} := \cup_i B_i$. We distinguish this set from the tree $\mathcal{T}$ itself only when context does not make the distinction clear.

Every branch $i$ is decorated with a set of mutations $\mathcal{M}_i$. The set of all mutations on the tree is $\mathcal{M} := \cup_i \mathcal{M}_i$. A mutation $m$ is a tuple $(i,t,a,\ell,b)$, which specifies a change in the state of site $\ell$ from $a$ to $b$ at $[[i,t]]$. In the discussions below, we omit some members of this tuple when they are clear from context or they are irrelevant. In diagrams, we represent mutations as crosses at $[[i,t]]$ labeled $a\ell b$, e.g., `A123C`.

The sequence at $x$ is denoted by the states $s_x^{(\ell)}$ at every site $\ell$. The sequence above the root, denoted by $s_\perp^{(\ell)}$, is an (arbitrary) reference sequence. We can reconstruct $s_x^{(\ell)}$ by successively applying to $s_\perp^{(\ell)}$ the mutations on the path from $\perp$ to $x$ in order of encounter. Note that $\mathcal{M}_r$, the mutations on the *root branch* are special: they simply detail the differences between the reference sequence and the sequence at the root, and their times are meaningless. Separating the reference and root sequences simplifies the implementation of tree moves that change the root, as described below. While we represent $s_\perp^{(\ell)}$ explicitly, we only reconstruct portions of $s_x^{(\ell)}$ as above when needed.

Each branch is also decorated at its start with "missations" $\Phi_i$, in analogy with mutations. The set of all missations on the tree is $\Phi := \cup_i \Phi_i$. A missation $\phi$ is a tuple $(i,a,\ell)$ that signals that every species below the starting point of branch $i$, including all downstream tips, are missing data at site $\ell$, whereas in the species immediately ancestral to the missation, the state of site $\ell$ is $a$. As with mutations, we omit below some members of this tuple when they are clear from context. In diagrams, we represent missations as a parenthesis ("(") labeled $a\ell$`N`, e.g., `A123N`.

Given an arbitrary point $x$ on the tree, we define $\xi(x)$ as the set of sites for which at least one tip downstream of $x$ is informative. We use $\xi(x)$ to implement our strategy for dealing with missing data, which we call "`N`-pruning" (Section 5). Missations efficiently encode the value of $\xi(x)$ in terms of deltas just before the start of each branch. To calculate $\xi(x)$, we start with the set $\{1,\ldots,L\}$ and successively remove any sites $\ell$ for which there is a missation at $\ell$ on the path from $\perp$ to $x$. Note that missations on the root branch signal sites for which none of the tips are informative. By construction, for a given set of sequences and tree topology, there is a unique way to decorate the tree with missations: $\xi(x)$ is defined in terms of $x$ and its downstream tips, and the missations at branch $i$ simply encode differences between $\xi(\varphi(i)-)$ and $\xi(\varphi(i)+)$ immediately above and below the starting point of branch $i$. This property is essential for correct sampling: each possible way of imputing all the missing data is thus associated with exactly one and only one EMAT. Inspired by MAPLE [12], we encode missations as a sorted list of disjoint contiguous ranges of missing sites (since such groupings are very common in real sequencing data) and the ancestral state is encoded only in terms of its difference from the reference sequence. This representation remains compact even in the face of long sequences with many large gaps (e.g., as is typical in mpox sequences).

In light of the success of UShER, it should be clear that the above representation allows for a very efficient encoding of the sequences in a genomic epidemiology dataset, including the portions of the sequences that are missing. As detailed below, it also allows for very efficient calculations and Monte Carlo moves.

To conclude this section, we introduce notation to integrate over branches and trees. We define the integral over the points $B_i$ of a branch $i$ in terms of the time coordinate, as follows:

$$\int_{x\in B_i} f(x)\,dx := \int_{t_{\varphi(i)}}^{t_i} f([[i,t]])\,dt.$$

Similarly, we define an integral over the tree by summing over its branches:

$$\int_{x\in\mathcal{T}} f(x)\,dx := \sum_i \int_{x\in B_i} f(x)\,dx.$$

Integrals over subtrees, including paths between points on a tree, are defined similarly.

As a trivial example, the length of branch $i$ can be calculated as follows:

$$\tau_i = \int_{x \in B_i} dx = t_i - t_{\varphi i}.$$

Analogously, the total branch length $T$ of the tree can be calculated as:

$$T = \int_{x \in \mathcal{T}} dx.$$

### 3 Posterior distribution

The posterior distribution that we sample is, up to normalization, as follows:

$$\mathrm{P}(\mathcal{T}, \boldsymbol{\theta}) \propto \mathrm{L}(\mathcal{T}) \cdot \mathcal{G}(\mathcal{T}|\boldsymbol{\theta}) \cdot \pi_{\mathrm{anc}}(\mathcal{T}|\boldsymbol{\theta}) \cdot \pi_{\boldsymbol{\theta}}(\boldsymbol{\theta}). \tag{1}$$

The four factors are:

- A *parameter prior* $\pi_{\boldsymbol{\theta}}(\boldsymbol{\theta})$ that is a product of priors on associated inference parameters $\boldsymbol{\theta}$, such as the mutation rate $\mu$.
- An *ancestry prior* $\pi_{\mathrm{anc}}(\mathcal{T}|\boldsymbol{\theta})$ on the genealogy of tree $\mathcal{T}$. We have implemented the standard neutral coalescent prior (adapted to be parallelizable as described in Section 9), but other priors are possible (e.g., a Yule birth-death prior).
- A *genetic prior* $\mathcal{G}(\mathcal{T}|\boldsymbol{\theta})$ for the exact root sequence and mutations layered onto the genealogy of $\mathcal{T}$. In the traditional Bayesian framework, this term forms the bulk of the *tree likelihood*, calculated via Felsenstein pruning [4]. Here, the *model* instead describes the exact genetic sequences at every point of the tree, so the terms of the usual tree likelihood appear in a prior.
- The *likelihood* $\mathrm{L}(\mathcal{T})$ of the *data* given the tree $\mathcal{T}$.

We discuss each of these four factors in more detail below.

#### 3.1 Likelihood

Given that $\mathcal{T}$ fully specifies the sequences of the tips except where there is missing data, the likelihood is either 1 if $\mathcal{T}$ agrees with the data, or 0 otherwise. In a more elaborate treatment, the likelihood would be a product of weight vectors at every tip and site, to model, say, sequencing errors; we leave that extension for future work. By choosing our starting tree and MCMC moves to always preserve the compatibility of the model with the experimental data, we avoid ever having to calculate the likelihood explicitly.

#### 3.2 Genetic prior

As usual, we assume that each site $\ell$ evolves independently according to a continuous-time Markov chain with transition rate matrix $Q^{(\ell)}_{ab}$. For convenience, we denote by $Q^{(\ell)}_a$ the negative of the diagonal elements, so that $Q^{(\ell)}_a := -Q^{(\ell)}_{aa}$. We denote by $\pi^{(\ell)}_a$ the stationary distribution of this Markov chain at site $\ell$, which satisfies $\sum_a \pi^{(\ell)}_a Q^{(\ell)}_{ab} = 0$.

The *total mutation rate at* $x$, denoted by $\lambda(x)$, is the rate at which mutations happen at point $x$ on the tree on any site in $\xi(x)$. It is given by

$$\lambda(x) := \sum_{\ell \in \xi(x)} Q^{(\ell)}_{s^{(\ell)}_x}. \tag{2}$$

As discussed in Section 5, the restriction to sites in $\xi(x)$ arises from allowing all possible genetic histories below any given missation.

A central result of this paper, which extends Nielsen's original formulation [19], is that the genetic prior $\mathcal{G}(\mathcal{T}|\boldsymbol{\theta})$ has the following form:

$$\mathcal{G}(\mathcal{T}|\boldsymbol{\theta}) = \left\{ \prod_{\ell \in \xi(r)} \pi^{(\ell)}_{s_r^{(\ell)}} \right\} \cdot \exp\left[ -\int_{x\in\mathcal{T}} \lambda(x)\, dx \right] \cdot \prod_{(a,\ell,b)\in\mathcal{M}} Q^{(\ell)}_{ab}. \tag{3}$$

This is simply the probability of the particular realizations of the jump chains for the continuous-time Markov processes that model evolution at each site. By contrast, the traditional tree likelihood $\mathcal{L}_{\text{tree}}(\mathcal{T}|\boldsymbol{\theta})$, as calculated by Felsenstein pruning and used in BEAST and other similar Bayesian phylogenetics tools, encodes the probability of observing particular states at every node, then sums over all possible values of those states. The equivalence of these two approaches is intuitive, and is indeed a standard result of continuous-time Markov processes. However, for readers unfamiliar with this result, a demonstration of the equivalence using the present notation is given in Section 4. A consequence of the equivalence of these two approaches is that the results from a Delphy run are statistically indistinguishable from an *equivalent* BEAST run.

The crucial property of Eq. (3) is that it is a straight product of simple factors, and the overwhelming majority of those factors cancel out when taking ratios of genetic priors of trees that only differ in a localized region. This observation implies that: (a) local MCMC moves on EMATs can be implemented very efficiently, in time proportional to the number of mutations and missation blocks in nearby branches, not the number of variable sites in the entire dataset (a fact not exploited in Nielsen's original formulation); and (b) local moves acting on different parts of the tree are completely decoupled, and may thus be attempted in parallel. Clearly the EMAT representation trades off some statistical efficiency for substantial mathematical simplifications. However, particularly in genomic epidemiology datasets, where $N$ sample sequences can generally be placed into a parsimony tree involving only $\sim N$ mutations [9, 27], this is a very favorable trade-off.

To establish intuition for the behavior of $\mathcal{G}(\mathcal{T}|\boldsymbol{\theta})$, we consider trees with no missing data and specialize to a Jukes-Cantor model of evolution with no site rate heterogeneity, whereby all possible mutations occur at the same rate $\mu/3$ at every site. In that scenario, the above expression takes the following trivial form:

$$\mathcal{G}(\mathcal{T}|\boldsymbol{\theta}) = \left(\frac{1}{4}\right)^L \cdot e^{-\mu L T} \cdot \left(\frac{\mu}{3}\right)^M, \qquad \text{(Jukes-Cantor model, no site-rate heterogeneity)}, \tag{4}$$

where $T$ is the total branch length of the tree and $M$ is the total number of mutations on it. We often use the intuition built on the Jukes-Cantor model to make good MCMC proposals, but then rely on the general expression for $\mathcal{G}(\mathcal{T}|\boldsymbol{\theta})$ in the Metropolis-Hastings ratio to obtain correct samples for the actual evolution model we use.

For some calculations, it is advantageous to organize the calculation of the genetic prior $\mathcal{G}(\mathcal{T}|\boldsymbol{\theta})$ in terms of the contributions made by each branch of $\mathcal{T}$. We thus define the *branch genetic prior* $\mathcal{G}_i(\mathcal{T}|\boldsymbol{\theta})$ for branches $i$ other than the root branch as follows:

$$\mathcal{G}_i(\mathcal{T}|\boldsymbol{\theta}) := \exp\left[ -\int_{x\in B_i} \lambda(x)\, dx \right] \cdot \prod_{(a,\ell,b)\in\mathcal{M}_i} Q^{(\ell)}_{ab}, \qquad (i \neq r). \tag{5}$$

An important simplification follows if we define the branch genetic prior of the root branch, $\mathcal{G}_r(\mathcal{T}|\boldsymbol{\theta})$, as follows:

$$\mathcal{G}_r(\mathcal{T}|\boldsymbol{\theta}) = \prod_{\ell\in\xi(r)} \pi^{(\ell)}_{s_\perp^{(\ell)}} \cdot \prod_{(a,\ell,b)\in\mathcal{M}_r} (\pi^{(\ell)}_b / \pi^{(\ell)}_a).$$

With this definition, most subsequent manipulations need not distinguish between the root and nonroot branches. For example, the complete genetic prior is given simply by:

$$\mathcal{G}(\mathcal{T}|\boldsymbol{\theta}) = \prod_i \mathcal{G}_i(\mathcal{T}|\boldsymbol{\theta}).$$

In the definition of $\mathcal{G}_r(\mathcal{T}|\boldsymbol{\theta})$, we have also factored out the contribution of the reference sequence for the sites $\xi(r)$ that are missing in every input sequence; the leftover factors then arise from mutations above the root. By arranging for most MCMC moves to leave $s_\perp^{(\ell)}$ unchanged ($\xi(r)$ is fixed throughout the run), only the factors arising from mutations above the root survive when taking ratios of branch genetic priors for the root branch.

We define path and subtree genetic priors in the obvious way as products of the relevant branch genetic priors.

Just as it is sometimes convenient to split the calculation of the genetic prior according to the contributions made by each branch, it is also sometimes convenient to split it according to the contributions made by each site. Thus, we define the *site genetic prior* $\mathcal{G}^{(\ell)}(\mathcal{T}|\boldsymbol{\theta})$ as the factors of $\mathcal{G}(\mathcal{T}|\boldsymbol{\theta})$ that relate to site $\ell$. Explicitly,

$$p\mathcal{G}^{(\ell)}(\mathcal{T}|\boldsymbol{\theta}) = \begin{cases} \pi^{(\ell)}_{s_r^{(\ell)}} \cdot \exp\left[-\int_{x\in\mathcal{T}} \lambda^{(\ell)}(x)\,dx\right] \cdot \prod_{(a,b)\in\mathcal{M}^{(\ell)}} Q^{(\ell)}_{ab}, & \ell \in \xi(r); \\ 1, & \text{otherwise.} \end{cases} \tag{6}$$

Here, $\lambda^{(\ell)}(x)$ is either the single term $Q^{(\ell)}_{s_x^{(\ell)}}$ if $\ell \in \xi(x)$, or 0 otherwise; and $\mathcal{M}^{(\ell)}$ is the subset of mutations in $\mathcal{M}$ occurring at site $\ell$. By construction,

$$\mathcal{G}(\mathcal{T}|\boldsymbol{\theta}) = \prod_\ell \mathcal{G}^{(\ell)}(\mathcal{T}|\boldsymbol{\theta}).$$

Clearly, the site genetic prior is closely related to the traditional site likelihood. This relation is detailed in Section 4.

Finally, we can decompose $\mathcal{G}(\mathcal{T}|\boldsymbol{\theta})$ yet more finely into a product of *site-branch genetic priors* $G_i^{(\ell)}(\mathcal{T}|\boldsymbol{\theta})$ as follows:

$$\mathcal{G}_i^{(\ell)}(\mathcal{T}|\boldsymbol{\theta}) = \begin{cases} \pi^{(\ell)}_{s_r^{(\ell)}}, & i = r \text{ and } \ell \in \xi(r); \\ \exp\left[-\int_{x\in\mathcal{B}_i} \lambda^{(\ell)}(x)\,dx\right] \cdot \prod_{(a,b)\in\mathcal{M}_i^{(\ell)}} Q^{(\ell)}_{ab}, & i \neq r \text{ and } \ell \in \xi(r); \\ 1, & \text{otherwise.} \end{cases} \tag{7}$$

As above, $\mathcal{M}_i^{(\ell)}$ is the subset of mutations in $\mathcal{M}$ occurring at site $\ell$ on branch $i$.

Finally, we remark that calculating the integral $\int_{x\in B_i} \lambda(x)\,dx$, and thus $\mathcal{G}(\mathcal{T}|\boldsymbol{\theta})$, is considerably simplified if we keep track of the value of $\lambda(x)$ at every node $i$, which we denote by $\lambda_i$. First, we note that we always store the mutations in $\mathcal{M}_i$ sorted by ascending time. Label their identities by the tuples $(a_j, \ell_j, b_j, t_{i,j})$ for $1 \le j \le |\mathcal{M}_i|$ in the obvious way, and set $t_{i,0} = t_{\varphi(i)}$ and $t_{i,|\mathcal{M}_i|+1} = t_i$. Then compute the quantities $\lambda_{i,j}$, which are the values of $\lambda(x)$ between mutations $j$ and $j+1$, with $0 \le j \le |\mathcal{M}_i|$. These obey the following recursion relations:

$$\lambda_{i,|\mathcal{M}_i|} = \lambda_i, \tag{8}$$

$$\lambda_{i,j-1} = \lambda_{i,j} - Q^{(\ell_j)}_{b_j} + Q^{(\ell_j)}_{a_j}, \qquad (1 \le j \le |\mathcal{M}_i|). \tag{9}$$

With these quantities, we obtain:

$$\int_{x\in B_i} \lambda(x)\,dx = \sum_{j=0}^{|\mathcal{M}_i|} \lambda_{i,j}(t_{i,j+1} - t_{i,j}).$$

To calculate $\lambda_i$ for all nodes, note that there is a simple recurrence relation between the total mutation rate at a node $i$ and that at its ancestor:

$$\lambda_i = \lambda_{\varphi(i)} + \sum_{(a,\ell,b)\in\mathcal{M}_i} (Q^{(\ell)}_b - Q^{(\ell)}_a) - \sum_{(a,\ell)\in\Phi_i} Q^{(\ell)}_a. \tag{10}$$

Since $\lambda_\perp = \sum_\ell Q^{(\ell)}_{s_\perp^{(\ell)}}$, it follows that having calculated $\lambda_\perp$, the subsequent calculation of $\lambda_i$ for every node $i$ can be done in time linear with the size of the tree and the number of mutations and missations on its branches. Hence, $\mathcal{G}(\mathcal{T}|\boldsymbol{\theta})$ can also be calculated in linear time, and changes to it arising from changes to a subtree can be calculated in time linear with the number of mutations and missations in that subtree.

##### 3.3 Ancestry prior

Any ancestry prior suitable for Bayesian phylogenetics can be used in our framework. For concreteness, we have implemented the traditional Kingman coalescent prior[7, 8]. A significant complication in parallelizing the coalescent prior is that it directly couples all the nodes at the end of branches that cross a given time $t$. We discuss the details of the ancestry prior and an equivalent formulation that is parallelizable in Section 9.

##### 3.4 Parameter priors

The genetic prior is defined in terms of the site transition rate matrices $Q^{(\ell)}_{ab}$. Our framework accommodates all common choices for these, e.g., multiple site partitions with linked or unlinked mutation rates, with or without site rate heterogeneity, with more (GTR) or fewer (HKY) free exchangeabilites, with some proportion of invariant sites, etc. With minor modifications, more elaborate schemes can also be accommodated, e.g., uncorrelated relaxed molecular clocks. Similarly, the ancestral prior is defined in terms of the population curve $N(t)$, and all models in common use can be applied, from a fixed flat population, to parametrized exponential growth, to more flexible schemes such as a Skygrid prior [17].

For concreteness, however, we have thus far only implemented an evolution model consisting of a single-partition HKY model with optional site-rate heterogeneity and no invariant sites, and a population model representing exponential growth, all with hard-coded parameters to mimic the model used in [11][1]. We leave it to future work to implement more flexible specifications of evolution and population models, along with their respective MCMC moves.

In detail, the concrete evolution model we have implemented is:

$$Q^{(\ell)}_{ab} = \mu\nu^{(\ell)} q_{ab},$$

where $\mu$ is the overall mutation rate, $\nu^{(\ell)}$ is a site-relative rate, and $q_{ab}$ is the usual reduced transition rate matrix for the HKY model with stationary base frequencies $\pi_a$ and transition-transversion ratio $\kappa$, given by:

$$q_{ab} = r_{ab}\pi_b/R, \quad (a \neq b) \qquad \text{with } r_{ab} := \begin{pmatrix} 0 & 1 & \kappa & 1 \\ 1 & 0 & 1 & \kappa \\ \kappa & 1 & 0 & 1 \\ 1 & \kappa & 1 & 0 \end{pmatrix} \text{ and } R := \sum_{ab} \pi_a r_{ab} \pi_b.$$

The hyperparameters $\mu$, $\nu^{(\ell)}$, $\pi_a$ and $\kappa$ have the same priors as those produced by default when configuring a BEAST 2 run with BEAUti 2 for an HKY model and a strict molecular clock, which are those used in [11]:

- Mutation rate $\mu$: a uniform, improper prior,

$$\pi_\mu(\mu) = 1.$$

- Site relative mutation rates: We either set $\nu^{(\ell)} = 1$ for all $\ell$ (no site rate heterogeneity), or we choose $\nu^{(\ell)}$ to be Gamma distributed with mean 1 and shape parameter $\alpha$:

$$\pi_\nu(\{\nu^{(\ell)}\}|\alpha) = \prod_\ell \frac{\alpha^\alpha}{\Gamma(\alpha)} \left[\nu^{(\ell)}\right]^{\alpha-1} e^{-\alpha\nu^{(\ell)}}.$$

  We emphasize that in our framework, all the $\nu^{(\ell)}$ parameters are explicit, independent, and continuous, not restricted in the usual way to discrete categories [26]. The site rate heterogeneity parameter $\alpha$ is an additional latent parameter, itself associated with a prior distribution $\pi_\alpha(\alpha)$, which we take to be exponential with mean $\mu_\alpha = 1$:

$$\pi_\alpha(\alpha) = \frac{1}{\mu_\alpha} e^{-\alpha/\mu_\alpha}.$$

[1]We have also successfully run a two-partition model in the context of analyzing mpox sequences, where some sites are susceptible to fast APOBEC-mediated mutations[20] while other sites exhibit only mutations from polymerase errors (unpublished).

- Evolution model parameters: the stationary base frequencies have a $\mathrm{Dir}(1,1,1,1)$ prior, while $\kappa$ has a log-normal prior with mean $\mu_{\ln\kappa}$ and variance $\sigma^2_{\ln\kappa}$:

$$\pi_Q(\kappa,\{\pi_a\}) = 1 \cdot \frac{1}{\ln\kappa \cdot \sqrt{2\pi\sigma^2_{\ln\kappa}}} e^{-(\ln\kappa - \mu_{\ln\kappa})^2/2\sigma^2_{\ln\kappa}}.$$

Similarly, the concrete population model we have implemented is that of exponential growth,

$$N(t) := n_0 \cdot e^{g(t-t_0)},$$

where $t_0$ is the time of the latest tip. As with the evolution model, we use the same priors for $n_0$ and $g$ that are output by default by BEAUti 2 when preparing a BEAST 2 run with a "Coalescent Exponential Population", namely:

- An improper $1/x$-prior on the effective population size $n_0$ at time $t_0$,

$$\pi_{n_0}(n_0) = 1/n_0.$$

- A Laplace prior on the growth rate $g$, with mean $\mu_g = 0.001\,\mathrm{yr}^{-1}$ and scale $s_g = 30.701135\,\mathrm{yr}^{-1}$.

$$\pi_g(g) = \frac{1}{2s_g} \exp\left[-\frac{|g-\mu_g|}{s_g}\right].$$

With the concrete choices above, the overall prior on latent parameters is simply the product of the above priors:

$$\pi_{\boldsymbol{\theta}}(\boldsymbol{\theta}) = \pi_\mu(\mu)\cdot\pi_\nu(\{\nu^{(\ell)}\}|\alpha)\cdot\pi_\alpha(\alpha)\cdot\pi_Q(\kappa,\{\pi_a\})\cdot\pi_{n_0}(n_0)\cdot\pi_g(g).$$

### 4 Genetic prior vs. tree likelihood

In this section, we explicitly establish that integrating the genetic prior (Equation (3)) over all possible EMATs with fixed topology yields the traditional tree likelihood, as computed via Felsenstein pruning [4]. Hence, what distinguishes Delphy from the traditional Bayesian methods based on Felsenstein pruning is that it samples over explicit topologies and genetic histories using a simple posterior, instead of integrating out the genetic histories analytically and then sampling only over topologies using the resulting more computationally demanding posterior. Otherwise, the methods are statistically equivalent. A reader comfortable with this standard result can safely skip this section.

Throughout this section, we assume that no data is missing. Section 5 details how implicit tracking of some ambiguous sites minimally alters the genetic prior.

First, we make precise the notion of "integrating over all genetic histories". Consider the evolution of single site over a single lineage and a fixed period of time. Define a site genetic history $\mathcal{M}$ as a set of $K \geq 0$ mutations, specified as tuples $(t_k, u_k, v_k)$ where the state at the site changes from $u_k$ to $v_k$ at time $t_k$. Such a history satisfies three coherency conditions: (a) $u_k \neq v_k$ for all $k$; (b) $v_k = u_{k+1}$ for $0 \leq k < K$; and (c) $t_1 < \cdots < t_K$. Let $\mathcal{H}_{ab}(p,q)$ be the set of site genetic histories that further have starting state $a$ at time $p$ and an ending state $b$ at time $q$. Formally, this means that each such site genetic history satisfies the following additional conditions: (a) if $K > 0$, then $u_1 = a$ and $v_K = b$; otherwise, if $K = 0$, then $\mathcal{H}_{ab}(p,q)$ is nonempty only when $a = b$, in which case, the only genetic history it contains is the empty set; (b) $p < t_1$ and $t_K < q$. We define an integral over all such site genetic histories as follows:

$$\int_{\mathcal{H}_{ab}(p,q)} d\mathcal{M}\, f(\mathcal{M}) := \delta_{ab} f(\emptyset) + \sum_{K=1}^{\infty} \sum_{\{u_k,v_k\}}{}' \int_{p<t_1<\cdots<t_K<q} dt_1 \cdots dt_K\, f(\mathcal{M}).$$

The primed sum is over all possible assignments of states to $u_k$ and $v_k$ that produce a valid site genetic history in $\mathcal{H}_{ab}(p,q)$.

We define $R^{(\ell)}_{ab}(p,q)$ as the integral over all site genetic histories in $\mathcal{H}_{ab}(p,q)$ of the kinds of factors that appear in the site genetic prior $\mathcal{G}^{(\ell)}(\mathcal{T}|\boldsymbol{\theta})$:

$$R^{(\ell)}_{ab}(p,q) := \int_{\mathcal{H}_{ab}(p,q)} d\mathcal{M}\, \exp\left[-\int_p^q dt\, Q^{(\ell)}_{s^{(\ell)}(t)}\right] \cdot \prod_{(a,b)\in\mathcal{M}} Q^{(\ell)}_{ab} \tag{11}$$

Here, $s^{(\ell)}(t)$ is the state of site $\ell$ at time $t$ according to the genetic history $\mathcal{M}$. That is, first $s^{(\ell)}(p) = a$, then $s^{(\ell)}(t)$ assumes the various values of $v_k$ as $t$ crosses $t_k$, until finally, $s^{(\ell)}(q) = b$.

We show below that the above integral over all genetic histories is closely related to the traditional evolution matrix $\boldsymbol{P}^{(\ell)}(t)$. The element $P^{(\ell)}_{ab}(t)$ of this matrix is the probability that a state $a$ at site $\ell$ evolves to a state $b$ over a time $t$ along a single lineage. This matrix is defined by the forward Kolgomorov equation of the Markov process and a suitable initial condition:

$$\frac{dP^{(\ell)}_{ab}(t)}{dt} = \sum_c P^{(\ell)}_{ac}(t) Q^{(\ell)}_{cb};$$
$$P^{(\ell)}_{ab}(0) = \delta_{ab}.$$

For time-independent $Q^{(\ell)}_{ab}$, one obtains the usual result, $\boldsymbol{P}^{(\ell)}(t) = \exp(\boldsymbol{Q}^{(\ell)}t)$. We note in passing that our treatment below is almost unchanged for time-dependent $Q^{(\ell)}_{ab}$, and may thus provide a tractable way of modeling time-inhomogeneous evolution in a Bayesian phylogenetics context.

We can now state the key result of this section, namely, that the above integral over all site genetic histories evaluates to an evolution matrix element:

$$R^{(\ell)}_{ab}(p,q) = P^{(\ell)}_{ab}(q-p). \tag{12}$$

To show this, we rewrite the definition of $R^{(\ell)}_{ab}(p,q)$ in Equation (11) by first separating the term where there are no mutations from $p$ to $q$, then writing the remainder in terms of the time $t'$ of the last mutation and the site genetic history preceding that mutation. For later convenience, we also replace $Q^{(\ell)}_b$ by $-Q^{(\ell)}_{bb}$:

$$R^{(\ell)}_{ab}(p,q) := \delta_{ab} e^{Q^{(\ell)}_{bb}(q-p)} + \int_p^q dt' \sum_{c\neq b} R^{(\ell)}_{ac}(p,t') Q^{(\ell)}_{cb} e^{Q^{(\ell)}_{bb}(q-t')}.$$

We then calculate the derivative of $R^{(\ell)}_{ab}(p,q)$ with respect to $q$:

$$\begin{aligned}\frac{dR^{(\ell)}_{ab}(p,q)}{dq} &= \delta_{ab} e^{Q^{(\ell)}_{bb}(q-p)} Q^{(\ell)}_{bb} \\ &\quad + \int_p^q dt' \sum_{c\neq b} R^{(\ell)}_{ac}(p,t') Q^{(\ell)}_{cb} e^{Q^{(\ell)}_{bb}(q-t')} Q^{(\ell)}_{bb} \\ &\quad + \sum_{c\neq b} R^{(\ell)}_{ac}(p,q) Q^{(\ell)}_{cb} \\ &= R^{(\ell)}_{ab}(p,q) Q^{(\ell)}_{bb} + \sum_{c\neq b} R^{(\ell)}_{ac}(p,q) Q^{(\ell)}_{cb} \\ &= \sum_c R^{(\ell)}_{ac}(p,q) Q^{(\ell)}_{cb}.\end{aligned}$$

We also observe that when $p = q$, we have:

$$R^{(\ell)}_{ab}(p,p) = \delta_{ab}.$$

Since $R_{ab}^{(\ell)}(p,q)$ satisfies the same differential equation and initial conditions as $P_{ab}^{(\ell)}(t)$, we conclude they are equivalent (Equation (12)).

Next, we generalize the concept of site genetic histories to encompass mutations at any site along a single branch $j$. For convenience, let $i = \varphi(j)$. Thus, branch $j$ starts at node $i$, with time $t_i$, and ends at node $j$, with time $t_j$. Define a branch genetic history $\mathcal{M}_j$ on branch $j$ as a set of $K \geq 0$ mutations, specified as tuples $(j, t', a', \ell, b')$, where $t_i < t' < t_j$ and $1 \leq \ell \leq L$. Each tuple records that at time $t'$, the state of site $\ell$ on branch $j$ changes from $a'$ to $b'$. A branch genetic history can be partitioned into individual site genetic histories, each of which must satisfy the coherency conditions (a)–(c) given at the beginning of this section. Denote by $\mathcal{H}_j$ the set of branch histories over branch $j$, compatible with fixed sequences $\{s_i^{(\ell)}\}$ and $\{s_j^{(\ell)}\}$ at its endpoints. Thus, an arbitrary member of $\mathcal{H}_j$ consists of the union of $L$ sets, one for each site $\ell$, derived each from $\mathcal{H}_{s_i^{(\ell)}\, s_j^{(\ell)}}(t_i, t_j)$. We define an integral over all branch genetic histories in $\mathcal{H}_j$ as follows:

$$\int_{\mathcal{H}_j} d\mathcal{M}_j\, f(\mathcal{M}_j) := \left[\prod_\ell \int_{\mathcal{H}_{s_i^{(\ell)}\, s_j^{(\ell)}}(t_i,t_j)} d\mathcal{M}^{(\ell)}\right] f\Big(\bigcup_\ell g_j^{(\ell)}[\mathcal{M}^{(\ell)}]\Big). \tag{13}$$

The function $g_j^{(\ell)}[\mathcal{M}^{(\ell)}]$ maps each tuple $(t_k, u_k, v_k)$ in $\mathcal{M}^{(\ell)}$ to $(j, t_k, u_k, \ell, v_k)$.

With the above definitions in place, we note that the product of all the $R_{ab}^{(\ell)}(p,q)$ factors associated with a given branch $j$ can be simply expressed as an integral over branch genetic histories in $\mathcal{H}_j$, where mutations can occur over any site. Explicitly, it follows from Equations (11) and (13) that

$$\prod_\ell R^{(\ell)}_{s_i^{(\ell)}\, s_j^{(\ell)}}(t_i, t_j) = \int_{\mathcal{H}_j} d\mathcal{M}_j\, \exp\left[-\int_{t_i}^{t_j} dt \sum_\ell Q^{(\ell)}_{s^{(\ell)}(t)}\right] \cdot \prod_{(a,\ell,b)\in\mathcal{M}_j} Q^{(\ell)}_{ab}.$$

Using Equation (12) to express the left-hand side in terms of evolution matrix elements, and then using Equation (2) for the total mutation rate $\lambda(x)$ at a point $x$ on the tree (when all sites are explicitly tracked everywhere), we obtain the equivalent relation

$$\prod_\ell P^{(\ell)}_{s_i^{(\ell)}\, s_j^{(\ell)}}(t_j - t_i) = \int_{\mathcal{H}_j} d\mathcal{M}_j\, \exp\left[-\int_{x\in\mathcal{B}_j} \lambda(x)\, dx\right] \cdot \prod_{(a,\ell,b)\in\mathcal{M}_j} Q^{(\ell)}_{ab} = \int_{\mathcal{H}_j} d\mathcal{M}_j\, \mathcal{G}_j(\mathcal{T}|\boldsymbol{\theta}).$$

At the end, we've noted that the integrand above is just the branch genetic prior $\mathcal{G}_j(\mathcal{T}|\boldsymbol{\theta})$ of Equation (5).

We now further generalize genetic histories to cover the entire tree. Define a tree genetic history $\mathcal{M}_\mathcal{T}$ as a union of branch genetic histories, one for each branch except the root branch. A tree genetic history must satisfy the obvious coherency condition that the ending state of the branch genetic history of branch $\varphi(i)$ is the starting state of branch $i$. Denote by $\mathcal{H}_\mathcal{T}$ the set of tree genetic histories compatible with fixed sequences $\{s_i^{(\ell)}\}$ at every node $i$, including the root. We define an integral over all tree genetic histories in $\mathcal{H}_\mathcal{T}$ as follows:

$$\int_{\mathcal{H}_\mathcal{T}} d\mathcal{M}_\mathcal{T}\, f(\mathcal{M}_\mathcal{T}) := \left[\prod_{j\neq r} \int_{\mathcal{H}_j} d\mathcal{M}_j\right] f\Big(\bigcup_j \mathcal{M}_j\Big). \tag{14}$$

With the above definition, we can finally relate the full genetic prior to a product of evolution matrix elements and base frequencies $\pi_a$:

$$\int_{\mathcal{H}_\mathcal{T}} d\mathcal{M}_\mathcal{T}\, \mathcal{G}(\mathcal{T}|\boldsymbol{\theta}) = \prod_\ell \left[\pi^{(\ell)}_{s_r^{(\ell)}} \cdot \prod_{j\neq r} P^{(\ell)}_{s_i^{(\ell)}\, s_j^{(\ell)}}(t_j - t_i)\right]. \tag{15}$$

To complete the link between this work and Bayesian methods based on Felsenstein pruning, we first recast in our notation the traditional tree likelihood as computed via Felsenstein pruning [4]:

$$L_{\text{tree}}(\mathcal{T}_{\text{FT}}|\boldsymbol{\theta}) = \prod_{\ell} \left\{ \sum_{\substack{\{s_i^{(\ell)}\} \\ \text{fixed } \ell \\ \text{inner } i}} \left[ \pi_{s_r^{(\ell)}} \cdot \prod_{j \neq r} P^{(\ell)}_{s^{(\ell)}_{\varphi(j)}\, s^{(\ell)}_j}(t_j - t_i) \right] \right\}. \tag{16}$$

The tree $\mathcal{T}_{\text{FT}}$ appearing on the left-hand side is the tree $\mathcal{T}$ stripped of its mutational history, i.e., the object that is usually tracked when using Felsenstein pruning. The outer product on the right-hand side is over all sites $\ell$, so the factor in braces is the site likelihood of site $\ell$. The inner sum expands into $N-1$ nested sums over the possible values of all states $s_i^{(\ell)}$ of all inner nodes $i$, for a fixed $\ell$ (the values of $s_i^{(\ell)}$ for tips $i$ are simply the input sequences). The factor $\pi_{s_r^{(\ell)}}$ is the stationary base frequency of the state at site $\ell$ of the root node. The first innermost product is over all nonroot branches $j$ and its factors are the matrix elements of the evolution matrix $P_{ab}^{(\ell)}(t)$.

By distributing the first product over the sum in Eq. (16), we obtain

$$L_{\text{tree}}(\mathcal{T}_{\text{FT}}|\boldsymbol{\theta}) = \sum_{\substack{\{s_i^{(\ell)}\} \\ \text{all } \ell \\ \text{inner } i}} \left\{ \prod_{\ell} \left[ \pi_{s_r^{(\ell)}} \cdot \prod_{j \neq r} P^{(\ell)}_{s^{(\ell)}_{A_j}\, s^{(\ell)}_j}(t_j - t_i) \right] \right\}. \tag{17}$$

Note that the sum in Eq. (16) expands into $N-1$ simpler nested sums, whereas that in Eq. (17) above expands into $(N-1) \times L$ simpler nested sums.

Substituting Equation (15) into the above, we immediately obtain the desired equivalence between the traditional tree likelihood on the one hand, and integrals over genetic histories of the product of the genetic prior and the likelihood on the other hand:

$$L_{\text{tree}}(\mathcal{T}_{\text{FT}}|\boldsymbol{\theta}) = \sum_{\substack{\{s_i^{(\ell)}\} \\ \text{all } \ell \\ \text{inner } i}} \int_{\mathcal{H}_{\mathcal{T}}} d\mathcal{M}_{\mathcal{T}}\, \mathcal{G}(\mathcal{T}|\boldsymbol{\theta}) \tag{18}$$

Similar considerations lead to the following link between the traditional site likelihood and integrals over genetic histories of the site genetic prior and the site likelihood:

$$L^{(\ell)}_{\text{tree}}(\mathcal{T}_{\text{FT}}|\boldsymbol{\theta}) = \sum_{\substack{\{s_i^{(\ell)}\} \\ \text{fixed } \ell \\ \text{inner } i}} \int_{\mathcal{H}^{(\ell)}_{\mathcal{T}}} d\mathcal{M}^{(\ell)}_{\mathcal{T}}\, \mathcal{G}^{(\ell)}(\mathcal{T}|\boldsymbol{\theta}).$$

The above derivations can be generalized straightforwardly in various ways, e.g., to account for site weight vectors at the tips, non-strict clocks, etc.

### 5 Dealing with missing data: `N`-pruning

In Section 4, we established that if we track all sites explicitly everywhere, then integrating the product of the genetic prior and the likelihood over all genetic histories yields the traditional tree likelihood. Throughout this paper, we argue that it's enormously beneficial to perform this integration statistically instead of analytically. However, in the presence of ambiguous tip sequences, a mixed strategy, which we call "`N`-pruning", retains almost all the benefits of the explicit approach while treating sequence ambiguity efficiently.

To illustrate the basic idea, consider the portion of a 1-site tree depicted in Figure 12(a). It includes 2 tips, $A$ and $B$, their common ancestor, $C$, and another ancestor further removed, $D$. For clarity, we omit site superscripts in this 1-site example. Tip $A$ is an N while tip $B$ is a T. The traditional partial site likelihood at $D$ is:

$$L_d = \sum_{a,c} P_{dc}(\tau_C) P_{ca}(\tau_A) P_{c\mathtt{T}}(\tau_B).$$

Note that we sum over all possible states $a$ of tip $A$, but the state of tip $B$ is fixed to T. The summation over $a$ is simple to perform explicitly:

$$L_d = \sum_{a,c} P_{dc}(\tau_C) P_{ca}(\tau_A) P_{c\mathtt{T}}(\tau_B),$$
$$= \sum_{c} P_{dc}(\tau_C) P_{c\mathtt{T}}(\tau_B).$$

This follows from $\sum_a P_{ca}(\tau_A) = 1$, i.e., the state $c$ definitely evolves to *something* after a time $\tau_A$. We now apply the Markov property of the evolution process, expressed as $\sum_c P_{dc}(\tau_C) P_{c\mathtt{T}}(\tau_B) = P_{d\mathtt{T}}(\tau_C + \tau_B)$, to evaluate the sum over $c$:

$$L_d = P_{d\mathtt{T}}(\tau_C + \tau_B).$$

Finally, observe that this is just the traditional partial site likelihood at $D$ of the tree depicted in Figure 12(b), which is the result of "pruning" the branch that ended in an N from the original tree.

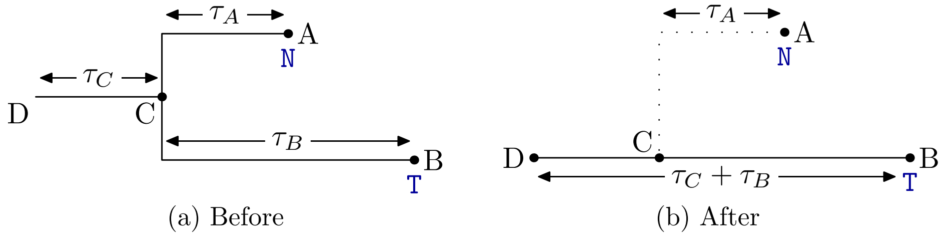


**Supplementary Figure 12:** N-pruning: Integrating over all possible histories of branch $C$–$A$ is equivalent to pruning it off.

Clearly, the above pruning process can be repeated recursively to show that the site likelihood of a tree with ambiguous tips is equivalent to the site likelihood of the same tree with all subtrees ending in N tips pruned off. By the arguments of Section 4, this is equivalent to restricting the products, sums and integrals in Equation (3) according to $\xi(x)$. Thus, missations are revealed to be the pruning points at particular sites for each of the site trees that make up $\mathcal{T}$. The equivalence is sketched out in Figure 13.

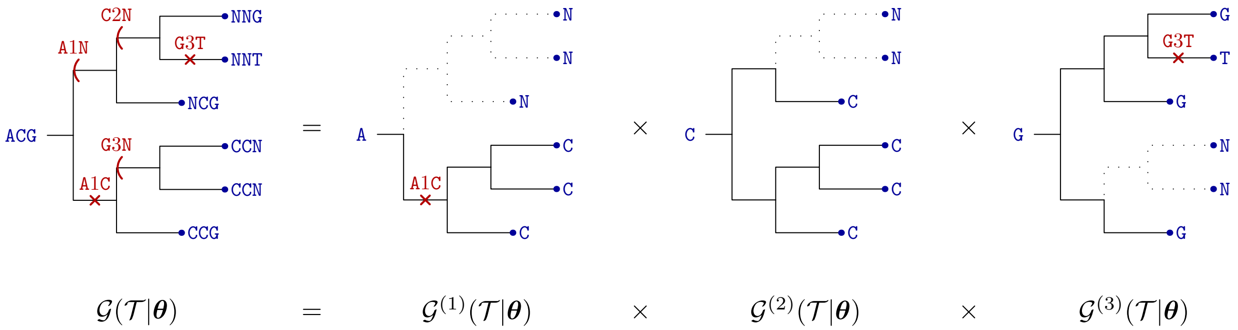


**Supplementary Figure 13:** N-pruning: A tree with "missations" decomposes into site trees, each of which has had its subtrees ending in N-tips pruned off. The genetic prior for the whole tree equals the product of site genetic priors for the pruned site trees.

In a very real sense, an EMAT with missations stands for all trees resulting from realizing all possible genetic histories below each missation. In particular, the space of all possible EMATs without missations is thus partitioned by EMATs with missations into sets, according first to topology, then to genetic histories above the missations. The posterior probability of an EMAT with missations is then sum of the posterior probabilities of the contained EMATs without missations. In order for this scheme to work, it is essential that any one EMAT without missations be "contained" in exactly one EMAT with missations, as indeed they are.

Empirically, we have found that very local MCMC moves are effective for EMATs with explicitly tracked sites everywhere when the tips are densely sampled, as is the case for SARS-CoV-2 and will likely be the case when understanding outbreaks and/or future pandemics. This is compatible with the observation that most trees linking such densely sampled datasets are nearly parsimonious [6, 9]. Applying `N`-pruning to all tips with missing data (converting partially ambiguous sites into totally ambiguous ones where necessary) effectively reduces the problem of sampling trees with missing data to that of sampling smaller trees without missing data, so very local MCMC moves are also effective there.

The alternative to implicit tracking of sites with data is to explicitly realize and modify genetic histories that lead up to tips with ambiguous state. Implicit tracking allows us to encode large-scale rearrangements with small structural changes. For example, in Figure 14, by far the most likely state of site 1 in all tips below $X$ is `A`. An SPR move (Section 6.1) that detaches the subtree rooted at $X$ from $P$ and reattaches it at $P'$ changes the most likely state at site 1 everywhere downstream of $X$ to `C`, while only requiring very local rearrangements of the EMAT. In explicit tracking, local moves that do not perform this large-scale rearrangement are unlikely to be accepted because they introduce spurious additional mutations. We empirically verified that implementing only such local moves, e.g., an SPR move on tips that allows for the tip sequence to change where it is ambiguous, did not result in efficient sampling. While it is possible to improve the local moves we present in the following section to rearrange mutations in large clusters of the tree (perhaps inspired by the set of branches that an equivalent move in a parsimony tree would affect[27]), we have not yet explored this possibility fully. Certainly, `N`-pruning allows us to sidestep this issue for models where sites evolve independently and for datasets where sites are either fully determined by experiment or are completely ambiguous. This seems to cover the most common use cases we have seen for Bayesian phylogenetics. Nevertheless, a different approach would need to be developed to handle tip sites that are only partially ambiguous (i.e., for arbitrary weight vectors that model sequencer error), or to handle ambiguity when sites do not evolve independently (e.g., to correctly account for context-dependent APOBEC- or ADAR-induced mutations).

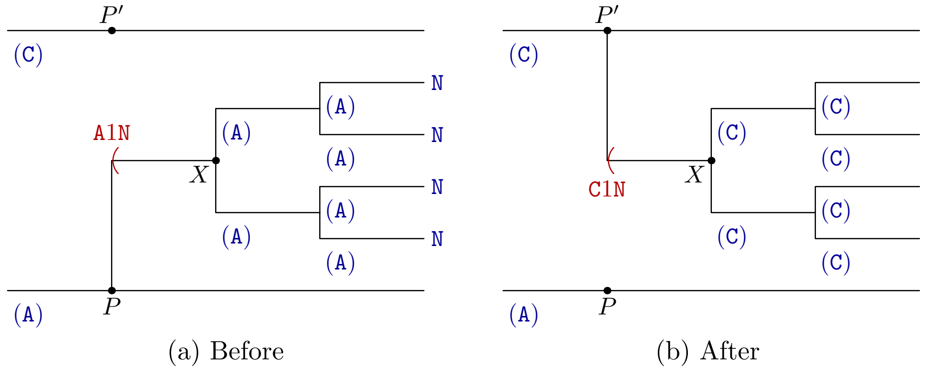


**Supplementary Figure 14:** `N`-pruning permits large-scale rearrangements with only local changes: By changing the single missation at the beginning of branch $X$ from `A2N` to `C2N`, the implied sequences downstream of $X$ all change from `A` to `C`.

One final complication with missing data is that often a sequence will have large numbers of consecutive sites that are missing, forming "gaps". To take one concrete if arbitrary example, GenBank sequence

OP612676, a nearly 200kb mpox sequence, has more than a dozen gaps each over 100 bases long, including one that is 5653 bases long. It's therefore essential to have an efficient representation of consecutive missations, different from a naive list with one entry per site. Further, efficient MCMC moves require knowledge of the state of a site preceding a missation (to implement the last term of the recurrence relation for $\lambda_i$ in Equation (10)). Fortunately, for outbreak-like datasets, the probability that a random site differs in state at a point $x$ on the tree from its state at the root is relatively small (for example, after 4 years of SARS-CoV-2 evolution, a typical sequence differs from the Wuhan reference sequence NC_045512.2 at only 150 or so of its 30 000 sites). Since high-quality sequences rarely have more than a few percent of sites with missing data, the probability that a random *missing* site has a state different from the reference state is even smaller. Hence, inspired by MAPLE's representation for missing data[12], we represent $\Phi_i$, the missations at the start of branch $i$, as a set of disjoint sorted intervals of sites that have no informative tips downstream of $\varphi(i)$, coupled with a sparse map from site $\ell$ to state $s_i^{(\ell)}$ for sites where this state differs from the reference state $s_\perp^{(\ell)}$. This representation admits efficient union and intersection operations, which we need to implement MCMC moves below, and the sparse map rarely contains more than a handful of entries. We further keep track of the cumulative sum of the terms $Q^{(\ell)}_{s_\perp^{(\ell)}}$, which allows us to implement the recurrence relation of Equation (10) even in the presence of multi-thousand base gaps.

We note briefly that the N-pruning scheme is not restricted to Bayesian phylogenetics. It may thus be possible to adapt parsimony and maximum likelihood approaches to use N-pruning with resulting efficiency improvements. Such an exploration is outside the scope of this paper.

Undoubtedly, N-pruning makes implementing MCMC moves on EMATs more complicated. We spell out all the details below.

### 6 Local MCMC moves

Here, we consider *local MCMC moves*, that is, moves that rearrange the tree $\mathcal{T}$ while keeping all the associated inference parameters $\boldsymbol{\theta}$ fixed. Hence, for clarity, we suppress the dependence on $\boldsymbol{\theta}$ on all quantities throughout this section.

A local move starts with an old configuration $\mathsf{o}$ of the tree and proposes a new configuration $\mathsf{n}$ with probability $\alpha(\mathsf{o} \to \mathsf{n})$. As usual, the move is accepted with a probability $P_{\text{acc}}(\mathsf{o} \to \mathsf{n})$ according to the Metropolis-Hastings criterion[5, 16],

$$P_{\text{acc}}(\mathsf{o} \to \mathsf{n}) = \min\left[1, \frac{P(\mathsf{n})}{P(\mathsf{o})}\frac{\alpha(\mathsf{n} \to \mathsf{o})}{\alpha(\mathsf{o} \to \mathsf{n})}\right]. \tag{19}$$

Here, $P(\mathcal{T})$ is the posterior of configuration $\mathcal{T}$, as given in Equation (1). Since the values of the parameters $\boldsymbol{\theta}$ are unchanged in local moves, the ratio of the posterior involves only ratios of the genetic and ancestral priors:

$$\frac{P(\mathsf{n})}{P(\mathsf{o})} = \frac{\mathcal{G}(\mathsf{n})}{\mathcal{G}(\mathsf{o})}\frac{\pi_{\text{anc}}(\mathsf{n})}{\pi_{\text{anc}}(\mathsf{o})}.$$

The ratio of ancestral priors is easily calculated for all local moves, whereas calculating the ratios of proposal probabilities and genetic priors involves many subtleties. We describe the latter two calculations in detail below for all the local MCMC moves that we have implemented.

#### 6.1 Subtree Pruning and Regrafting (SPR) moves

Delphy implements a subtree pruning and regrafting move that is conceptually related to Nielsen's original SPR move[19]. The main differences are: (a) the choice of grafting point is made general, with some specific choices detailed below; (b) the mutation proposal is adapted for efficiency for large genomes with few mutations per branch; (c) the move is generalized to arbitrary evolution models (not necessarily time-reversible); and (d) the move efficiently handles tip sequences with missing data.

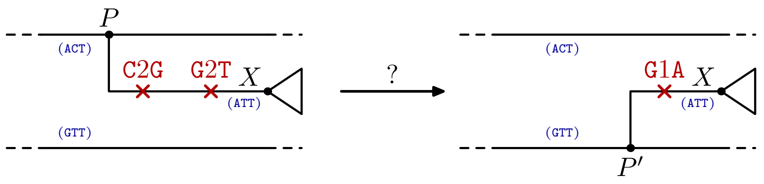


**Supplementary Figure 15:** A generic SPR move: a subtree rooted at $X$ is attached to the rest of the tree at point $P$. An SPR move proposes detaching this subtree and reattaching it at a different point $P'$. The move includes a proposal for a new mutational history, at the very least covering the $P'$–$X$ branch.

We first describe the move when all data is present and the root is not changing, then show how these restrictions are relaxed in the subsequent sections.

A general SPR move is depicted in Figure 15. A subtree rooted at $X$, grafted at $P$ in the old configuration $\mathsf{o}$, is to be regrafted at another point $P'$ in the new configuration $\mathsf{n}$. The proposal, picked with probability $\alpha(\mathsf{o} \to \mathsf{n})$, consists of choosing a new grafting point $P'$ with probability $\alpha_{\text{graft}}(\mathsf{o} \to \mathsf{n})$, followed by a new mutational history for the tree compatible with the tip sequences, picked with probability $\alpha_{\text{mut}}(\mathsf{o} \to \mathsf{n})$. We take $\alpha_{\text{graft}}(\mathsf{o} \to \mathsf{n})$ as given and focus on the second step here.

At one extreme, we could consider proposing a new mutational history for the entire tree via stochastic mapping[18]. The probability $\alpha_{\text{mut}}(\mathsf{o} \to \mathsf{n})$ is then proportional to $\mathcal{G}(\mathsf{n})$, and by construction, the normalization constant is the usual Felsenstein pruning tree likelihood $\mathcal{L}_{\text{tree}}(\mathsf{n})$ (see Section 4). This approach yields the following acceptance probability:

$$p_{\text{acc}} = \min\left[1, \frac{\mathcal{L}_{\text{tree}}(\mathsf{n})\pi_\varphi(\mathsf{n})}{\mathcal{L}_{\text{tree}}(\mathsf{o})\pi_\varphi(\mathsf{o})} \Big/ \frac{\alpha_{\text{graft}}(\mathsf{o} \to \mathsf{n})}{\alpha_{\text{graft}}(\mathsf{n} \to \mathsf{o})}\right], \qquad \text{(full stochastic mapping)}.$$

This is nothing but the standard acceptance probability for SPR moves when using Felsenstein pruning, as implemented in BEAST and other similar tools. In the explicit mutation representation, we thus see that these moves correspond to performing stochastic mapping over the entire tree after every single move, which hints at why this operation is relatively expensive.

At the other extreme, we could consider proposing a new mutational history which is identical to the old one except for the the branch connecting $X$ to $P$ or $P'$. In such a scheme, the sequence at $X$, $P$ and $P'$ are the same in the old and new configurations. By walking in the old configuration along the path from $X$ to $P$ and then to $P'$, we can enumerate the $M(\mathsf{n})$ differences in the sequences of $X$ and $P'$ in time proportional to the number of mutations along that path, which we expect to be very small. An “optimistic” parsimony-inspired scheme for proposing a new mutational history from $P'$ to $X$ would be to omit any mutations in sites where the sequences of $X$ and $P'$ agree, and propose a single mutation where the sequences disagree, distributed uniformly over a time $t_X - t_{P'}$. Such a scheme has:

$$\alpha_{\text{mut}}(\mathsf{o} \to \mathsf{n}) = \left(\frac{1}{t_X - t_{P'}}\right)^{M(\mathsf{n})}, \qquad \text{(optimistic scheme)}. \tag{20}$$

The optimistic scheme works and is efficient as long as the mutational history of the $P$–$X$ branch in the old configuration has at most one mutation per site. In particular, note that the cost of the proposal scales as the number of mutations along the $X$–$P$–$P'$ path, which is typically of order unity, not the genome size, which can easily exceed several thousands for real-world datasets. However, if the $P$–$X$ branch in the old configuration has 2 or more mutations on the same site, then the proposal probability of the reverse move is zero, and the SPR move is always rejected. Hence, the fatal flaw of the optimistic scheme is that, in the absence of more sophisticated moves, the MCMC lacks a mechanism for modifying the tree around a branch with multiple mutations on the same site.

One way to improve the optimistic scheme is to apply stochastic mapping along the single $P'$ to $X$ branch given its fixed endpoints, using the site-specific evolution model for each site (this is the essence of Nielsen’s original SPR move[19]). In that case, the probability of generating the specific configuration for site $\ell$ is proportional to $\mathcal{G}_X^{(\ell)}(\mathsf{n})$, with a normalization constant that can be calculated explicitly in terms of evolution matrix elements:

$$\alpha_{\mathrm{mut}}(\mathsf{o} \to \mathsf{n}) = \frac{\prod_\ell \mathcal{G}_X^{(\ell)}(\mathsf{n})}{\prod_\ell P^{(\ell)}_{s_{P'}^{(\ell)}\, s_X^{(\ell)}}(t_X - t_{P'})}, \qquad \text{(exact stochastic mapping on } P'\text{–}X \text{ branch).}$$

In this scheme, the ratios of site-branch genetic priors $G_X^{(\ell)}(\mathsf{n})/G_X^{(\ell)}(\mathsf{o})$ exactly cancel the ratio of genetic priors $\mathcal{G}(\mathsf{n})/\mathcal{G}(\mathsf{o})$. However, since each site $\ell$ in principle has a different transition rate matrix $Q_{ab}^{(\ell)}$, the stochastic mapping needs to be performed site by site, so the cost of the proposal scales as the genome size. Moreover, the state at $P'$ must be reconstructed at each site, which needs a walk up to the root of the tree. These are both serious flaws.

A simple adaptation tames the high cost of the above scheme. First, note that whenever $\mu(t_X - t_{P'}) \ll 1$, the parsimonious solution of the optimistic scheme dominates regardless of the details of the evolution model. All that is really required is a scheme that exhibits a similar dominance but that also allows, in principle, for any mutational history to be proposed. To this end, we use a simple Jukes-Cantor model with a fictitious mutation rate $\tilde{\mu} = \lambda(X)/L$ and no site-rate heterogeneity. Let $P_{=}^{JC}(t)$ and $P_{\neq}^{JC}(t)$ be the evolution matrix elements of this model when the endpoints respectively do or do not match, given by the well-known formulas

$$P_{=}^{JC}(t) = (1/4)(1 + 3e^{-(4/3)\tilde{\mu}t}) \approx 1 - \tilde{\mu}t;$$
$$P_{\neq}^{JC}(t) = (1/4)(1 - e^{-(4/3)\tilde{\mu}t}) \approx (\tilde{\mu}/3)t.$$

The approximations hold in the limit of $\tilde{\mu}t \ll 1$. Since we use the same model for every site, and multiple mutations in sites with equal states at the endpoints of a short branch are very rare, we can perform the stochastic mapping using this model roughly in time proportional to $M(\mathsf{n})$ (see below). The final proposal probability is given in terms of $M(\mathsf{n})$, the actual number $\mathcal{M}(\mathsf{n})$ of mutations proposed, and the above evolution matrix elements, as follows:

$$\alpha_{\mathrm{mut}}(\mathsf{o} \to \mathsf{n}) = \frac{e^{-\tilde{\mu}L(t_X - t_{P'})}(\tilde{\mu}/3)^{\mathcal{M}(\mathsf{n})}}{[P_{\neq}^{JC}(t_X - t_{P'})]^{M(\mathsf{n})}[P_{=}^{JC}(t_X - t_{P'})]^{L - M(\mathsf{n})}},$$
(approximate stochastic mapping on $P'$–$X$ branch).

Note that in the limit where $\tilde{\mu}(t_X - t_{P'}) \ll 1$ and $\mathcal{M}(\mathsf{n}) = M(\mathsf{n})$, the above proposal probability reduces to that of the optimistic scheme (Eq. (20)), which is independent of $\tilde{\mu}$.

#### 6.2 Efficient Jukes-Cantor stochastic mapping in short branches

We first consider the simpler problem of performing a stochastic mapping on a branch of length $\tau$ with a Jukes-Cantor model having mutation rate $\tilde{\mu}$, and where the starting and ending state at all $L$ sites is `A`. For sufficiently small $\tilde{\mu}\tau$, we expect this procedure to result in no mutations at all with overwhelming likelihood, so we arrange the algorithm to produce that result efficiently when necessary. At the end of this section, we discuss the refinements needed to implement stochastic mapping when a few sites have different starting and ending states, and the other sites' equal starting and ending states are not all `A`.

Conceptually, proposing $L$ independent trajectories from `A` to `A` over a time $\tau$ proceeds via $L$ repetitions of a variation of Nielsen's rejection sampling algorithm[18]. A trajectory is first proposed as follows: the number of mutations on the trajectory is drawn from a Poisson distribution with mean $\tilde{\mu}\tau$, with times distributed uniformly from 0 to $\tau$, and with identities chosen at random (subject to the end state of one mutation matching the start state of the next mutation). The trajectory is accepted if the state at time $\tau$ is `A`, otherwise it is rejected and the algorithm begins anew. The course of the algorithm is generally as follows. If zero mutations are proposed, which happens with probability $p_0 = e^{-\tilde{\mu}\tau}$, the trajectory is empty and is immediately accepted. If one mutation is proposed, which happens with probability $p_1 = \tilde{\mu}\tau e^{-\tilde{\mu}\tau}$, the trajectory necessarily ends in a state other than `A` and is rejected. Otherwise, the acceptance or rejection of the trajectory depends on the details of the proposed mutation identities.

Executing the above procedure directly is expensive and wasteful. Overwhelmingly, the above procedure simply proposes and rejects a one-mutation trajectory $k$ times, with $k = 0$ almost always, and then proposes a zero-mutation trajectory, which is accepted. This observation suggests that we *simulate* the above procedure by deciding up front whether or not it ends in an acceptance of an empty trajectory before any trajectory with 2 or more mutations is proposed. The probability that no trajectory with 2 or more mutations is proposed before a zero-mutation proposal is accepted is

$$p_* = \sum_{k=0}^{\infty} p_1^k p_0 = \frac{p_0}{1 - p_1}$$

Hence, we can directly produce an empty trajectory for a site with probability $p_*$. With probability $1 - p_*$, we instead directly make a proposal with 2 or more mutations: if it ends in `A`, we accept it; otherwise, we repeat this procedure. Finally, instead of doing this independently for each site, we can draw a number of sites from $\mathrm{Geom}(1 - p_*)$ distribution to determine for how many sites we trivially propose an empty trajectory before we have to resort to more detailed sampling. The final algorithm is presented in Algorithm 1, which we note must be implemented carefully to avoid numerical issues with small $p_*$ and large numbers drawn from $\mathrm{Geom}(1 - p_*)$.

**Algorithm 1** Efficient stochastic mapping for a branch of length $\tau$ using a Jukes-Cantor model with mutation rate $\tilde{\mu}$, with starting and ending states `A` at all $L$ sites.

$p_0 \leftarrow e^{-\tilde{\mu}\tau}$
$p_1 \leftarrow \tilde{\mu}\tau e^{-\tilde{\mu}\tau}$
$p_* \leftarrow p_0/(1 - p_1)$
$\ell \leftarrow 0$
**while** $\ell < L$ **do**
    $\Delta \sim \mathrm{Geom}(1 - p_*)$
    $\ell \leftarrow \ell + \Delta$
    **if** $\ell < L$ **then**
        $n \sim \mathrm{KPois}(\tilde{\mu}\tau, 2)$ ▷ $\mathrm{KPois}(\lambda, k)$ is the restriction of $\mathrm{Pois}(\lambda)$ to at least $k$ events.
        $s_0 \leftarrow$ `A`
        **for** $i \leftarrow 1$ to $n$ **do**
            $s_i \sim \mathrm{DUnif}(\{\mathtt{A}, \mathtt{C}, \mathtt{G}, \mathtt{T}\} \setminus \{s_{i-1}\})$
        **end for**
        **if** $s_n =$ `A` **then** ▷ Acceptance of non-empty mutational history
            $t_1 \leq \ldots \leq t_n \sim \mathrm{Unif}(0, \tau)$
            Output mutations $s_0 \ell s_1$ at $t_1$, …, $s_{n-1} \ell s_n$ at $t_n$
            $\ell \leftarrow \ell + 1$
        **else**
            (Rejection sampling starts anew at site $\ell$, not $\ell + 1$).
        **end if**
    **end if**
**end while**

For a typical genomic epidemiology tree of a SARS-CoV-2-like virus, we have $\tilde{\mu} \approx 10^{-3}\,\mathrm{yr}^{-1}$, $L \approx 30\,000$ and $\tau \approx 2\,\mathrm{weeks}$, whereby $1 - p_* \approx 7 \times 10^{-10}$. The probability that the initial draw from $\mathrm{Geom}(1 - p_*)$ doesn't immediately skip over all $L$ sites is $1 - (1 - p_*)^L \approx 2 \times 10^{-5}$. Even for a much longer branch with $\tau = 1\,\mathrm{year}$, this probability only rises to around 1.5 %. Hence, overwhelmingly, this procedure immediately produces an empty mutational trajectory for all $L$ sites after drawing a single random number.

Two further refinements are needed to apply the above ideas to complete the proposal of the mutational history between $P'$ and $X$. First, we need to treat differently the $M(\mathsf{n})$ sites where the states at $P'$ and $X$

disagree. Since these are few in number, and their discovery unearths the actual states of that site at $P'$ and $X$, we apply a small variation on Nielsen's rejection sampling scheme directly, site by site: we repeatedly choose a Poisson-distributed number of mutations $n$, subject to $n \geq 1$, produce random mutation identities and times as above, and accept the proposal only if the start and end states agree with the actual states at $P'$ and $X$. When we then use Algorithm 1 to effect a stochastic mapping of the remaining sites, if we stop at a site $\ell$ that where $P'$ and $X$ differ, we immediately skip over this site; this is equivalent to first executing stochastic mapping over all $L$ sites, then filtering out the mutations on the $M(\mathsf{n})$ sites where the states at $P'$ and $X$ differ, which itself is equivalent to performing the detailed stochastic mapping only on the $L - M(\mathsf{n})$ sites where starting and ending states are equal. Finally, in the unlikely case that this stochastic mapping does introduce mutations in sites where $P'$ and $X$ agree, we must discover the actual start and end state of the site, which is unlikely to be `A`, by looking for the nearest mutation at that site on the path from $P'$ to the root of the tree; if no such mutation exists, we read off the state from the reference sequence. With the start and end states known, we then cyclically rotate the states on the proposed mutational history until the start and end states match the known state. For example, if the start and end states of site 1 are `G` and the proposed mutational history was `A1G`, followed by `G1T`, followed by `T1A`, the cyclic rotation modifies the proposal to `G1A`, followed by `A1C`, followed by `C1G`.

#### 6.3 SPR moves involving root changes but no missing data

An SPR move can cause the root of the tree to change if $P$ is the root of the old configuration $\mathsf{o}$ and/or if $P'$ is the root of the new configuration $\mathsf{n}$. We focus here on the latter scenario, as the treatment of the former one is completely analogous. In this section, we are still restricted to sample sequences with no missing data.

Since an EMAT does not explicitly represent the mutational history of the lineage ancestral to the root node, the SPR proposal needs to include these details. Concretely, let $S$ be the sibling of $X$ in the old configuration $\mathsf{o}$, and let $S'$ be the sibling of $X$ in the new configuration $\mathsf{n}$, as depicted in Figure 16. Delphy's SPR move in this case does not consider the sequence at $P'$ to be fixed; instead, the mutational history along the entire $X$–$P'$–$S'$ path is proposed afresh, subject to the sequences at $X$ and $S'$ being fixed. To do this, we exploit the time-reversibility of the Jukes-Cantor model to propose a single trajectory of length $\tau' = (t_X - t_{P'}) + (t_{S'} - t_{P'})$ with the end sequences of $X$ and $S'$. Although we're exploiting the time-reversibility of the Jukes-Cantor model, note that the actual evolution model used in the posterior distribution need not be time-reversible. The proposal probability given by:

$$\alpha_{\text{mut}}(\mathsf{o} \to \mathsf{n}) = \frac{e^{-\tilde{\mu} L \tau'} (\tilde{\mu}/3)^{\mathcal{M}(\mathsf{n})}}{[P^{JC}_{\neq}(\tau')]^{M(\mathsf{n})} [P^{JC}_{=}(\tau')]^{L-M(\mathsf{n})}}, \qquad \text{(when } P' \text{ is the root of the new configuration).}$$

Note that the ratio of genetic priors in the acceptance probability (Eq. (19)) includes a ratio of root priors between the old and new configurations (see below for details). This ratio can be evaluated efficiently in time proportional to the number of mutations proposed along the $P'$ to $S'$ branch in the new configuration and/or the $P$ to $S$ branch in the old configuration.

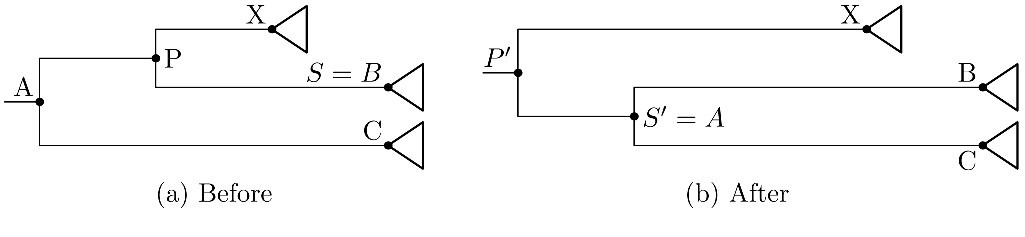


**Supplementary Figure 16:** An SPR move that changes the root. Whereas the mutational history along the $A$–$B$ path can remain fixed, a new mutational history must be proposed for the entire $X$–$P'$–$S'$ path.

A useful viewpoint for the remainder of this section is to view the root of an EMAT as missing data *upstream* at every site, as opposed to downstream at some sites towards the tips, so that the sequence at $P'$

is not highly constrained by the three branches that impinge on it. The way the above scheme deals with such missing data is by extending the path over which mutations are proposed from $X$–$P'$ to $X$–$P'$–$S'$ for all sites, and stopping at $S'$ because the sequence at $S'$ *is* strongly constrained by those of the two children of $X$ and $S'$. This viewpoint is expanded in the sections below on how Delphy deals with missing data.

#### 6.4 SPR moves involving missing data but no root changes

Consider the situations in Figures 17(a) and (b) involving a dataset with $L = 100$ sites. The subtree rooted at $X$ is detached at $P$ from the rest of the tree, with the entire subtree, its mutational history and the sequence at $X$ held fixed in sites where at least one downstream tip is informative. In the original configuration, the branch leading to $X$ is decorated with missations on sites 10–20, while the one leading to $X$'s sibling $S$ is decorated with missations on sites 30–40. These missations imply that at least one of the tips downstream of $X$ *is* informative for sites 30–40, and analogously, at least one of the tips downstream of $S$ is informative for sites 10–20 (otherwise, the corresponding missations would appear further upstream). When $X$ is detached, the points immediately upstream of $P$ no longer have downstream tips that are informative for sites 30–40. It follows that the corresponding missations will be forced to appear further upstream. Any mutations at sites 30–40 on the branch above $P$, such as the `G33T`, must not appear on the tree after $X$ is detached. This process continues recursively towards the root: while in the original configuration, $P$'s sibling has missations on sites 35–45, the subset of those missations on sites 35–40 are forced to appear further upstream, removing the `C37G` mutation. This example captures the essential complication of implementing SPR moves in the presence of missing data. In a very concrete way, parts of the path from $X$ all the way to the root are "linked" to the attachment of $X$ at $P$. SPR moves in the presence of missing data must consider these paths and the mutations on them in their proposal and evaluation of their Metropolis-Hastings factor.

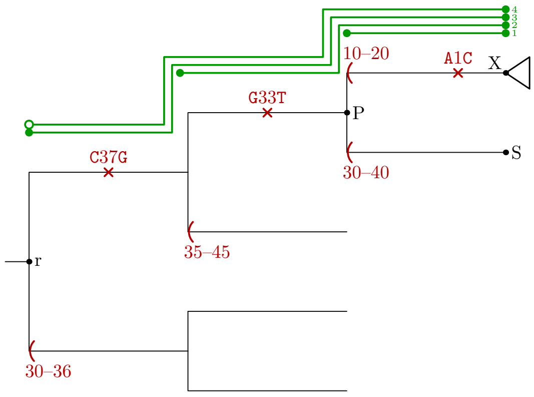


(a) Before pruning $X$

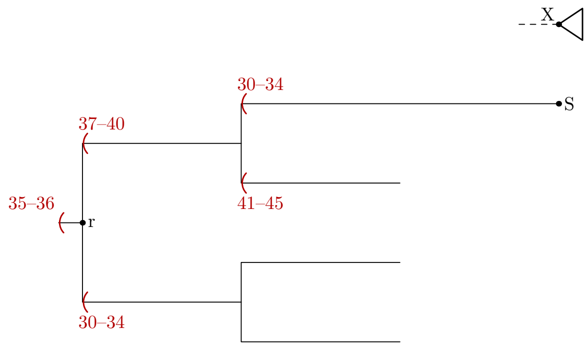


(b) After pruning $X$

**Supplementary Figure 17:** The pruning step of an SPR move in the presence of missations. The paths labeled 1 through 4 all need to be treated differently. See text for details.

To discuss the move in detail, we first introduce helpful notation. Let $\phi_i := \varphi^i(X)$ be the $i^{\text{th}}$ ancestor of $X$, with $i = 0, \ldots, I$. It follows that $\phi_0$ is $X$ and $\phi_1$ is $P$; as for $I$, it is chosen such that $\phi_I$ is the root node. In this section, we assume $X$ is not a child of the root, so that $I$ is at least 2; we relax that restriction in the next section. For $i$ positive, denote by "branch $i$" the branch from $\phi_i$ to $\phi_{i-1}$. Likewise, denote by "path $i$" the path from $\phi_i$ to $X$, which has length $T_i$ given by $T_i = t_X - t_{\phi_i}$. Next, denote by $S_i$ the child of $\phi_i$ that is not on path $i$. Denote by $\mathcal{N}_i$ the missations on the branch from $\phi_i$ to $S_i$.

We first identify the sets of sites $\tilde{\mathcal{L}}_i$ along branch $i$ that are "linked" to the attachment of $X$ at $P$. We say that these sites are "warm" on branch $i$. Clearly

$$\tilde{\mathcal{L}}_i = \tilde{\mathcal{L}}_{i-1} \cap \mathcal{N}_{i-1}, \qquad (2 \le i \le I + 1).$$

The base case of this recurrence is $\tilde{\mathcal{L}}_1 = \xi(X)$, i.e., the set of sites present at $X$. We discuss the meaning of $\tilde{\mathcal{L}}_{I+1}$ below, but for now assume that it is empty. For example, in the scenario of Figure 17(a), but with no missations on the lower branch from the root, we have $\tilde{\mathcal{L}}_1 = [1,9] \cup [21,100]$, $\tilde{\mathcal{L}}_2 = [30,40]$, $\tilde{\mathcal{L}}_3 = [35,40]$ and $\tilde{\mathcal{L}}_4 = \emptyset$. We re-examine the full scenario with $\tilde{\mathcal{L}}_4 \neq \emptyset$ at the end of this section. In general, calculating $\tilde{\mathcal{L}}_1$ involves an undesirable walk from $X$ all the way to the root to enumerate all the sites for which there is a missation on path $I$. We can almost always avoid this calculation by noting that if $\ell \in \mathcal{N}_1$, it is necessarily present along the branch $P$–$X$, and is thus a member of $\tilde{\mathcal{L}}_1$. Hence, $\tilde{\mathcal{L}}_2 = \mathcal{N}_1$, and in general,

$$\tilde{\mathcal{L}}_i = \mathcal{N}_{i-1} \cap \cdots \cap \mathcal{N}_1, \qquad (2 \leq i \leq I+1).$$

These subsets, represented as sorted lists of intervals, can be calculated efficiently in time proportional to the number of such intervals because missations are themselves represented as sorted lists of intervals. We'll see below that we usually only need to know the size of $\tilde{\mathcal{L}}_1$, not its exact contents. We also note that very often, the above recurrence often reaches a point where $\mathcal{L}_i = \emptyset$ for small $i$, so we often don't need to walk more than a few branches upstream of $X$ in the calculations below (in the extreme case, when there are no missations on the branch from $P$ to $S$, we have $\mathcal{L}_2 = \emptyset$ already, and the entire scheme here reduces to that presented in Section 6.1).

We further define a set of "hot" sites $\mathcal{L}_i$ on path $i$ as follows:

$$\mathcal{L}_i := \tilde{\mathcal{L}}_i \setminus \tilde{\mathcal{L}}_{i+1}, \qquad (1 \leq i \leq I).$$

For example, in Figure 17(a) but without the missations on the lower branch from the root, we have $\mathcal{L}_1 = [1,9] \cup [21,29] \cup [41,100]$, $\mathcal{L}_2 = [30,34]$ and $\mathcal{L}_3 = [35,40]$. The sets $\mathcal{L}_1, \ldots, \mathcal{L}_I$ partition the set of sites $\tilde{\mathcal{L}}_1$ that are present at $X$ such that a site $\ell$ is in $\mathcal{L}_i$ if it is warm on path $i$ but not on path $i+1$. When viewed in terms of the pruned site-trees (Figure 13), path $i$ corresponds to the branch upstream of $X$ for the site-$\ell$ pruned tree for which $\ell$ is hot in path $i$.

The SPR move proposal is as follows. For every path $i$, we hold the sequences and mutational histories of sites $\mathcal{L}_i$ fixed everywhere except in the interior of path $i$. We then use the scheme described in Section 6.2 to propose new mutational histories along the entirety of path $i$, with length $T_i$, subject to fixed end states, with fake mutation rate $\tilde{\mu} = \lambda_X/|\xi(X)|$. The set of sites that are hot in path $i$ and have different end states is small and easy to construct while calculating $\mathcal{L}_i$. For the remaining sites, we expect the probability of even a single non-parsimonious mutation to appear to be very small, so we simply run Algorithm 1 over all $L$ sites, then filter the result to include only sites in $\mathcal{L}_i$; in the exceedingly unlikely case that the result is not empty, we walk up the tree to deduce the actual state of site $i$ at $\phi_i$, as in Section 6.2. The resulting proposal probability is

$$\alpha_{\text{mut}}(\mathsf{o} \to \mathsf{n}) = \prod_{i=1}^{I} \frac{e^{-\tilde{\mu}|\mathcal{L}_i|T_i} (\tilde{\mu}/3)^{\mathcal{M}_i(\mathsf{n})}}{[P^{JC}_{\neq}(T_i)]^{M_i(\mathsf{n})} [P^{JC}_{=}(T_i)]^{|\mathcal{L}_i| - M_i(\mathsf{n})}},$$

(approximate stochastic mapping on hot sites in paths above $X$).

In analogy to the situation without missing data, we have that $|\mathcal{L}_i|$ is the number of hot sites on path $i$, $T_i$ is the path's length, $\mathcal{M}_i(\mathsf{n})$ is the actual number of proposed mutations on hot sites of path $i$ and $M_i(\mathsf{n})$ is the number of hot sites on path $i$ whose start and end state differ. As hinted above, this formula requires us to know $|\mathcal{L}_1| = |\xi(X)|$, i.e., the number of sites present at node $X$, but rarely its exact membership, which is expensive to calculate (the exception occurs when a hot site on path 1 accumulates mutations when the start and end states are equal, which we must filter to exclude sites that are missing at $X$). We keep track $|\xi(X)|$ for all nodes $X$ separately, and update it progressively during all tree topology changes.

To calculate the ratio of genetic priors in the new and old configurations, we actually calculate a ratio of ratios of the genetic priors between each configuration with $X$ detached or attached (the situations in Figures 17(a) and (b)). Denote by $\mathcal{G}'(\mathsf{n})$ (respectively, $\mathcal{G}'(\mathsf{o})$) this latter ratio for the new (respectively, old) configuration. Since only the hot sites along each path $i$ change when detaching $X$, the calculation resembles the analogous calculation when there is no missing data. In particular,

$$\mathcal{G}' = \prod_{i=1}^{I} \left[ e^{-\int_{x \in \text{ path } i} \lambda_i(x)\,dx} \prod_{(a,\ell,b)\in\mathcal{M}_i} Q^{(\ell)}_{ab} \right].$$

Here, $\lambda_i(x) = \sum_{i\in\mathcal{L}_i} Q^{(\ell)}_{s_i^{(\ell)}(x)}$, that is, the contribution to $\lambda(x)$ from the hot sites of path $i$, while $\mathcal{M}_i$ are the mutations on hot sites along path $i$. Calculating this quantity given $\mathcal{L}_i$, represented as a sorted list of intervals, is efficient because, as noted earlier, we keeping a cumulative sum of all terms $Q^{(\ell)}_{s_\perp^{(\ell)}}$ for the reference sequence, and the reference sequence rarely changes. The value of $\lambda_i(\phi_i)$ if the sequence at $\phi_i$ were the reference sequence follows straightforwardly. To adjust it to the actual sequence at $\phi_i$, we note that all sites which are hot on path $i$ are also in the missations $\mathcal{N}_i$, and those missations record state differences with the reference sequence (of which there are usually few) explicitly.

For completeness, the acceptance probability of the SPR move in the presence of missing data is

$$p_{\text{acc}} = \min\left[1, \frac{\mathcal{G}'(\mathsf{n})}{\mathcal{G}'(\mathsf{o})} \Big/ \frac{\alpha_{\text{mut}}(\mathsf{o}\to\mathsf{n})\alpha_{\text{graft}}(\mathsf{o}\to\mathsf{n})}{\alpha_{\text{mut}}(\mathsf{n}\to\mathsf{o})\alpha_{\text{graft}}(\mathsf{n}\to\mathsf{o})}\right].$$

One final complication arises when there are sites $\ell$ for which only the tips downstream of $X$ are informative. These are precisely the members of $\mathcal{L}_{I+1}$, which we assumed empty above. One possibility would be to *define* $\mathcal{L}_{I+1} := \emptyset$, which leads to proposals where the state of the root at these sites is fixed. If we proceeded thus, and the current states of a site $\ell$ at the root and at $X$ were different, then the SPR move would have no mechanism of proposing a mutation-free path for site $\ell$, which would change the state of site $\ell$ at the root to its state at $X$; this is unfortunate because in the near-parsimonious limit, that is overwhelmingly the most likely configuration. So, instead, we treat sites in $\tilde{\mathcal{L}}_{I+1}$ specially. First, we declare them to be hot by definition, that is:

$$\mathcal{L}_{I+1} := \tilde{\mathcal{L}}_{I+1}.$$

Then, the SPR mutational history proposal for a site $\ell \in \mathcal{L}_{I+1}$ consists of a simple open-ended trajectory with end state $s_X^{(\ell)}$ (in Figure 17(a), this is indicated by the open circle at the end of path 4). We construct this trajectory backwards, starting at time $t_X$ and repeatedly drawing exponentially distributed times between successive mutations at a rate $\tilde{\mu}$ until the next mutation time precedes the root time. For each mutation time, we pick a corresponding site $\ell$ uniformly over all sites $L$, and discard it unless the site is in $\mathcal{L}_{I+1}$. On the rare occasions when one mutation in $\mathcal{L}_{I+1}$ is proposed, the reference sequence, the mutations on branch $X$ and the missations $\mathcal{N}_1$ are used to reconstruct $s_X^{(\ell)}$, and then a mutation identity is chosen uniformly at random. The state at $\ell$ is recorded and the algorithm proceeds further until the next proposed mutation occurs at a time $t < t_r$. The proposal probability gets an additional term, as follows:

$$\alpha'_{\text{mut}}(\mathsf{o}\to\mathsf{n}) = \alpha_{\text{mut}}(\mathsf{o}\to\mathsf{n}) \cdot e^{-\tilde{\mu}|\mathcal{L}_{I+1}|T_I} (\tilde{\mu}/3)^{\mathcal{M}_{I+1}(\mathsf{n})}.$$

The time of the open-ended trajectory is $T_I$ (there is no $T_{I+1}$) and there is no denominator in the extra factor because the trajectory is open-ended. Conversely, the genetic prior ratio $\mathcal{G}'$ is modified to reflect the change in root state for each $\ell \in \mathcal{L}_{I+1}$ to that at $X$ once $X$ is detached:

$$\mathcal{G}'' = \mathcal{G}' \cdot e^{-\int_{x\in\text{path } I} \lambda_{I+1}(x)\,dx} \prod_{(a,\ell,b)\in\mathcal{M}_{I+1}} Q^{(\ell)}_{ab} \cdot (\pi_a^{(\ell)}/\pi_b^{(\ell)}).$$

We close this section by noting that the book-keeping burden of updating missations and the differences between their "from" states and the reference sequence as trees are pruned and regrafted elsewhere is nontrivial but tractable. Since the details of such book-keeping are not particularly illuminating, we omit them here.

#### 6.5 SPR moves involving root changes and missing data

Consider the situation in Figure 18. Here, a subtree rooted at $X$ is directly attached to the root of the tree. Detaching $X$ also removes the branch from the root to its sibling $S$. Doing so is complicated by the presence of missations on either the $P$–$X$ or the $P$–$S$ branch. However, unlike in the previous section, the missations upstream of the $X$, $P$ and $S$ nodes suffice to build the sets of sites with no informative tips downstream of these three nodes. Hence, we can implement a variant of the scheme of the previous section in terms of three paths: a path $S$–$P$–$X$ with fixed endpoints, and two open-ended paths $P$–$X$ and $P$–$S$.

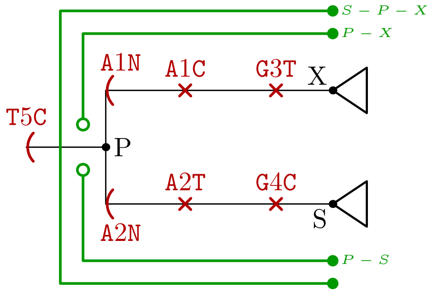


**Supplementary Figure 18:** The situation preceding the pruning step of an SPR move when the pruned subtree is a direct of the root and there are missations around the root. The paths labeled $S - P - X$, $P - X$ and $P - S$ all need to be treated differently. See text for details.

Denote by $\mathcal{N}_X$, $\mathcal{N}_P$ and $\mathcal{N}_S$ the missations on the branches ending at nodes $X$, $P$ and $S$, respectively. Denote by $\mathcal{P}_i$ the set of sites present at node $i$, i.e., for which there is at least one informative tip downstream of $i$. Clearly, we have

$$\begin{aligned}\mathcal{P}_P &= \{1, \ldots, L\} \setminus \mathcal{N}_P;\\ \mathcal{P}_X &= \mathcal{P}_P \setminus \mathcal{N}_X;\\ \mathcal{P}_S &= \mathcal{P}_S \setminus \mathcal{N}_S.\end{aligned}$$

Each of the sites present at $P$ is next classified as a hot site in one of the paths $S$–$P$–$X$, $P$–$X$ and $P$–$S$. Concretely,

$$\begin{aligned}\mathcal{L}_{S-P-X} &= \mathcal{P}_P \setminus (\mathcal{N}_X \cup \mathcal{N}_S);\\ \mathcal{L}_{P-X} &= \mathcal{N}_S;\\ \mathcal{L}_{P-S} &= \mathcal{N}_X.\end{aligned}$$

The SPR mutation history proposal treats each of these three paths separately, as described below.

For the $S$–$P$–$X$ path, we again use a variation of the scheme described in Section 6.2 to propose mutations for the sites in $\mathcal{L}_{S-P-X}$ according to a Jukes-Cantor model with mutation rate $\tilde{\mu}$ over a time $T_{S-P-X} = (t_X - t_P) + (t_S - t_P)$, with fixed endpoints. By walking over the mutations along the path $S$–$P$–$X$ (which are typically few in number), we identify all sites whose starting and ending states differ, as well as what those states are. For the remaining sites, if any mutations are proposed, we can reconstruct the state at $X$ by looking directly at the reference sequence and the mutations above $P$. Since a Jukes-Cantor model is time-reversible and has all root prior frequencies of 1/4, no complications arise if the proposed root states differ from those at $X$. Mutations with times in the range $[0, t_S - t_P)$ are mapped with reversed identities to the range $t_S$ to $t_P$ on the $P$–$S$ branch, while those with times in the range $[t_S - t_P, T_{S-P-X})$ are mapped with their proposed identities to the range $t_P$ to $t_X$ on the $P$–$X$ branch. The part of the proposal probability due to the $S$–$P$–$X$ path is given by

$$\alpha_{\text{mut, } S\text{–}P\text{–}X}(\mathsf{o} \to \mathsf{n}) = \frac{e^{-\tilde{\mu}|\mathcal{L}_{S-P-X}|T_{S-P-X}}(\tilde{\mu}/3)^{\mathcal{M}_{S-P-X}(\mathsf{n})}}{[P^{JC}_{\neq}(T_{S-P-X})]^{M_{S-P-X}(\mathsf{n})}[P^{JC}_{=}(T_{S-P-X})]^{|\mathcal{L}_{S-P-X}|-M_{S-P-X}(\mathsf{n})}},$$

(approximate stochastic mapping on hot sites of the $S$–$P$–$X$ path).

As above, $M_{S-P-X}(\mathsf{n})$ is the actual number of mutations proposed, while $\mathcal{M}_{S-P-X}(\mathsf{n})$ is the number of sites in $\mathcal{L}_{S-P-X}$ where the states at $X$ and $S$ differ. The ratio of genetic priors before and after detaching $X$ is as in the previous section, with a slight change to reflect that the root sequence after detachment is that at $S$:

$$\mathcal{G}'_{S-P-X} = e^{-\int_{x\in S-P-X} \lambda_{S-P-X}(x)\,dx} \prod_{(a,\ell,b)\in\mathcal{M}_{S-P-X}} Q^{(\ell)}_{ab} \prod_{(a,\ell,b)\in\mathcal{M}_{S-P-X} \text{ on } P\text{–}S \text{ branch}} (\pi^{(\ell)}_a/\pi^{(\ell)}_b).$$

In analogy to the open-ended paths of the previous section, the mutational history proposal for the $P$–$X$ path is an open-ended proposal for sites in $\mathcal{L}_{P-X}$ ending in the current state at $X$. As above, this leads to a proposal probability of

$$\alpha_{\text{mut, } P\text{–}X}(\mathsf{o}\to\mathsf{n}) = e^{-\tilde\mu|\mathcal{L}_{P-X}|T_{P-X}}(\tilde\mu/3)^{\mathcal{M}_{P-X}(\mathsf{n})},$$

(approximate stochastic mapping on hot sites of the open-ended $P$–$X$ path).

The corresponding genetic ratio reflects that the root state after detachment for these sites is set to the state at $X$:

$$\mathcal{G}'_{P-X} = e^{-\int_{x\in P-X} \lambda_{P-X}(x)\,dx} \prod_{(a,\ell,b)\in\mathcal{M}_{P-X}} Q^{(\ell)}_{ab}\cdot(\pi^{(\ell)}_a/\pi^{(\ell)}_b).$$

The treatment of the $P$–$S$ path is exactly analogous.

The final proposal probability for the mutations is given by

$$\alpha_{\text{mut}}(\mathsf{o}\to\mathsf{n}) = \alpha_{\text{mut, } S\text{–}P\text{–}X}(\mathsf{o}\to\mathsf{n})\cdot\alpha_{\text{mut, } P\text{–}X}(\mathsf{o}\to\mathsf{n})\cdot\alpha_{\text{mut, } P\text{–}S}(\mathsf{o}\to\mathsf{n}).$$

Similarly, the final ratio of genetic priors before and after grafting is given by

$$\mathcal{G}'(\mathsf{n}) = \mathcal{G}'_{S-P-X}(\mathsf{n})\cdot\mathcal{G}'_{P-X}(\mathsf{n})\cdot\mathcal{G}'_{P-S}(\mathsf{n}).$$

#### 6.6 Concrete SPR moves

In the current version of Delphy, we have implemented two of BEAST's SPR moves, subtree slide and Wilson-Balding, as well as a more sophisticated one we call "mutation-directed SPR" (mdSPR). For completeness, we briefly describe the first two to present explicit expressions for $\alpha_{\text{graft}}(\mathsf{o}\to\mathsf{n})$. We then describe mdSPR in detail.

For subtree slide, we pick $X$ as a random non-root node. We then find a point $P'$ at a time $\delta$ downstream or upstream of $X$'s parent $P$, where $\delta\sim\mathcal{N}(0,\Delta^2)$ and $\Delta = \lambda_X^{-1}/2$ (this choice makes it possible for the path from $P$ to $P'$ to sometimes have one mutation, but rarely more). When $\delta$ is negative, the choice of $P'$ is unambiguous. Conversely, in the reverse move and when $\delta$ is positive, there are $K$ points on the tree to choose from that are descended from $P$, active at time $t_P+\delta$ and exist before $t_X$. If $K=0$, we reject the move. Otherwise, we choose one of these uniformly at random. With the above considerations, the ratio of proposal probabilities is:

$$\frac{\alpha_{\text{graft}}(\mathsf{n}\to\mathsf{o})}{\alpha_{\text{graft}}(\mathsf{o}\to\mathsf{n})} = \begin{cases} K, & \delta>0; \\ 1/K, & \delta\le 0, \end{cases} \qquad \text{[Subtree slide]}.$$

For the Wilson-Balding move, we pick $X$ randomly such that neither $X$ nor $P$ is the root. We then pick what will be $X$'s sibling after the move, $S'$, randomly, subject to the following condition. Let $G=\varphi(P)$ and $G'=\varphi(S')$. First, the path $G'$–$S'$ must have no overlap with the path $G$–$S$ (including the endpoints). Second, we need $t_{G'}>t_X$, as otherwise we cannot attach $X$ to the branch $S'$ with a positive-length branch. If either of these conditions is not met, we reject the move. Otherwise, we pick the new attachment time $t_{P'}\sim U(t_{G'},\min[t_X,t_{S'}])$. With the above considerations, the ratio of proposal probabilities is:

$$\frac{\alpha_{\text{graft}}(\mathsf{n}\to\mathsf{o})}{\alpha_{\text{graft}}(\mathsf{o}\to\mathsf{n})} = \frac{\min[t_X,t_{S'}]-t_{G'}}{\min[t_X,t_S]-t_G}, \qquad \text{[Wilson-Balding]}.$$

###### 6.6.1 Mutation-directed SPR moves (mdSPR)

In Figure 19, the possible grafting points for the SPR move that prunes the subtree rooted at $X$ are classified into regions along each branch, demarcated from each other by an intervening mutations; all points in a single region are in the same branch and have the same sequence. It is easy to efficiently calculate the number of sequence differences between the sequence at $X$ and every point in each region, by a simple local walk of the tree: these numbers are written in red inside parentheses. These annotations clearly suggest an effective way to propose a new grafting point for $X$: overwhelmingly propose points where those differences are minimized. Indeed, the Wilson and Balding's original move implemented a crude rendering of this idea in the context of microsatellites[24]. Zhang, Huelsenbeck and Ronquist's pSPR moves[28], implemented in Mr. Bayes, similarly make use of a parallel parsimony tree to guide move proposals. And matOptimize, which pursues strict parsimony improvements, would *always* pick one of the points with minimal sequence difference, with ties broken arbitrarily.

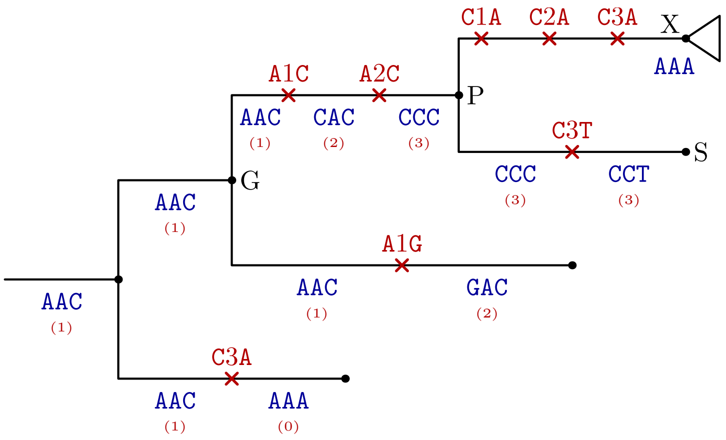


**Supplementary Figure 19:** Setup for an SPR move. A subtree rooted at $X$ will be detached from the remaining tree, where it is currently grafted at $P$. The sequence at $X$ is AAA, while the sequence at $P$ is CCC. Each branch is divided by its mutations into distinct regions, here annotated with their sequence (blue) and the number of differences between that sequence and the sequence at $X$ (red, in parentheses).

Our concrete proposal below, which we call "mutation-directed SPR moves" (mdSPR), exploits the EMAT representation that Delphy already uses as the guide to making SPR proposals. The guide is the "best" one in the sense that a pure mdSPR move reduces to Gibbs sampling when the evolution model is Jukes-Cantor, the mutation rate is low enough that all branches are nearly parsimonious, there is no site-rate heterogeneity, and there is no missing data. In the more general scenario, mdSPR generalizes in a disciplined way the intuition that small changes to the tree are likely to result in a plausible new tree, but that sometimes large-scale changes are essential to correctly sample all plausible trees, not just a subset of closely related ones. While a long enough sequence of local SPR moves are in principle sufficient to turn any tree topology into any other, mdSPR moves ensure that large-scale rearrangement are possible without going through extremely improbable intermediate states. As we describe below, mdSPR moves provide a clear rationale for deciding what constitutes a local move, what balance to strike between local and global moves, how to choose global rearrangements that are likely to result in posterior increases.

To proceed, we restate the observation in Eq. (4) that the posterior of Eq. (1) simplifies drastically when specialized to a Jukes-Cantor substitution model with no site rate heterogeneity and there is no missing data:

$$\mathcal{G}_{JC}(\mathcal{T}) \propto e^{-\mu L T} \cdot \left(\frac{\mu}{3}\right)^{M}. \tag{21}$$

As before, $T$ is the total branch length and $M$ is the total number of mutations. The first factor of Eq. (4), equal to $(1/4)^L$, has been subsumed into the constant of proportionality.

Denote by $N(x)$ the number of differences between the sequences at $X$ and an arbitrary point $x$ on the tree. The above picture suggests that, to a good approximation, the posterior ratio $\mathcal{P}(\mathsf{n})/\mathcal{P}(\mathsf{o})$, is given by

$$\frac{\mathcal{P}(\mathsf{n})}{\mathcal{P}(\mathsf{o})} \approx \frac{\exp[-\mu L(t_X - t_{P'})] \cdot [\mu/3]^{N(P')}}{\exp[-\mu L(t_X - t_P)] \cdot [\mu/3]^{N(P)}}.$$

Unlike above, here, $L = |\xi(X)|$ is the number of sites with informative tips downstream of $X$ and $\mu$ is an effective site mutation rate, given by $\mu = \lambda(X)/L$. The approximation would be exact only if the substitution model were indeed Jukes-Cantor, there were indeed no missing data, and there were exact cancellation in the ancestry priors. Nevertheless, intuitively, the approximation captures the dominant factors in the posterior ratio. We can thus use it to make a proposal, and thus obtain an MCMC move with near-unity acceptance probability; the deviations from unity exactly correct for the approximate nature of the proposal. As with the proposal for the mutational history along a branch, this proposal using a Jukes-Cantor model and near-parsimonious intuition to make a good proposal, while relying on the Metropolis-Hastings criterion to obtain correct results in the general case.

Concretely, we adopt the following form for the grafting point proposal probability:

$$\alpha_{\text{graft}}(\mathsf{o} \to \mathsf{n}) \propto \exp[-\mu L(t_X - t_{P'})] \cdot [\mu/3]^{N(P')} \cdot (t_X - t_{P'})^{N(P')}. \tag{22}$$

The additional factors of $(t_X - t_{P'})^{N(P')}$ approximately cancel the corresponding factors in $\alpha_{\text{mut}}(\mathsf{o} \to \mathsf{n})$ that account for choosing concrete times for all the relevant mutations (cf., Eq. (20)). The above equation provides an unnormalized probability density for every possible regrafting point $P'$ in terms of easily calculable quantities (see Figure 20). In particular, the quantity $N(P')$ can be computed trivially in an EMAT through a depth-first traversal of the tree starting at $P$ (see annotations in Figure 19). It is this property of EMATs that greatly facilitates the calculation of $N(x)$ everywhere in the tree that motivates our naming this technique *mutation-directed* SPR moves: in effect, the explicit mutations on the tree serve as an excellent and efficient guide to choosing regrafting points well, even over large distances.

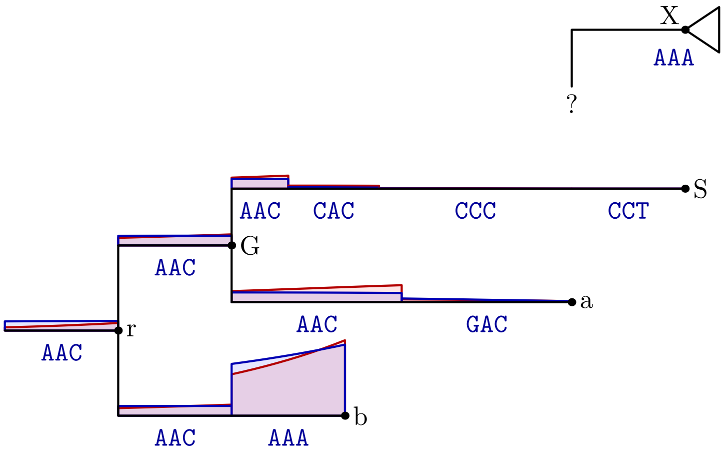


**Supplementary Figure 20:** Cartoon of grafting site probabilities calculated by mutation-directed SPR move (red) vs. distribution implied by the posterior (blue). In this example, the real posterior has an HKY substitution model with $\kappa = 5.25$, $\pi = [0.29, 0.18, 0.19, 0.33]$, which is representative of the SARS-CoV-2 data in [11]. Note that probabilities are broadly similar. The parsimonious region with sequence AAA, where attaching $X$ necessitates no mutations on the $P'$–$X$ branch, is clearly the most likely choice, but all other regions are possible, and their combined weight need not be negligible.

To be more concrete, we divide the tree left over after pruning $X$ into $K$ regions as follows. For branch $i$, let $t_1, \ldots, t_{|\mathcal{M}_i|}$ be the times of the mutations on it; let $t_0$ be the time of the start of the branch (or $-\infty$

if $i$ is the root) and let $t_{|\mathcal{M}_i|+1}$ be the time of the end of the branch. Then record the region of branch $i$ spanning the time $(t_j, t_{j+1})$ for each $j$ between 0 and $|\mathcal{M}_i|$ (inclusive). Finally, remove all regions that are entirely to the future of $t_X$; those regions that span $t_X$ are also truncated at $t_X$. All points $x$ in each region have the same value of $N(x)$, which we also record while constructing the regions. In the end, a region $k$ thus consists of a tuple $(i_k, t_k, t'_k, N_k)$, respectively the branch index, the region's start and end times, and the number of sequence differences between any point in the region and $X$. We call such a set of annotated regions an "SPR study". With this information, we define a weight $w_k$ for each region as follows:

$$w_k = \int_{t_k}^{t'_k} dt\, \exp[-\mu L(t_X - t)][\mu(t_X - t)/3]^{N_k}. \tag{23}$$

To pick a new regrafting point, pick a region $k$ in proportion to the weights $\{w_k\}$. Then pick a point between $t_k$ and $t_{k'}$ in proportion to

$$p(t|k) \propto \exp[-\mu L(t_X - t)][\mu(t_x - t)/3]^{N_k}. \tag{24}$$

All of the above calculations can be implemented in terms of incomplete Gamma functions and their inverses, which are widely available (e.g., in `boost::math` for C++ and in `scipy.special` in Python).

The above procedure describes the "pure" version of mdSPR. In practice, we apply the following changes that make the move more efficient to execute:

- When walking the tree to enumerate regions and calculate $N(x)$, we skip over mutations on sites $\ell \notin \xi(X)$, since they would not require additional mutations along the $P'$–$X$ branch owing to `N`-pruning.
- Particularly after the burn-in period of the MCMC, it is wasteful to consider possible new grafting points over the entire tree, since $X$ is presumably grafted near a locally optimal spot. Thus, most of the time, we limit the walk on the tree to regions for which $N_k \leq 1$; when the walk would cross a mutation that would make $N_k = 2$, we truncate the walk and begin to backtrack. Nevertheless, about 1 % of the time, we do perform complete explorations of the tree, to account for nodes $X$ which could be comfortably grafted in multiple, separated areas of the tree, and to always allow the MCMC some chance of escaping local minima.
- We observed that sometimes mdSPR would make good proposals too eagerly compared to what was implied by the actual evolution model, and which would then be rejected according to the Metropolis-Hastings criterion. For example, proposals near the beginning of a run where the $P$–$X$ branch had dozens of mutations and the $P'$–$X$ branch had 0 or 1 would be spuriously rejected! To mitigate this issue, we actually use a dampened p.d.f. for the grafting point:

  $$\alpha_{\text{graft}}(\mathsf{o} \to \mathsf{n}) \propto \left[\exp[-\mu L(t_X - t_{P'})] \cdot [\mu/3]^{N(P')} \cdot (t_X - t_{P'})^{N(P')}\right]^f.$$

  The annealing factor $f$, so-called because it plays an analogous role to a temperature factor in simulated annealing, leads to mdSPR making the most promising grafting proposals less eagerly than implied by a Jukes-Cantor model, which then leaves some room for the Metropolis-Hastings factor to make corrections without driving the acceptance probability to 0 (at the cost of occasionally making and rejecting grafting proposals that are less promising). Empirically, we found that $f = 0.8$ is sufficient to keep mdSPR effective while accepting most of the beneficial proposals in the burn-in phase.
- Finally, since incomplete Gamma functions and their inverses are expensive to calculate, we actually approximate the integrand in the definition of $w_K$ by its value at the midpoint time $t_* = (t'_k + t_k)/2$, so that

  $$w_k \approx (t'_k - t_k)\left[\exp[-\mu L(t_X - t_*)][\mu(t_X - t_*)/3]^{N_k}\right]^f.$$

  Correspondingly, in those regions, we approximate $p(t|k)$ by a uniform distribution between $t_k$ and $t'_k$. We have not made any attempt to detect regions for which this approximation would be inappropriate, such as long branches or branches that span the maximum of the integrand. However, the approximation cannot be applied for the region above the root (where $t_k = -\infty$), so there, we do use the full expression and pay the cost of using incomplete Gamma functions.

As noted in the main text, we have empirically observed that MCMC runs for genomic epidemiology datasets (real and simulated), which previously ran into convergence problems and/or for which we needed to hand-tune step sizes in the subtree slide move, all converge reliably when mdSPR moves are used, even though mdSPR exposes no tunable parameters. We thus believe that mdSPR plays a foundational role in making our MCMC moveset robust without requiring any parameter tuning.

##### 6.7 Other local moves

Besides SPR moves, we have implemented two other cheap move types to rapidly and efficiently sample a tree's mutational history and branch lengths while keeping its topology fixed:

1. *Inner node displacement moves*: We first pick a non-tip node $X$ and note its allowable range of times $(t_{\min}, t_{\max})$, subject to preserving the tree topology and the times of all mutations on the branches impinging on it. If $X$ is not the root, we pick a new time directly from the exponential distribution $\exp[+\lambda_X t]$ with $t_{\min} < t < t_{\max}$. This choice of $t_X$ exactly cancels the ratio of genetic priors in the acceptance probability, so only the ratio of coalescent priors must be calculated, which is usually close to 1. If, however, $X$ is the root node, we instead add to the old time a Gaussian-distributed displacement with mean 0 and standard deviation $\lambda_X^{-1}/2$ (as in the subtree slide move). For non-root nodes, this move efficiently approximates Gibbs sampling of branch lengths conditioned on the topology and mutational history not changing. In this sense, it is reminiscent of the Thorney Beast extension to BEAST, which has been used[21] for Bayesian inference of very large trees with à prior fixed topologies (up to polytomies).

2. *Branch reform moves*: We first pick a random branch $i$ other than the root branch. If $i$ is also not a child of the root, we pick new times for the mutations on branch $i$ uniformly between $t_{\varphi(i)}$ and $t_i$ (if there are multiple mutations on the same site, we sort the proposed times and preserve the relative order of the existing mutations). Because mutations only change $\lambda(x)$ very slightly over the length of the branch, this proposal is close to a Gibbs sampling of the mutation times, and so is almost always accepted. When $i$ is a child of the root, however, we instead execute an SPR move with $P' = P$, as described above; this allows mutations on one child branch of the root to easily migrate to the other branch, with a concomitant root sequence change.

Throughout the development of Delphy, we experimented with many other possibilities for local MCMC moves. These included: (a) "junction reform moves", which would resample the mutational history of all three branches impinging on a node simultaneously while keeping the rest of tree fixed; (b) narrow and wide exchange moves, analogous to those implemented in BEAST; and (c) "cluster" moves, wherein an SPR move proposes changing the mutational history of a connected cluster of branches, of which the $P'$–$X$ is but one, and amounts to a local ancestral state reconstruction. All of these were technically more complex to implement than the battery of moves described in this local MCMC moves section (particularly cluster moves), and yet we did not notice an appreciable effect on MCMC convergence from disabling them. In the interests of simplicity, robustness and ease of evolving Delphy in the future, we dropped all support for these other kinds of moves. In agreement with the experience of matOptimize, it seems that simple SPR moves, particularly when mutation-directed, suffice to explore tree space for the densely sampled datasets we are targeting.

#### 7 Global MCMC moves

We now focus on *global* MCMC moves, where we keep the topology and genetic history of the tree fixed but vary the associated parameters $\boldsymbol{\theta}$. As noted by Lartillot as early as 2006 [10], an explicit-mutation representation lends itself to surprisingly cheap global moves, often perfect Gibbs samplers. In contrast, when using an implicit representation, global moves involve a full recalculation of the tree likelihood, and so are among the most expensive moves in the MCMC. Here, we summarize the global moves used by Delphy, which essentially recap Lartillot's earlier proposals.

#### 7.1 Mutation rate $\mu$

The conditional posterior distribution for $\mu$ follows from Equation (1) up to normalization:

$$P_\mu(\mu) \propto e^{-\mu L\tilde{T}} \cdot \mu^M \cdot \pi_\mu(\mu). \tag{25}$$

Here, $M$ is the total number of mutations and $\tilde{T}$ is an effective total branch length of the tree, defined as

$$\tilde{T} = \frac{\int_{x\in\mathcal{T}} \lambda(x)\, dx}{\mu L}.$$

The right-hand side is independent of $\mu$, as follows from expanding the definitions of $\lambda(x)$ and $Q^{(\ell)}_{ab}$. We note that maintaining an updating the values of $M$ and $\tilde{T}$ is straightforward, and both can be calculated efficiently from scratch in time linear with the size of the tree.

An MCMC move that proposes a change from $\mu$ to $\mu'$ with probability $\alpha(\mu \to \mu')$ has acceptance probability

$$P_{\text{acc}}(\mu \to \mu') = \min\left[1, e^{-(\mu'-\mu)L\tilde{T}} \cdot \left(\frac{\mu'}{\mu}\right)^M \cdot \frac{\pi_\mu(\mu')}{\pi_\mu(\mu)} \cdot \frac{\alpha(\mu' \to \mu)}{\alpha(\mu \to \mu')}\right].$$

For a general prior $\pi_\mu(\mu)$, this acceptance probability can be applied to moves that perturb $\mu$, e.g., scaling moves. Because such a move is very cheap, it's beneficial to attempt many such moves in quick succession without changing the tree topology, in effect approximating a Gibbs sampler for $\mu$.

For typical forms of $\pi_\mu(\mu)$, namely an improper uniform prior and a $1/x$ prior, the conditional posterior in Equation (25) is a Gamma distribution, so we can go further and directly and cheaply Gibbs sample $\mu$:

$$\mu \sim \begin{cases} \text{Gamma}(M-1, L\tilde{T}), & \text{for } \pi_\mu(\mu) = 1; \\ \text{Gamma}(M, L\tilde{T}), & \text{for } \pi_\mu(\mu) = 1/\mu. \end{cases}$$

The above procedure also makes it clear that the conditional average of $\mu$ is close to $M/(L\tilde{T})$ (exactly so for a $1/x$ prior), a physically pleasing result.

In our initial implementation of Delphy, we have not implemented a move analogous to BEAST's up-down operator that can simultaneously scale the mutation rate and all the branch lengths while keeping their product roughly unchanged. While doing so would speed convergence for trees where tips are concentrated in a small time window at the end of the tree[3], we are targeting use cases where tips occur throughout a large fraction of the range of the tree, where such a simultaneous scaling move would have low acceptance probability.

#### 7.2 Evolution model parameters $\{\pi_a\}$, $\{q_a\}$ and $\{q_{ab}\}$

As for the mutation rate $\mu$, we can construct a joint conditional posterior distribution for the variables that specify the evolution model parameters, $\{\pi_a\}$, $\{q_a\}$ and $\{q_{ab}\}$. For the HKY model we use, while $\{\pi_a\}$ is specified directly, the other parameters are derived from the variable $\kappa$; the procedure below, however, is easily adapted to any other model of evolution. Thus, the joint conditional posterior distribution for $\kappa$ and $\{\pi_a\}$ that follows from Equation (1) is, up to normalization:

$$P_Q(\kappa, \{\pi_a\}) \propto \prod_a \pi_a^{R_a} \cdot e^{-\mu L \sum_a q_a \tilde{T}_a} \cdot \prod_{a,b} (q_{ab})^{M_{ab}} \cdot \pi_Q(\kappa, \{\pi_a\}). \tag{26}$$

Here, $R_a$ is the number of sites with state $a$ in the root sequence, $M_{ab}$ is the number of $a$-to-$b$ mutations in the tree, and $\tilde{T}_a$ is an effective total branch length in state $a$, defined as

$$\tilde{T}_a = \frac{1}{q_a \mu L} \int_{x\in\mathcal{T}} \sum_{\ell\in\xi(x)} [s^{(\ell)}(x) = a] Q^{(\ell)}_a.$$

As above, $\tilde{T}_a$ is actually independent of the values of $\mu$ and $q_a$. The above expression is valid when the same substitution model applies at all sites, modulated only by the site rates $\nu^{(\ell)}$, but is easy to generalize to multiple partitions. As noted before, these values can be calculated efficiently from scratch and maintained up to date after local MCMC moves.

As with the mutation rate, the above conditional posterior suggests very cheap moves for making small trial changes to $\kappa$ and $\{\pi_a\}$, for example, by scaling and delta-exchange, respectively. A long sequence of such moves approximates a Gibbs sampler for $\kappa$ and $\{\pi_a\}$, and can be implemented efficiently if there are no intervening topological or mutational changes to the tree.

Unlike with the mutation rate, the above conditional posterior does not immediately suggest a direct Gibbs sampler for any common choice of prior $\pi_Q(\kappa, \{\pi_a\})$. However, Nicola de Maio has pointed out to us that an UNREST evolution model[25] may be suitable for densely sampled datasets. In this model, each transition rate is inferred independently under a Gamma-distributed prior, and so the conditional posterior for all rates is a product of independent Gamma distributions; similarly, the parameters $\pi_a^{(\ell)}$ stop playing the role of the stationary distribution of the evolution model, and instead become free parameters with Dirichlet priors, so that their conditional posterior is a Dirichlet-multinomial distribution. These changes imply efficient Gibbs samplers for all the rates $Q_{ab}^{(\ell)}$ and $\pi_a^{(\ell)}$ [10]. We leave the exploration of this possibility for future work.

#### 7.3 Population model $N(t)$

The conditional posterior distribution for the parameters $n_0$ and $g$ that specify the population curve $N(t) = n_0 e^{gt}$ depends only on the coalescent prior and parameter priors, as follows:

$$P_N(n_0, g) \propto \pi_{\text{anc}}(\mathcal{T}|\boldsymbol{\theta})\pi_N(n_0, g).$$

The coalescent prior depends on the tree topology only through (a) the function $k(t)$ that counts the number of active branches at time $t$; and (b) the times $\{t_i\}$ of all the inner nodes. By using a fine-grained piecewise-constant approximation to $k(t)$, as described in Section 9, we can efficiently evaluate ratios of the above conditional posterior for old and new values of $n_0$ and $g$. Hence, we can execute a long sequence of consecutive MCMC moves that perturb $n_0$ and $g$ so as to approximate a Gibbs sampler for both.

#### 7.4 Site-rate heterogeneity parameters, $\alpha$ and $\{\nu^{(\ell)}\}$

In the standard approach to modeling site-rate heterogeneity, the Gamma distribution for the site relative rates $\{\nu^{(\ell)}\}$ is approximated by a discrete distribution with $K$ categories because it is not possible to analytically integrate the tree likelihood for all choices of $\{\nu^{(\ell)}\}$. Instead, the integration is approximated by $K$ evaluations of the tree likelihood at evenly distributed quantiles of $\nu^{(\ell)}$, with a concomitant $K$-fold increase in cost to every MCMC move that changes the tree likelihood. In contrast, in our explicit approach, we do not attempt to integrate these degrees of freedom analytically, but instead let the integration occur implicitly via sampling. In other words, both $\alpha$ and $\{\nu^{(\ell)}\}$ are explicit, continuous parameters of the inference, and individual MCMC moves are not appreciably more expensive when including vs excluding site-rate heterogeneity effects. We note that recent versions of MAPLE have also implemented an equivalent idea in the context of maximum likelihood phylogenetics [13].

It is tempting to evaluate conditional posterior distributions for $\alpha$ and $\{\nu^{(\ell)}\}$ separately, and then implement separate moves on each. The relevant joint conditional posterior distribution again follows from Equation (1) up to normalization:

$$P_\nu(\alpha, \{\nu^{(\ell)}\}) \propto \pi_\alpha(\alpha) \cdot \prod_\ell e^{-\mu\nu^{(\ell)}\tilde{T}^{(\ell)}} \cdot \left[\nu^{(\ell)}\right]^{M^{(\ell)}} \cdot \frac{\alpha^\alpha}{\Gamma(\alpha)} \left[\nu^{(\ell)}\right]^{\alpha-1} e^{-\alpha\nu^{(\ell)}}. \tag{27}$$

Hence, the individual conditional posterior distributions are:

$$P_\alpha(\alpha) \propto \pi_\alpha(\alpha) \cdot \prod_\ell \frac{\alpha^\alpha}{\Gamma(\alpha)} \left[\nu^{(\ell)}\right]^{\alpha-1} e^{-\alpha\nu^{(\ell)}},$$

$$P_{\nu^{(\ell)}}(\nu^{(\ell)}) \propto e^{-\mu\nu^{(\ell)}\tilde{T}^{(\ell)}} \cdot \left[\nu^{(\ell)}\right]^{M^{(\ell)}} \cdot \left[\nu^{(\ell)}\right]^{\alpha-1} e^{-\alpha\nu^{(\ell)}}.$$

Here, $M^{(\ell)}$ is the number of mutations in site $\ell$ and $\tilde{T}^{(\ell)}$ is an effective total branch length for site $\ell$, given by

$$\tilde{T}^{(\ell)} := \frac{1}{\mu} \int_{x \in \mathcal{T}} [\ell \in \xi(x)] Q_a^{(\ell)}\, dx.$$

It is impractical to update the vector $\tilde{T}^{(\ell)}$ after every local MCMC move. However, this vector can be efficiently calculated in time proportional to the sum of the tree size and the number of mutations and missations as follows: (a) accumulate the total branch length of every subtree rooted at inner node $i$, as if there were no missing data, by a post-order traversal of the tree; (b) calculate an initial value of $\tilde{T}^{(\ell)}$ for each site using the reference sequence and the total branch length at the root node, as if there were no mutations or missations on the tree; then (c) walk the tree and adjust $\tilde{T}^{(\ell)}$ for each observed mutation and missation at site $\ell$.

Observe that, like the prior $\pi_\nu(\nu^{(\ell)})$, the conditional posterior for $\nu^{(\ell)}$ given above is the distribution Gamma$(\alpha + M^{(\ell)}, \alpha + \mu\tilde{T}^{(\ell)})$. The tree topology and mutational history of site $\ell$ simply change the prior's Gamma parameters in a way that compares the number of mutations on this site expected in the absence of site-rate heterogeneity ($\mu\tilde{T}^{(\ell)}$) with the actual number ($M^{(\ell)}$), and $\alpha$ modulates the extent to which discrepancies are ascribed to real differences in rates vs stochastic fluctuations. Moreover, because the posterior distribution for $\nu^{(\ell)}$ is a Gamma distribution, we can draw new conditional samples of $\{\nu^{(\ell)}\}$ efficiently. For $\alpha$, though, we are again reduced to making trial changes, e.g., by scaling, and accepting or rejecting them in the usual way.

In practice, the above scheme of separately sampling $\alpha$ and $\{\nu^{(\ell)}\}$ is not effective. The values of $\alpha$ and $\{\nu^{(\ell)}\}$ are very correlated, and independent moves along each of them do not effectively explore the joint parameter space. However, one can marginalize the $\{\nu^{(\ell)}\}$ in Equation (27) to get an effective conditional posterior distribution for $\alpha$:

$$\tilde{P}_\alpha(\alpha) \propto \pi_\alpha(\alpha) \cdot \prod_\ell \frac{\Gamma(\alpha + M^{(\ell)})}{\Gamma(\alpha)} \frac{\alpha^\alpha}{(\alpha + \mu\tilde{T}^{(\ell)})^{\alpha + M^{(\ell)}}}$$

Although the function $\tilde{P}_\alpha(\alpha)$ is expensive to calculate, it is relatively smooth, so repeated trial changes to $\alpha$ will have high acceptance probabilities and quickly approximate a Gibbs sampler of $\tilde{P}_\alpha(\alpha)$. After thus obtaining a new value of $\alpha$, we directly sample all $L$ new values of $\nu^{(\ell)}$ as described above.

### 8 Scaling strategy

In matOptimize, the search for a parsimonious tree is parallelized by observing that the effect of an SPR moves is quite localized, so two SPR moves in different portions of the tree can first be evaluated separately, and their combined effect deduced from those separate evaluations. In other words, the net effect of two non-conflicting SPR moves is independent of the order in which those moves are performed. The MCMC SPR moves on EMATs described above have an analogous property: if we partition a tree into non-overlapping subtrees and independently perform SPR moves in each subtree in parallel, the net result is equivalent to performing those moves serially by interleaving the subtree moves in any way. This idea is depicted schematically in Figure 21.

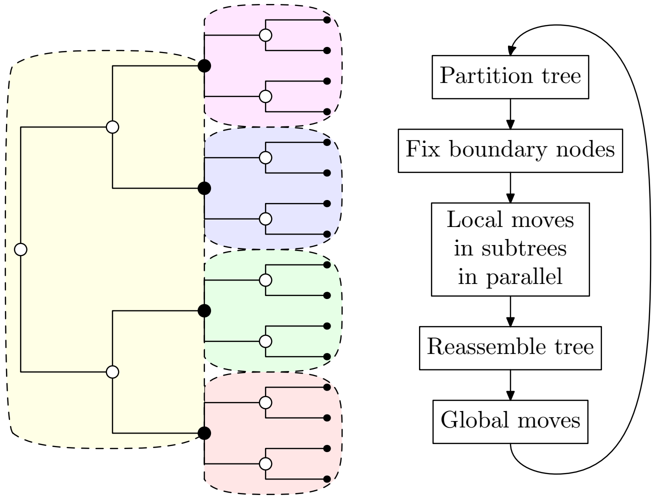


**Supplementary Figure 21:** Schematic of Delphy's scaling strategy. In each cycle, the sequence and times of the boundary nodes (filled) is fixed, while the sequence, times and topology of the remaining nodes (unfilled) can change. Within each shaded subtree, local MCMC moves are confined to that subtree. To ensure ergodicity, the fixed boundary nodes on successive cycles do not overlap.

### 9 Parallelizable coalescent prior

As is common in Bayesian phylogenetics of viruses, our ancestry prior $\pi_\varphi(\mathcal{T}|\boldsymbol{\theta})$ is the Kingman coalescent [7, 8], given by

$$\pi_{\mathrm{anc}}(\mathcal{T}|\boldsymbol{\theta}) := \exp\left[-\int \binom{k(t)}{2}\frac{dt}{N(t)}\right]\cdot \prod_{\text{inner nodes } j}\frac{1}{N(t_j)}, \tag{28}$$

where $k(t)$ is the number of active branches at time $t$ and $N(t)$ is the product of the effective population size and the generation time.

A significant complication in parallelizing the coalescent prior is that it directly couples all the nodes at the end of branches that cross a given time $t$. Hence, at first glance, the use of a coalescent prior seems in conflict with the scaling strategy described above. In this section, we describe in detail one scheme that resolves that conflict through the introduction of an augmented coalescent prior. We emphasize that this scheme is not the only possible scheme, nor is it necessarily the best one. Better schemes may yet improve the sampling.

First, instead of evaluating Eq (28) piecewise directly, we start by discretizing it into finite time segments "cells" of width $\Delta$ over the range of times spanned by the tree:

$$\pi_\varphi(\mathcal{T}|\boldsymbol{\theta}) \approx \bar{\pi}_\varphi(\mathcal{T}|\boldsymbol{\theta}) := \exp\left[-\sum_c \binom{\bar{k}_c}{2}\frac{\Delta}{\bar{N}_c}\right]\cdot \prod_{\text{inner nodes } j}\frac{1}{N(t_j)}. \tag{29}$$

Here and below, a barred variable, say $\bar{k}_c$, stands for the mean value of a corresponding function of time, say $k(t)$, over the time interval spanned by cell $c$. In our current implementation, we partition the range over which $k(t) > 0$ into roughly 400 intervals, periodically changing $\Delta$ if the range of $k(t)$ changes drastically. In practice, $\Delta$ is readjusted a handful of times during the burn-in period as the tree height converges, but remains stable throughout the production period of the MCMC run. Note that this discretization already results in substantial efficiency improvements, because the cost to displacing an inner node scales with the change in time, not with the density of coalescence events in other parts of the tree around the node's time.

Because the coalescent prior introduces only modest gradients into the posterior, the crudeness of the above approximation does not seem to impact results appreciably. One notable exception is in very large datasets, such as the simulated 100 000-sample outbreak, where the $k(t)^2$ dependence of the integrand and the typically high values of $k(t)$ require a much higher number of cells to mitigate the discretization errors introduced by the above approximation. It remains to be seen whether more sophisticated approximations to the integral in Eq. (28) are also amenable to the parallelization scheme described below.

Second, we split the tree $\mathcal{T}$ into $P$ partitions. It follows that we can decompose the number of active lineages $\bar{k}_c$ into a sum over the numbers of active lineages $\bar{k}_{p,c}$ in partition $p$, $\bar{k}_c = \sum_p \bar{k}_{p,c}$. Rewriting Eq. (29) in terms of partitions yields

$$\bar{\pi}_\varphi(\mathcal{T}|\boldsymbol{\theta}) = \prod_p \left\{ \prod_c \exp\left[ -\Delta \left( \frac{\bar{k}_{p,c} \cdot [\sum_q \bar{k}_{q,c}]}{2\bar{N}_c} - \frac{\bar{k}_{p,c}}{2\bar{N}_c} \right) \right] \cdot \prod_{j \in p} \frac{1}{N(t_j)} \right\}. \tag{30}$$

This form emphasizes the coupling between partitions $p$ and $q$ owing to the quadratic term in the exponential.

An analogy with electrostatics can be drawn at this point: the potential energy of a set of particles can be calculated either as the sum of pairwise direct Coulomb interactions between all pairs of particles, or by a sum of local interactions between charges and an induced local potential. In other words, we can replace direct interaction at a distance between charges with a local interaction mediated by a potential that transmits information through space. This picture suggests introducing for each cell $c$ a set of $P$ augmenting Gaussian variables $\tilde{k}_{p,c}$ that couple linearly to the $\{\bar{k}_{p,c}\}$ according to the following *augmented coalescent prior*:

$$\tilde{\pi}_\varphi(\mathcal{T}|\boldsymbol{\theta}, \{\tilde{k}_{p,c}\}) := \prod_p \left\{ \prod_c \mathcal{C}_{p,c} \exp\left[ -\Delta \left( \frac{(\tilde{k}_{p,c} - \mu_{p,c})^2}{2\sigma_{p,c}^2} + \frac{\tilde{k}_c \cdot \bar{k}_{p,c}}{\bar{N}_c} - \frac{\bar{k}_{p,c}}{2\bar{N}_c} \right) \right] \cdot \prod_{j \in p} \frac{1}{2N(t_j)} \right\}. \tag{31}$$

Above, we've introduced the sum $\tilde{k}_c$ analogous to $\bar{k}_c$, defined as $\tilde{k}_c := \sum_p \tilde{k}_{p,c}$. The mean $\mu_{p,c}$ and variance $\sigma_{p,c}^2$ of the free variable $\tilde{k}_{p,c}$, as well as the normalization constants $\mathcal{C}_{p,c}$ are not yet specified. We make the following choices:

$$\mu_{p,c} = \bar{k}_{p,c}, \qquad \sum_p \sigma_{p,c}^2 = \bar{N}_c, \qquad \mathcal{C}_{p,c} = \left(2\pi\sigma_{p,c}^2/\Delta\right)^{-1/2}. \tag{32}$$

The second equation does not fully specify $\sigma_{p,c}$; for concreteness, we choose $\sigma_{p,c}^2 = \bar{N}_c/P_c$, where $P_c$ is the number of partitions with active branches in cell $c$, i.e., where $k_{p,c} > 0$. We claim that the above choices lead to the augmented coalescent prior reducing to Eq. (30) once the augmenting variables $\tilde{k}_{p,c}$ are integrated out, that is,

$$\int \left\{ \prod_{p,c} d\tilde{k}_{p,c} \right\} \tilde{\pi}_\varphi(\mathcal{T}|\boldsymbol{\theta}, \{\tilde{k}_{p,c}\}) = \bar{\pi}_\varphi(\mathcal{T}|\boldsymbol{\theta}). \tag{33}$$

The calculation is tedious but straightforward:

$$\int \left\{ \prod_{p,c} d\tilde{k}_{p,c} \right\} \tilde{\pi}_\varphi(\mathcal{T}|\boldsymbol{\theta}, \{\tilde{k}_{p,c}\}) \tag{34}$$

$$= \prod_{p,c} \int d\tilde{k}_{p,c}\, \mathcal{C}_{p,c} \exp\left[ -\Delta \left( \frac{(\tilde{k}_{p,c} - \mu_{p,c})^2}{2\sigma^2_{p,c}} + \frac{\tilde{k}_{p,c} \cdot \bar{k}_c}{\bar{N}_c} - \frac{\bar{k}_{p,c}}{2\bar{N}_c} \right) \right] \cdot \prod_j \frac{1}{2N(t_j)}, \tag{35}$$

$$= \prod_{p,c} \int d\tilde{k}_{p,c}\, \mathcal{C}_{p,c} \exp\left[ -\Delta \left( \frac{\tilde{k}^2_{p,c}}{2\sigma^2_{p,c}} + \frac{(\tilde{k}_{p,c} + \mu_{p,c}) \cdot \bar{k}_c}{\bar{N}_c} - \frac{\bar{k}_{p,c}}{2\bar{N}_c} \right) \right] \cdot \prod_j \frac{1}{2N(t_j)}, \tag{36}$$

$$= \prod_{p,c} \exp\left[ \frac{(\Delta\bar{k}_c/\bar{N}_c)^2}{2\Delta/\sigma^2_{p,c}} \right] \cdot \exp\left[ -\Delta \left( \frac{\mu_{p,c} \cdot \bar{k}_c}{\bar{N}_c} - \frac{\bar{k}_{p,c}}{2\bar{N}_c} \right) \right] \cdot \prod_j \frac{1}{2N(t_j)}, \tag{37}$$

$$= \prod_{p,c} \exp\left[ -\Delta \left( -\frac{\bar{k}^2_c \sigma^2_{p,c}}{2\bar{N}^2_c} + \frac{\mu_{p,c} \cdot \bar{k}_c}{\bar{N}_c} - \frac{\bar{k}_{p,c}}{2\bar{N}_c} \right) \right] \cdot \prod_j \frac{1}{2N(t_j)}, \tag{38}$$

$$= \prod_{c} \exp\left[ -\Delta \left( -\frac{\bar{k}^2_c}{2\bar{N}_c} + \frac{\bar{k}_c \cdot \bar{k}_c}{\bar{N}_c} - \frac{\bar{k}_c}{2\bar{N}_c} \right) \right] \cdot \prod_j \frac{1}{2N(t_j)}, \tag{39}$$

$$= \exp\left[ -\sum_c \binom{\bar{k}_c}{2} \frac{\Delta}{\bar{N}_c} \right] \cdot \prod_j \frac{1}{2N(t_j)}, \tag{40}$$

$$= \bar{\pi}_\varphi(\mathcal{T}|\boldsymbol{\theta}). \tag{41}$$

The first equation follows from the relation $\sum_p \tilde{k}_c \cdot \bar{k}_{p,c} = \sum_{p,p'} \tilde{k}_{p,c} \cdot \bar{k}_{p',c} = \sum_p \tilde{k}_{p,c} \cdot \bar{k}_c$. The second equation follows from a change of variables $\tilde{k}'_{p,c} = \tilde{k}_{p,c} - \mu_{p,c}$. The third equation follows from the well-known Gaussian integral $\int dx\, \exp[-(ax^2 + bx + c)] = \sqrt{\pi/a} \exp[b^2/4a - c]$. The fourth equation simply regroups terms, while the fifth simultaneously substitutes $\mu_{p,c}$ and $\sigma^2_{p,c}$ with the values in Eq. (32), turns the product over $p$ of exponentials into an exponential of sums over $p$, and evaluates them. The equality with $\bar{\pi}_\varphi(\mathcal{T}|\boldsymbol{\theta})$ then follows straightforwardly.

The above transformation paves the way for parallelizing the coalescent prior. First, notice that for fixed $\{\bar{k}_{p,c}\}$, the variables $\{\tilde{k}_{p,c}\}$ can be Gibbs-sampled, because they are independent Gaussians. Concretely, the dependence of $\tilde{\pi}_\varphi$ on a particular $\tilde{k}_{p,c}$ conditional on all other variables being fixed is:

$$\tilde{\pi}_\varphi(\mathcal{T}|\boldsymbol{\theta}, \{\tilde{k}_{p,c}\}) \sim \exp\left[ -\Delta \left( \frac{(\tilde{k}_{p,c} - \bar{k}_{p,c})^2}{2\sigma^2_{p,c}} + \frac{\tilde{k}_{p,c} \cdot \bar{k}_c}{\bar{N}_c} \right) \right], \tag{42}$$

$$\sim \exp\left[ -\frac{\left( \tilde{k}_{p,c} - \left\{ \bar{k}_{p,c} - \frac{\bar{k}_c \sigma^2_{p,c}}{\bar{N}_c} \right\} \right)^2}{2\sigma^2_{p,c}/\Delta} \right], \tag{43}$$

$$\tag{44}$$

and so

$$\tilde{k}_{p,c} \sim \mathcal{N}\left( \bar{k}_{p,c} - \frac{\bar{k}_c \sigma^2_{p,c}}{\bar{N}_c}, \sigma^2_{p,c}/\Delta \right), \qquad \text{for fixed } \{\bar{k}_{p,c}\}. \tag{45}$$

Second, for fixed $\{\tilde{k}_{p,c}\}$, Eq. (31) is a product of independent terms, one for each partition $p$. Hence, we can make independent local moves in each partition $p$ with only knowledge of $\tilde{k}_{p,c}$, $\bar{k}_c$ and $\bar{N}_c$.

The above considerations lead to the following parallelization scheme:

1. Cut up the tree $\mathcal{T}$ into $P$ partitions.

2. For every cell $c$, calculate $\bar{k}_{p,c}$ for every partition $p$ and then calculate their sum $\bar{k}_c$.
3. For every cell $c$, sample random Gaussian fields $\tilde{k}_{p,c}$ for every partition $p$ as per Eq. (45), then calculate their sum $\tilde{k}_c$.
4. Distribute the $P$ partitions over $P$ parallel workers. Worker $p$ receives the subtree in partition $p$, and the full set of values for $\tilde{k}_{p,c}$ and $\tilde{k}_c$.
5. Apply local MCMC moves within each worker using the augmented coalescent prior of Equation (31). By the above arguments, each worker has enough information to do this correctly and independently.
6. Reassemble the modified subtrees from each of the workers into a global tree.
7. Apply global MCMC moves, such as changes in mutation rates or evolution model parameters. This is done with the discretized coalescent prior given by Eq. (29).

Intuitively, at the beginning of each cycle, a central process has a global view of the tree. It takes a fuzzy "snapshot" of this global view, in the form of $\{\tilde{k}_{p,c}\}$ and $\tilde{k}_c$. Then, each worker adjusts the various $\{\bar{k}_{p,c}\}$ in accordance with that earlier snapshot, not in accordance to the instantaneous $\bar{k}_c$, which they cannot access. Periodically, the tree is reassembled to be repartitioned and resnapshotted. Statistically, this process ends up transmitting the global knowledge needed to impose the coalescent prior to every worker.

A subtle disadvantage of the partitioning scheme presented in this section is that the partitioning enters the posterior distribution indirectly, so that factors arising from the partitioning in principle enter the acceptance probability of all MCMC moves. We describe the problem in more detail below, and present our strategy for mitigating it almost completely.

#### 9.1 Partitioning a tree into $P$ partitions

The key mechanism for parallelization in Delphy is to partition the tree $\mathcal{T}$ into roughly $P$ partitions, and then perform local moves in all partitions in parallel. The basic scheme is inspired by JUNIPER [22], and is itself based on the work of Borndörfer, Elijazyfer and Schwartz [1]. It identifies a series of nodes, which we call *cut points*, where one subtree ends and another begins. Each subtree $\mathcal{T}_p$ is rooted at either the root of $\mathcal{T}$ or one of the cut points, and extends as far as the closest descendant cut points or the tips of the original tree, whichever is closets. A simple heuristic tries to make the subtrees of roughly equal size, and we add an element of randomness to ensure that any particular point on the tree has a decent chance of being inside a subtree instead of at a boundary.

In concrete terms, we choose the cut points with the following algorithm. We perform a post-order traversal of the tree, descending from a node to its children in random order. In the process, we keep track of the number of nodes $m$ yet to be visited (initially $2N-1$) and the number of partitions $p$ yet to be formed (initially $P$). Throughout, we also construct the number of descendants $n_i$ of each visited node $i$; if $i$ is a tip, then $n_i = 1$, otherwise $n_i = 1 + \sum_j n_j$, where the sum runs over the children of $i$. If at any point $n_i$ is at least $\lambda = \max(10, m/(p+1))$, then node $i$ is considered a viable cut point. If the number of nodes remaining to visit is also at least $\lambda$, and a random number $u \sim U(0,1)$ satisfies $u < 0.5$ (i.e., a fair coin toss), then node $i$ becomes an actual cut point. We record it, then set $n_i = 0$ before continuing the walk. Finally, we always record the root node as a cut point. The various thresholds and randomizations ensure that no one partition is too large or too small; that if an overlay small or large partition is formed by chance, then the target size for the remaining partitions is adjusted accordingly; and that different executions of the algorithm can yield different partitionings of the same tree.

As long as partitionings happen regularly and they only enter in the proposal probabilities, such as by limiting the possible regrafting points $P'$ of an SPR proposal to the same partition that contains the pruned subtree, the exact partitioning details do not generally affect the results of the MCMC. However, as mentioned above, our scheme for the coalescent prior goes beyond that: the details of the partitioning appear directly in the posterior distribution. To be concrete, suppose that the probability of the above algorithm producing a partition $\Upsilon$ given a tree $\mathcal{T}$ and parameters $\boldsymbol{\theta}$ is $P(\Upsilon|\mathcal{T}, \boldsymbol{\theta})$. We can conceive of the parallelized

MCMC as sampling from an augmented state space consisting of trees $\mathcal{T}$, parameters $\boldsymbol{\theta}$, partitions $\Upsilon$ and the partition-dependent variables $\{\tilde{k}_{p,c}\}$. The posterior distribution sampled by this MCMC is actually

$$P_{\|}(\mathcal{T}, \boldsymbol{\theta}, \Upsilon, \{\tilde{k}_{p,c}\}) \propto P(\{\tilde{k}_{p,c}\}|\Upsilon, \mathcal{T}) P(\Upsilon|\mathcal{T}) P(\mathcal{T}, \boldsymbol{\theta}).$$

Here, the last factor is the posterior distribution that we actually want to sample (Eq. (1)), while

$$P(\{\tilde{k}_{p,c}\}|\Upsilon, \mathcal{T}) = \frac{\tilde{\pi}_{\varphi}(\mathcal{T}|\boldsymbol{\theta}, \{\tilde{k}_{p,c}\})}{\bar{\pi}_{\varphi}(\mathcal{T}|\boldsymbol{\theta})}.$$

As detailed in the previous section, $P(\{\tilde{k}_{p,c}\}|\Upsilon, \mathcal{T})$ is such that the entire posterior can be decomposed into independent factors for each partition, while its integral over all possible values of $\{\tilde{k}_{p,c}\}$ is exactly 1 (Eq. (33)).

The key flaw in $P_{\|}(\mathcal{T}, \boldsymbol{\theta}, \Upsilon, \{\tilde{k}_{p,c}\})$ is that, conditioned on $\{\tilde{k}_{p,c}\}$ being fixed, the factor $P(\Upsilon|\mathcal{T})$ depends on $\mathcal{T}$, so moves that propose a change in the topology of a subtree, say from $\mathcal{T}_{\mathsf{o}}$ to $\mathcal{T}_{\mathsf{n}}$, require a factor of $P(\Upsilon|\mathcal{T}_{\mathsf{n}})/P(\Upsilon|\mathcal{T}_{\mathsf{o}})$, whose neglect leads to incorrect sampling. We have verified empirically in a simple toy setting of a 2-tip tree whose MRCA is restricted to one of a finite number of positions, and a simplified partitioning scheme that admits an exact expression for $P(\Upsilon|\mathcal{T})$, that: (a) failing to account for $P(\Upsilon|\mathcal{T})$ does lead to incorrect sampling; and (b) adding the missing ratios of this probability to the Metropolis-Hastings criterion restores correct sampling.

One correct but inefficient solution to the above problem is to pick a partitioning scheme where $P(\Upsilon|\mathcal{T})$ is a constant, so that the "missing" correction factor in the Metropolis-Hastings criterion is just 1. One such scheme would be to choose $P$ cutpoints uniformly at random from the $N-1$ inner nodes of the tree, subject to one of these being the root. However, most inner nodes are close to a tip; for example, in the limit where all the tips occur at the same time and the tree is a complete binary tree, then around half of the inner nodes are the parent of two tips. Hence, this simple scheme tends to produce $P-1$ partitions with a handful of nodes, and a single enormous partition with the remainder of the tree, which renders parallelization utterly ineffective.

Given the above considerations, we instead introduce the following controlled approximation. Periodically, we choose a set of $D$ fixed partitionings (currently, 10), which we call "stencils". Throughout the MCMC, we sample partitionings uniformly among these $D$ stencils, so that Metropolis-Hastings correction owing to $P(\Upsilon|\mathcal{T})$ is again 1. After sufficiently many MCMC steps, the samples drawn from $P_{\|}$, when projected to remove $\Upsilon$ and $\{\tilde{k}_{p,c}\}$, are indistinguishable from those drawn from $P(\mathcal{T}|\boldsymbol{\theta})$ (indeed, this is the essential property of the augmented coalescent prior). At that point, it we pick a new set of $D$ stencils, and start this cycle again.

In detail, the final approximate scheme we have implemented in Delphy is as follows:

1. Use any algorithm, such as the one at the beginning of this section, to produce $D$ distinct stencils. Each stencil consists of the node identifiers of all the cutpoints, so any topology of $\mathcal{T}$ can be partitioned according to a particular stencil.

2. Proceed with the MCMC as described at the end of Section 9, whereby "cut up the tree $\mathcal{T}$ into $P$ partitions" means "sample one of the $D$ stencils uniformly and use it to partition $\mathcal{T}$".

3. After many rounds of the previous step (currently, 100), go back to the Step 1 to produce a fresh new set of $D$ stencils.

We have verified in our toy 2-tip system that such uniform cycling over fixed stencils, with infrequent periodic redrawing of the stencils, recovers correct MCMC sampling, even in the face of complex distributions for $P(\Upsilon|\mathcal{T})$. The excellent agreement of Delphy's results with the BEAST benchmark results also suggests that the scheme works in the general setting of complex trees. We note that as one of the above cycles progresses, and inner nodes in the tree move, the stencils used to partition the tree become increasingly ill-fitting, and would eventually degrade to the random-cut-points scheme of the previous paragraph if they are not refreshed. Refreshing every 100 iterations of Step 2 seems to be a reasonable compromise between complete correctness and efficiency.

### 10 Overall schedule of MCMC moves

Having described all local and global moves, as well as our parallelization scheme, we briefly sketch out how these are assembled in an overall move schedule. Most of these choices are arbitrary; many alternative choices would also be correct, but may impact the efficiency of sampling. Our concrete choices below are inspired by the BEAST run in [11]:

- Inside an outer loop, repeatedly create $D = 10$ partition stencils, then run the following steps 100 times (see previous section):
  - Inside a middle loop: (1) run a fixed schedule of global moves; (2) partition the tree into subtrees according to one of the $D$ stencils chosen at random; (3) run a fixed number of independent local moves in each subtree in parallel; (4) reassemble the tree from the modified subtrees. Concretely:
    - Run the following global moves:
      - A Gibbs sample of $\mu$, as described in Section 7.1.
      - A series of 10 delta-exchange moves to the HKY stationary base frequencies $\{\pi_a\}$ interleaved with 10 scaling moves of the HKY $\kappa$ parameter, as described in Section 7.2. This approximates a Gibbs sample of the evolution model parameters.
      - When site-rate heterogeneity is enabled: a series of 10 scaling moves of the $\alpha$ parameter after marginalizing the individual relative rates $\{\nu^{(\ell)}\}$, followed by Gibbs sampling of these rates, as described in Section 7.4. This approximate a Gibbs sample of the site-rate heterogeneity parameters.
      - A series of 50 scaling moves on the population size $n_0$ interleaved with 50 uniform delta moves on the population growth rate $g$. This approximate a Gibbs sample of the population model parameters.
    - We partition the tree into around $P$ subtrees using one of the $D$ stencils. We then calculate the number of active branches $\{k_{p,c}\}$ in each partition and cell, and use them to Gibbs sample values for $\{\tilde{k}_{p,c}\}$ of the augmented coalescent prior, as described in Section 9, as well as their sums $\{\tilde{k}_p\}$.
    - We then dispatch an inner loop to as many independent workers as there are partitions. Worker $p$ receives a copy of subtree $p$, the values of $\{\tilde{k}_{p,c}$ for its $p$, as well as the sum $\tilde{k}_c$, and copies of all the global model parameters: $\mu$, $\{\nu^{(\ell)}\}$, $\{\pi_a\}$, $\{q_a\}$, $\{q_{ab}\}$, $n_0$ and $g$. Each worker then independently executes $W_p$ steps in an inner loop, where $W_p$ is a large multiple of the number of nodes in subtree $p$ (by default, 50). The intent is to locally equilibrate the subtree conditional on the global parameters and the fixed times and sequences of the boundary nodes between it and its ancestral and descendant subtrees. Each local move is chosen at random from the following weighted list:
      - Inner node displacement move (weight: 15.0).
      - Branch reform move (weight: 15.0)
      - Subtree slide move (weight: 1).
      - Mutation-directed SPR moves (weight: 1; around 1 % of these explore the entire subtree of worker $p$, while the rest explore only the vicinity of $P$ up to 2 mutations away; see Section 6.6.1).
    - The partitions are always reassembled so that at the end of each iteration of the middle loop, where we can sample a complete configuration of the tree and the associated global parameters. We also use this moment to change the reference sequence to match the current root sequence, which requires an expensive walk of the tree to adjust all missations; while not required, this step avoids trapping the MCMC in a state where the root and reference sequences differ a lot, which would make root-changing moves unduly expensive.

We note that our current implementation of the parallelization scheme is merely a proof-of-concept to show that parallelization is possible. There is a lot of needless copying and recalculation when partitioning the tree, distributing the subtrees to workers and then reassembling the full tree from the modified subtrees. Further, while the above scheme in principle supports distributing the work over a cluster of networked machines, our implementation only distributes work among the CPU cores of a single machine. We expect that engineering improvements in both of these areas will unlock scaling to much larger datasets than we have yet demonstrated here.

### 11 Fast MCC trees, approximate incremental MCC trees

Delphy's web interface rederives an MCC every time a new posterior tree is sampled, so it needs to do this quickly. In contrast, BEAST2's TreeAnnotator takes tens of seconds to deduce the MCC of the SARS-CoV-2 results from Lemieux et al [11]. Here, we describe how we derive such MCCs in milliseconds instead, using a technique inspired by hash trees (also known as Merkle trees) [2, 14, 15].

First, recall how MCCs are usually derived. For each posterior tree, we enumerate all its *clades*, i.e., the sets of tips descended from each node. Any particular clade is present in a fraction of the posterior trees, called the clade's *posterior support*. The MCC is defined as a tree with: (a) the topology of the posterior tree with highest product of posterior support over all its nodes; and (b) node times that summarize those of the corresponding nodes in the posterior trees. Most commonly, these latter times are the average (or median) of the times of the inner node for the same clade in each posterior tree where it is present.

What most slows down TreeAnnotator's MCC calculation is its representation of clades as explicit sets of tip indices. This leads to high memory consumption and high runtimes as explicit sets are built, inserted and retrieved from hash tables, and compared to each other. Instead, we introduce a probabilistic approach below that has proven much more effective in practice.

In Delphy's MCC calculation, we represent the clade of a node $i$ as a fixed-size $C$-bit fingerprint $F_i$. We assign completely random fingerprints to tips. The fingerprint of a clade that is the union of two smaller and disjoint clades is calculated as the exclusive-or (XOR) of those two clade's fingerprints. Hence, these fingerprints can be calculated very efficiently and in time linear in the tree size for a single posterior tree, via a post-order traversal of the tree. When this procedure completes, the fingerprint for an arbitrary inner node is the XOR of the fingerprints of all its descendant tips:

$$F_i = \bigotimes_{\text{tips } j \text{ below } i} F_j.$$

If the tip fingerprints are uniformly random, then so are the fingerprints of the inner nodes. Thus, an MCC of $M$ posterior trees, each with $N$ inner nodes, has at most $MN$ distinct clades, whose fingerprints are all essentially independent random $C$-bit fingerprints. We can reduce the probability of a fingerprint collision to an arbitrary level $p$ by using a sufficiently large value of $C$, as follows:

$$p \approx 1 - \exp\left[-\frac{(MN)^2}{2 \cdot 2^C}\right] \approx \frac{(MN)^2}{2 \cdot 2^C}, \qquad \text{(large } C\text{)}.$$

Equivalently,

$$C \approx \log_2\left[\frac{(MN)^2}{2p}\right].$$

Concretely, using $C = 64$ bits for 1000 posterior tree samples, each with 1000 inner nodes, yields $p \approx 3 \times 10^{-8}$. For pandemic scales, e.g., 1000 trees with 10 M inner nodes, one would need $C \gtrsim 104$ to achieve $p \lesssim 10^{-6}$. The natural choice of $C = 128$ bits would lead to $p \approx 1 \times 10^{-19}$. Hence, we see no practical danger in using 64-bit fingerprints for near-real-time analysis of smaller datasets and 128-bit fingerprints for pandemic-scale datasets.

In practice, we build an MCC in four passes. First, we build a map of clade fingerprints to lists of tuples "(posterior tree sample index, node index)". Second, we calculate the log-credibility of each clade $i$

as $\log(M_i/M)$, where $M_i$ counts how many posterior trees contain clade $i$. Third, we compute for each posterior tree the sum of the log-clade-credibilities of each inner node, and record the tree with the highest sum. This tree defines the topology of the MCC. Finally, we iterate over the nodes of the MCC, and for each one, we iterate over the corresponding nodes in the posterior trees to calculate the mean (or median) node time of each MCC node. The node $\phi_m(i)$ in a posterior tree $m$ corresponding to a given node $i$ in the MCC is the MRCA in the posterior tree $m$ of the nodes corresponding to the child nodes of $i$ in the MCC, i.e., $MRCA\{\phi_m(j) | j \text{ child of } i \text{ in MCC}\}$. Hence, node correspondences for a single posterior tree can be calculated efficiently in $\mathcal{O}(N)$ time (in aggregate, assuming all inner node times are unique, the MRCA calculations walk over every branch in the posterior tree exactly once). If all corresponding nodes are included when calculating the mean (median) time, we mimic the behavior of TreeAnnotator's `--heights ca` option; if only exact matches are included, where the clade fingerprint of $\phi_m(i)$ and $i$ are equal, we obtain the more standard MCC. Neglecting the weak $\log N$ dependence of $C$, which we have argued above is immaterial for all practical datasets, this four-pass algorithm runs in time $\mathcal{O}(MN)$, i.e., linear in the size of the results of the MCMC.

#### 11.1 Incremental calculation of MCCs

For continuously visualizing the MCC for an ongoing MCMC run, the above algorithm can still be very expensive, repeats large amounts of work, and slows down as the inference progresses. Concretely, continuously recalculating the MCC as $M$ grows results in an overall unacceptable $\mathcal{O}(M^2N)$ cost. While this has so far proven acceptable for interactive use in the Delphy web interface, we outline a proposal below for incremental MCC calculation that may prove useful for larger datasets and/or longer runs. We initially tested this proposal and verified its effectiveness, but since from-scratch recalculation of MCCs after every sample proved sufficient in the Delphy web interface, we have not implemented this idea in production.

We observe that the MCC derived from a sequence of posterior tree samples $(\mathcal{T}_1, \ldots, \mathcal{T}_M)$ is usually very similar to that derived from a sequence extended with one more sample, $\mathcal{T}_{M+1}$. Most of the time, the topology is unchanged, while the node times make subtle adjustments. But occasionally, the identity of the posterior tree sample with highest clade credibility changes, in which case the tree topology changes drastically.

We exploit the above insight by adapting "beam search" to the problem of incrementally building MCCs. Concretely, when a new posterior tree sample is added, we can calculate its inner node clade fingerprints (incremental pass 1) and update the map of clade fingerprints to pointers to posterior tree nodes (incremental pass 2) in $\mathcal{O}(N)$ time. Then, instead of recalculating the clade credibility of all the $M+1$ posterior tree samples in $\mathcal{O}(MN)$ time, we simply recalculate it for a "beam" of $K$ such trees in $\mathcal{O}(KN)$ time (incremental pass 3), which closely tracks the $K$ trees with highest clade credibilities. We also update the clade credibility of a random posterior tree sample, and if it is higher than the lowest-clade-credibility tree in the beam, it replaces that tree in the beam. We then pick the highest-clade-credibility tree in the beam for the MCC topology. In the likely case that the MCC topology did not change when adding $\mathcal{T}_{M+1}$, then the node times can be incrementally updated in $\mathcal{O}(N)$ time; otherwise, we recalculate them from scratch in $\mathcal{O}(MN)$ time. Overall, then, adding a single posterior tree sample to an incremental MCC takes $\mathcal{O}(N)$ time, and the cost of continuously recalculating the MCC throughout a run is close to $\mathcal{O}(MKN)$ time.

In essence, we obtain an $M/K$ speed-up by trading away some accuracy: the MCC topology might not always correspond to that of the highest-clade-credibility posterior tree sample. This trade-off seems acceptable for the real-time updating of the MCC of an ongoing MCMC run, though it's essential to recalculate the MCC correctly from scratch at the end of a run for further analysis. The parameter $K$ controls the lag between changes in the real MCC topology and that of this incremental MCC. Empirically, we have found that $K = 5$ works sufficiently well to continuously visualize a reasonably accurate MCC during the course of an MCMC run.

#### References

[1] R. Borndörfer, Z. Elijazyfer, and S. Schwartz. "Approximating Balanced Graph Partitions". In: *Zuse Institute Berlin Report* 19.25 (2019).

[2] John W Byers et al. "Informed content delivery across adaptive overlay networks". In: *IEEE/ACM transactions on networking* 12.5 (2004), pp. 767–780.

[3] Alexei J. Drummond et al. "Estimating Mutation Parameters, Population History and Genealogy Simultaneously From Temporally Spaced Sequence Data". In: *Genetics* 161 (2002), pp. 1307–1320.

[4] Joseph Felsenstein. "Evolutionary trees from DNA sequences: A maximum likelihood approach". In: *Journal of Molecular Evolution* 17.6 (1981), pp. 368–376. DOI: `https://doi.org/10.1007/BF01734359`.

[5] W.K. Hastings. "Monte Carlo sampling methods using Markov chains and their applications". In: *Biometrika* 57 (1970), pp. 97–109.

[6] William Howard-Snyder et al. "Densely sampled phylogenies frequently deviate from maximum parsimony in simple and local ways". In: *arXiv* (), 2311.10913 [q–bio.PE]. DOI: `https://doi.org/10.48550/arXiv.2311.10913`.

[7] J.F.C. Kingman. "On the Genealogy of Large Populations". In: *Journal of Applied Probability* 19 (1982), pp. 27–43.

[8] J.F.C. Kingman. "The Coalescent". In: *Stochastic Processes and the Applications* 13 (1982), pp. 235–248.

[9] Alexander M. Kramer et al. "Online phylogenetics with matOptimize produces equivalent trees and is dramatically more efficient for large SARS-CoV-2 phylogenies than de novo and maximum-likelihood implementations". In: *Systematic Biology* 72 (2023), pp. 1039–1051.

[10] Nicolas Lartillot. "Conjugate Gibbs sampling for Bayesian phylogenetic models". In: *Journal of Computational Biology* 13.10 (2006), pp. 1701–1722. DOI: `https://doi.org/10.1089/cmb.2006.13.1701`.

[11] Jacob E. Lemieux et al. "Phylogenetic analysis of SARS-CoV-2 in Boston highlights the impact of superspreading events". In: *Science* 371 (2021), eabe3261. DOI: `https://doi.org/10.1126/science.abe3261`.

[12] Nicola De Maio et al. "Maximum likelihood pandemic-scale phylogenetics". In: *Nature Genetics* 55 (2023), pp. 746–752. DOI: `https://doi.org/10.1038/s41588-023-01368-0`.

[13] Nicola De Maio et al. "Rate variation and recurrent sequence errors in pandemic-scale phylogenetics". In: *bioRxiv* (2024). DOI: `https://doi.org/10.1101/2024.07.12.603240`.

[14] Ralph C Merkle. "A digital signature based on a conventional encryption function". In: *Conference on the theory and application of cryptographic techniques.* Springer. 1987, pp. 369–378.

[15] Ralph C Merkle. *Method of providing digital signatures.* US Patent 4,309,569. 1982.

[16] Nicholas Metropolis et al. "Equations of state calculations byfast computing machines". In: *Journal of Chemical Physics* 21 (1953), pp. 1087–1091.

[17] Vladimir N. Minin, Erik W. Bloomquist, and Mark A. Suchard. "Smooth skyride through a rough skyline: Bayesian coalescent-based inference of population dynamics". In: *Molecular Biology and Evolution* 25 (), pp. 1459–1471. DOI: `https://doi.org/10.1093/molbev/msn090`.

[18] Rasmus Nielsen. "Mapping Mutations on Phylogenies". In: *Systematic Biology* 51.5 (2002), pp. 729–739. DOI: `https://doi.org/10.1080/10635150290102393`.

[19] Rasmus Nielsen. "Mutations as Missing Data: Inferences on the Ages and Distributions of Nonsynonymous and Synonymous Mutations". In: *Genetics* 159.1 (2001), pp. 401–411. DOI: `https://doi.org/10.1093/genetics/159.1.401`.

[20] Áine O'Toole et al. "APOBEC3 deaminase editing in mpox virus as evidence for sustained human transmission since at least 2016". In: *Science* 382 (2023), pp. 595–600. DOI: https://doi.org/10.1126/science.adg8116.

[21] Louis du Plessis et al. "Establishment and lineage dynamics of the SARS-CoV-2 epidemic in the UK". In: *Science* 371 (2021), pp. 708–712.

[22] Ivan Specht, Patrick Varilly, and Pardis C. Sabeti. "JUNIPER: Reconstructing Transmission Events from Next-Generation Sequencing Data at Pandemic Scale". In: *(in preparation)* (2024).

[23] Yatish Turakhia et al. "Ultrafast Sample placement on Existing tRees (UShER) enables real-time phylogenetics for the SARS-CoV-2 pandemic". In: *Nature Genetics* 53.6 (2021), pp. 809–816. DOI: https://doi.org/10.1038/s41588-021-00862-7.

[24] Ian J. Wilson and David J. Balding. "Genealogical Inference From Microsatellite Data". In: *Genetics* 150 (1998), pp. 499–510.

[25] Ziheng Yang. "Estimating patterns of nucleotide substitution". In: *Molecular Evolution* 39 (1994), pp. 105–111.

[26] Ziheng Yang. "Maximum Likelihood Phylogenetic Estimation from DNA Sequences with Variable Rates over Sites: Approximate Methods". In: *Journal of Molecular Evolution* 39 (1994), pp. 306–314.

[27] Cheng Ye et al. "matOptimize: a parallel tree optimization method enables online phylogenetics for SARS-CoV-2". In: *Bioinformatics* 38.15 (2022), pp. 3734–3740. DOI: https://doi.org/10.1093/bioinformatics/btac401.

[28] Chi Zhang, John P. Huelsenbeck, and Fredrik Ronquist. "Using Parsimony-Guided Tree Proposals to Accelerate Convergence in Bayesian Phylogenetic Inference". In: *Systematic Biology* 69.5 (2020), pp. 1016–1032. DOI: https://doi.org/10.1093/sysbio/syaa002.
